# Parallel differentiation extends beyond chromosomal inversions to small genomic islands on a background of rampant gene flow in the European flat oyster (*Ostrea edulis*)

**DOI:** 10.64898/2026.09.28.755087

**Authors:** Lila Colston-Nepali, Paul Doniol-Valcroze, Benjamin Penaud, Céline Reisser, Florence Cornette, Camille Roux, Christelle Fraïsse, Sylvie Lapègue, Nicolas Bierne

**Affiliations:** Biodiversité Marine, Exploitation, et Conservation (UMR MARBEC), Campus Triolet, 093 Place Eugène Bataillon, 34090 Montpellier, France; Institut des Sciences de l’Evolution de Montpellier (ISEM), Campus Triolet, 093 Place Eugène Bataillon, 34090 Montpellier, France; Biodiversité Marine, Exploitation, et Conservation (UMR MARBEC), Station Ifremer, 87 Avenue Jean Monnet, 34200 Sète, France

**Author notes:** CESCO, MNHN, CNRS, Sorbonne Université, Paris, France. co-last authors (Christelle Fraïsse, Sylvie Lapègue, Nicolas Bierne).

## Abstract

Heterogeneous genomic landscapes of differentiation can reveal evolutionary processes acting over different timescales. Whereas genetic structure in the European flat oyster, *Ostrea edulis*, is mainly shaped by geography, previous work suggested parallel differentiation between western and eastern Mediterranean populations and between Atlantic and North Sea populations, resulting in genetic convergence between range margin populations in some genomic regions. Here, we analyzed 27 oysters from four locations using moderate-coverage (∼20X) short-read whole-genome sequencing. We first examined three known chromosomal inversions and found evidence that they predate divergence with the sister species *O. chilensis*. We then scanned the genome for parallel differentiation at the individual-SNP rather than sliding-window scale using the product of allele-frequency differences across the two geographic contrasts. Two of the three inversions were detected, but we also identified 12 small non-inverted genomic regions showing even stronger signals of parallel differentiation. These “islands of parallelism” typically overlap one or a few genes, and most showed elevated d_XY_ consistent with long-term barriers to gene flow. Two islands overlap nuclear ATP synthase subunits, while mitochondrial ATP8 shows a convergent amino-acid change, suggesting possible nuclear–nuclear and mitonuclear incompatibilities. Finally, demographic inference with heterogeneous gene flow supported post-glacial secondary contact and a lineage-based rather than geography-based population topology. Our results suggest that the genomes of *O. edulis* lineages have been largely porous to gene flow upon secondary contact, except for chromosomal inversions and small genomic islands. These regions likely act as barrier loci, preserving a signal of a shared evolutionary history among marginal populations that has otherwise been erased by rampant gene flow.

## INTRODUCTION

Evolutionary processes do not impact the genome in a homogenous manner, and investigating the patterns of divergence across the genome allows us to address fundamental questions concerning evolution and speciation (Wolf and Ellegren 2017). A heterogenous landscape of genetic differentiation may reflect evolution acting upon multiple timescales, with different regions of the genome reflecting different processes, from genetic drift and demographic history to gene flow and linked selection (Bierne, Gagnaire and David 2013; Van Belleghem et al. 2018; Ravinet et al. 2017; Gwee et al. 2025). Certain genomic islands of differentiation within the same species may indeed represent different processes: between the Mediterranean and Atlantic lineages of the European sea bass (*Dicentrarchus labrax*), some islands contain barrier loci where divergence (d_XY_) has accumulated between lineages over time, whereas others result from linked-selection (hitchhiking and background selection) within lineages (Duranton et al. 2018). Determining which processes contributed to these islands is difficult, as very different processes can generate similar genomic landscapes (Ravinet et al. 2017). Investigating population replicates of divergence might therefore be an important tool for better understanding these complex genomic landscapes (Wolf and Ellegren 2017).

Parallel evolution, whereby evolutionary change proceeds in the same direction across comparable evolutionary contrasts, provides natural replicates that can reveal fundamental patterns of adaptive evolution (Elmer and Meyer 2011), and help investigate the predictability of evolution and its underlying constraints (Conte et al. 2012; Faria et al. 2014; Nosil et al. 2024). Parallel patterns of genomic differentiation provide a useful framework for disentangling the effects of demographic history from those of selection. As opposed to convergent evolution, parallel genetic differentiation acts on closely related populations (Futuyma 1986; Elmer and Meyer 2011; Cerca 2023). Therefore, though it may occur due to *de novo* mutations, it is more likely to act upon alleles that are shared between populations, either through gene flow, or incomplete sorting of ancestral polymorphisms (see Faria et al. 2014, and Welch and Jiggins 2014). Furthermore, when selection acts in directions independent of, or opposed to, geographic structuring, genomic scans of parallel differentiation have a much lower rate of false positives than when selection creates differentiation aligned with isolation by distance (Bradburd, Ralph and Coop 2013; Fraïsse et al. 2014; Lotterhos and Whitlock 2015; Riquet et al. 2019).

Chromosomal inversions (hereafter: inversions) often contribute to both a heterogeneous genomic landscape, and parallel evolution (Westram et al. 2022). Recombination is suppressed at inversions between heterokaryotypes, which maintain regions of divergence despite gene flow (Akopyan et al. 2022), and may facilitate local adaptation (e.g. prairie sunflowers *Helianthus petiolaris*, Huang et al. 2020; rainbow trout *Oncorhynchus mykiss*, Arostegui et al. 2019; honeybees *Apis mellifera*, Christmas et al. 2019; deer mice *Peromyscus maniculatus*, Harringmeyer and Hoekstra 2022; long-snouted seahorses, *Hippocampus guttulatus*, Meyer et al. 2024; and *Littorina* snails, Le Moan et al. 2024). Inversions may therefore fuel genetic parallelism, by allowing the simultaneous transfer of multiple genes, functioning as ‘adaptive cassettes’ via gene flow (Morales et al. 2019). Alternatively, inverted alleles can be maintained over very large time-scales (Wellenreuther and Bernatchez 2018), thereby preserving shared ancestral variation that traces back to ancient episodes of introgression or common evolutionary history. Simulations have demonstrated that inversions can result in stronger molecular signals of parallel evolution and faster adaptation (Westram et al. 2022), and empirical studies have found a disproportionate contribution of inversions or large haploblocks to parallel differentiation (Vangestel et al. 2024; Battlay et al. 2025a; Battlay et al. 2025b). However, these studies share a limitation inherent to sliding-window approaches. Averaging differentiation across neighboring SNPs readily captures broad differentiated regions, such as chromosomal inversions, but can dilute narrow signals restricted to only a few SNPs. This problem has motivated the use of statistics based on the upper tail or maximum of SNP differentiation within windows (Fraïsse et al. 2016; da Silva Ribeiro et al. 2022). Thus, conclusions that differentiation is disproportionately concentrated within inversions may need to be revisited in light of this caveat.

Finally, in the context of parallel evolution, it is difficult, yet fundamental, to distinguish between two alternative scenarios that can generate similar genomic patterns, as outlined by Gwee et al. (2025): (i) genome-wide swamping (hereafter the “swamping hypothesis”), in which barrier loci remain associated with the original lineages while secondary gene flow homogenizes the rest of the genome (the genomic islands stay while the genomic background moves), and (ii) locus-specific introgression (hereafter the “introgression hypothesis”), in which the genomic background retains the original population history while locally favoured alleles move between populations (the genome stays while the genomic islands move). The latter corresponds to the “transporter hypothesis” of Schluter and Conte (2009), whereby adaptive alleles can spread between similar selective environments through gene flow across intervening populations where they are counter-selected, a property that distinguishes locally adaptive alleles from alleles involved in intrinsic genetic incompatibilities (Bierne, Gagnaire and David 2013).

The European flat oyster (*Ostrea edulis*) is an economically important species with a significantly reduced distribution due to over exploitation and disease (Pouvreau et al. 2023), and is currently the focus of restoration efforts (Pogoda et al. 2019). Lapègue et al. (2023) characterized its genetic structure across its range using 203 SNPs and identified four genetic clusters: Atlantic and North Sea clusters in the northern European seas, and western and eastern Mediterranean clusters in the southern European seas. Most of the genome followed geographic population structure, while two groups of linked SNPs, inferred to belong to chromosomal rearrangements, showed similar allele frequencies between the North Sea and eastern Mediterranean populations. These similarities in allele frequencies suggested the presence of two distinct genetic forms, although it remained unclear how much of the genome differentiated them and how long they had been diverging, leaving their status somewhere between historically diverged lineages and chromosomal forms defined by inversions. One form grouped the Atlantic and western Mediterranean populations, and the other form grouped the North Sea and eastern Mediterranean populations at the two edges of their distribution range. Whether the occurrence of these lineages is driven by independent selection regimes (“introgression hypothesis”), or by an episode of shared evolutionary history (“swamping hypothesis”) is unclear, and requires higher depth sequencing and demographic reconstruction.

Three chromosomal inversions were subsequently confirmed using low-coverage whole-genome sequencing (Alves Monteiro et al. 2024; though a fourth may also exist, see Ewers et al. 2025). Contrasting inversion allele frequencies between the North Sea and the rest of the range further supported the hypothesis of two chromosomal forms (Alves Monteiro et al. 2024; Sambade et al. 2025). However, because the eastern Mediterranean was not sampled, whether the North Sea chromosomal form also occurs there, as suggested by Lapègue et al. (2023), remained to be confirmed. In order to better understand these contrasts in genetic clustering along the *O. edulis* genome, 27 individuals were resequenced using ∼20X coverage short-read whole genome sequencing, representing the four genetic clusters identified by Lapègue et al. (2023). In this paper, we explore the genomic landscape of differentiation in the flat oyster and i) confirm the presence of the same chromosomal form at the two edges of the range, ii) describe additional genomic regions showing parallel allele frequency shifts between the two forms across both geographic regions, and iii) reconstruct the demographic history of the species to better understand the evolutionary processes underlying the parallel signals observed (“swamping” vs “introgression” hypotheses). Our sequencing coverage allowed stringent filtering of paralogous SNPs, which are frequent in this species (Colston-Nepali, Lapègue and Bierne 2026), and provided a dataset suitable for demographic inference.

## METHODS

### Sampling and sequencing

*O. edulis* samples were collected from four different locations: the North Sea, the Atlantic, the western Mediterranean Sea, and the Black Sea (Figure 1A, Supplementary Table 1). These locations were selected to represent the four genetic clusters identified by Lapègue et al. (2023), and also represent two regions: the Northern European Seas (including the samples from the Atlantic and the North Sea, hereafter: NES), and the Southern European Seas (including the samples from the western Mediterranean and Black Sea, hereafter: SES). These locations also represent the two putative lineages suggested by Lapègue et al. (2023), where Lineage 1 includes the North Sea and Black Sea, and Lineage 2 includes the Atlantic and western Mediterranean (Figure 1).

**Figure 1.**
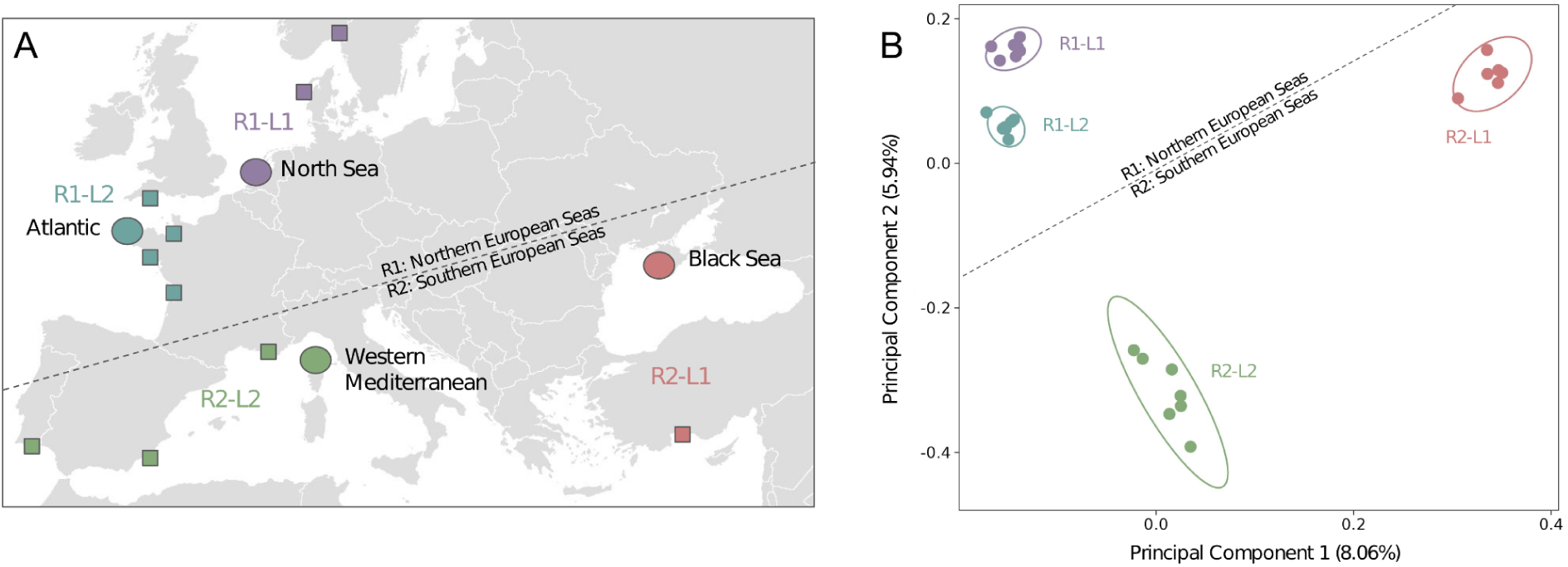
**A** Map of sampling locations. Circles indicate the sampling locations used in this project. Squares represent the additional sampling locations included by Lapègue et al. (2023). Colours indicate the genetic clusters assigned by Lapègue et al. (2023). Dashed line indicating proposed regional split between Northern European Seas and Southern European Seas. Additional labels indicating proposed regional assignments (R1, R2), and proposed lineage assignments (L1, L2). Note that one North Sea sample in our study originates from Denmark. **B** Corresponding principal components analysis of the using 423,929 SNPs. Population indicated by colour (Blue: Atlantic, Purple: North Sea, Green: western Mediterranean, Red: Black Sea). Dashed line indicating proposed regional split between Northern European Seas and Southern European Seas. Additional labels indicating proposed regional assignments (R1, R2), and proposed lineage assignments (L1, L2).

Seven individuals were sampled from all sites, except the Black Sea with six individuals (totalling 27 samples). A last sample of the dataset was a technical replicate used to adjust our filtering parameters, and was subsequently discarded from the analysis.

In order to have an outgroup species for demographic history inference, we also sequenced one individual from the species *O. chilensis*. This individual was sampled in the Menai Strait, Wales (see Gayral et al. 2013 for sampling), and sequenced together with the 27 (+1 replicate) *O. edulis* samples. DNA preparation, sequencing, read processing, variant calling, and filtering are described in detail in the Supplementary Methods.

### Chromosomal inversions

Pairwise genome comparisons undertaken by the program D-GENIES (Cabanettes and Klopp 2018), and multi-dimensional scaling (MDS) using the R package lostruct on the genome level (R Core Team 2024, Li and Ralph 2019) were used to verify the presence and location of large chromosomal inversions. Inversions were then more closely investigated using a local principal components analysis using lostruct on the chromosome level. Local PCAs were conducted on 5kb SNP windows.

Genomewide pairwise F_ST_ for all population combinations was calculated using the program pixy (Korunes and Samuk 2021). To further investigate the inversions, pixy was also used to calculate pairwise d_XY_ (a measure of genetic divergence) between inversion alleles, and pi (a measure of genetic diversity) for each inversion allele. To explore the relationship between the North Sea and the Black Sea, the pi for each population at the inversion region on chromosome 9 was also calculated. Finally, an identity by state (IBS) analysis was performed with the R package SNPRelate (R Core Team 2024, Zheng et al. 2012).

The program SnpEff (Cingolani et al. 2012) was used to annotate the SNPs found in inverted regions to determine whether these regions are enriched for particular functional genes. A new database for *O. edulis* was built using annotation and coding sequence (CDS) for the assembly, and separate annotations created for each inverted region. The program Seqtk (https://github.com/lh3/seqtk) was used to convert the genelists output by SnpEff into fasta files. These fasta files were used as input for the KEGG Orthology-Based Annotation System (KOBAS, Bu et al. 2021) in order to perform gene ontology (GO) enrichment analysis. The most closely related species in the KOBAS database was *Magallana gigas* (previously known as *Crassostrea gigas*), which was used as the reference species. Enrichment analysis was run using Fisher’s exact test, with Benjamini and Hochberg (1995) correction for false discovery.

To investigate whether inversions have a higher genetic load than collinear (non-inverted) regions, SnpEff was also run on the collinear regions of chromosomes 4, 5, and 9. Centromeric regions were estimated from pixy results (identified by valleys in estimates of d_XY_ and pi), and removed from analysis. SnpEff provides counts of variants in several categories of impact. ‘Low’ includes low impact changes such as synonymous mutations, ‘Moderate’ includes moderate impact changes such as insertion of a codon, and ‘High’ includes high impact changes such as deletion of a gene. Counts for each of these categories were compared for collinear regions (for chromosomes 4, 5, and 9 combined), and each inversion region.

### Signals of parallelism

To further investigate parallel allele frequency shifts between putative lineages (L1 vs. L2) across geographic regions (R1 and R2), a single statistic was defined. The product of the allele frequency differential, Π= Δ_R1_. Δ_R2_ = (*p*_R1L1_ - *p*_R1L2_). (*p*_R2L1_ - *p*_R2L2_), is expected to provide a positive value when the signal is parallel (the same allele is at high or at low frequency in both L1 populations) and a negative value if it is antiparallel (the same allele is at high or at low frequency in one of the L1 populations, with the opposite frequency in the other).

Π is expected to reach high outlier values when the allele frequency differential is high in both regions. This event would be very difficult to obtain without selection, as the locus would need to be a false positive two times, and against the genomewide trend of geographic structure (Fraïsse et al. 2014). Such a product of allele frequency differential is commonly used to identify gene flow and is known as the f4 statistic (Patterson 2012). Given we use such a product of allele frequency differential for another purpose, and in another population configuration, we preferred to give it a different name, Π. The program BCFtools (Danecek et al. 2021) v.0.1.16. was used to calculate the allele frequencies for each SNP per population and the Π score was calculated across the genome with the following formula, using a custom script in R:

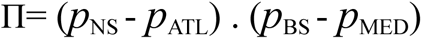

Where *p*_NS_ is the allele frequency in the North Sea population (representing R1L2), *p*_ATL_ is the allele frequency in the Atlantic population (representing R1L1), *p*_BS_ is the allele frequency in the Black Sea population (representing R2L2), and *p*_MED_ is the allele frequency in the western Mediterranean population (representing R2L1). SNPs with a strong positive Π score (0.5 or higher) were classified as parallel outliers, and SNPs with a strong negative Π score (−0.5 or lower) were classified as antiparallel outliers.

To further investigate parallel outliers, SNPs were annotated using SNPeff (Cingolani et al. 2012), and annotation categories simplified into four major categories: intronic, intergenic, synonymous, and nonsynonymous (see Supplementary Table 2 for details). The NCBI Basic Local Genome Alignment Tool (BLAST, Altschul et al. 1990) and the NCBI Genome Data Viewer were used to identify genes in the parallel regions.

Finally, mitochondrial genomes were assembled for all 27 individuals, and the outgroup sample using the program MitoZ (Meng et al. 2019). Unexpectedly, mitochondrial coverage was only 0.5-2 times the nuclear coverage across individuals, possibly reflecting a low mitochondrial content in the sampled tissue. Consequently, three individuals (one each from the Atlantic, Mediterranean, and Black Sea) did not yield complete mitochondrial genomes and were excluded from the analysis. A mitochondrial maximum-likelihood (ML) tree was constructed from the concatenated coding sequences of the 13 protein-coding genes using the General Time Reversible (GTR) model implemented in MEGA (Kumar et al. 2024). We mapped amino-acid changes in the mitochondrial ATP6 and ATP8 subunits onto the tree using parsimony, although amino-acid variation was only found in ATP8.

### Demographic history analyses

A modified version of the Python program vcf2fasta (https://github.com/santiagosnchez/vcf2fasta, see Code Repository) was run in order to produce the input fasta files needed for the demographic inferences. The option ‘gene’ was selected in vcf2fasta in order to include UTRs, introns and exons in the output. A custom python script was then used to insert headers into the file with population information.

#### Two-population models

To infer the demography of pairwise *O. edulis* populations, the Demographic Inferences with Linked Selection using ABC (DILS) software was used (Fraïsse et al. 2021, available at: https://github.com/popgenomics/DILS). The first step of DILS was run on the 23,615 genes to ascertain baseline statistics, and to test out filtering parameters within the program. We set a minimum sequence length of 500bp, a maximum fraction of 40% missing data in a sequence, and a minimum number of sequences per locus to 7 or 8 (depending on the pairwise comparison used, as the Black Sea population has one less individual than the other populations), resulting in ∼16,660 genes retained by the program.

For the full runs of the program, computational efficiency needed to be maximised so genes were filtered to a maximum of 5,000bp. One gene from each of the 12 parallel islands and each inversion were added to the input fasta file, and all others from the inversion regions excluded.

This was done to retain the signal of differentiation within each inversion while excluding inversions themselves whose specific features are not well accommodated by the inference model. A total of 3,166-3,168 genes were used by the program depending on the population comparisons. For each comparison, the program was run six times. Final configuration files for each comparison are included in the Demographic Inference Appendix.

We used models of two-population divergence with constant population size. Different timing of gene flow were tested, with two models assuming the absence of current gene flow (“strict isolation” and “ancient migration”), and two models assuming current gene flow (“isolation with continuous migration” and “secondary contact”). We also tested the effect of barriers to gene flow on genetic patterns modelling a beta distribution for the effective migration rate across the genome (modeBarrier: “beta”). We used a generation time of two years and a mutation rate of 2.76e-8 mutations/site/generation (Roux et al. 2014). *O. chilensis* was used as an outgroup to calibrate the per locus mutation rate, and to unfold the site frequency spectrum.

#### Four-population models

To jointly infer the evolutionary history of all four *O. edulis* populations, i.e. between both regions (Northern European Seas vs. Southern European Seas), and between populations in each lineage (the North Sea vs. the Black Sea and the Atlantic vs. the western Mediterranean), we next ran the software DILS_4pop (Postel et al. 2025, updated version available at: https://doi.org/10.5281/zenodo.22829689). The inference results from pairwise comparisons were used to constrain the parameter space of the four-population inference, which involves 54 parameters. Specifically, we used narrower prior bounds (drawn from uniform distributions) for the population sizes with *N* ∼ U(1,000-50,000) instead of U(100-100,000), as well as for the times with *T* ∼ U(1,000-200,000) instead of U(100-250,000). Prior bounds for the migration rates were the same in the pairwise and four-population inferences with *M* ∼ U(0.4-100). A total of 3620 genes were used in the four-population inference. The final configuration file is included in the Demographic Inference Appendix.

We statistically compared three topologies, and estimated the parameters of the best-supported topology: i) “geography” (topo1: (A,((B,C),D))), ii) “mitochondrial tree” (topo2: (A,(B,(C, D)))) and iii) “genetic lineage” (topo3: ((A, B),(C, D))), where A is the Black Sea, B is the North Sea, C is the Atlantic and D is the western Mediterranean. At each population split, the new ancestral population (backward in times) had an independent size from the daughter populations. The four current lineages had the possibility to exchange genes during six independent secondary contacts, i.e. migration rates were estimated between all possible pairs. As in the two-population models, the effect of barriers to gene flow on genetic patterns were accounted for by modelling a beta distribution for the effective migration rate across the genome. We applied the same generation time and mutation rate as in the pairwise inferences, however no outgroup was used.

## RESULTS

### Population structure validation

Before investigating chromosomal inversions and signals of parallel differentiation, we first verified that the genome-wide population structure of our samples was consistent with that previously inferred from a smaller set of markers by Lapègue et al. (2023). Population structure was assessed using a principal component analysis, which recovered the four geographic clusters corresponding to the North Sea, Atlantic, western Mediterranean and Black Sea populations (Figure 1B), providing the population framework used throughout the subsequent analyses. Full details of this preliminary analysis is provided in the Supplementary Analyses.

### Chromosomal inversions

D-GENIES analyses identified chromosomal inversions at the end of chromosomes 4 and 9, and at the beginning of chromosome 5 (Supplementary Figure 1). This was corroborated by the genome-wide MDS analyses (Supplementary Figure 2). It has to be noted that chromosome numbering differs between assemblies, and these numbers refer to the assembly by Li et al. (2023). Chromosome 9 here corresponds to chromosome 8 on both the Roscoff and Roslin assembly, so previous work has identified the largest inversion as occurring there. (See Supplementary Table 3 for chromosome numbering on different genome assemblies).

A geographic pattern was observed within each chromosomal inversion, differing between the three (Fig. 2). The individuals from the western Mediterranean are fixed for one inversion allele on chromosome 4 (Allele A, Fig. 2A, Supplementary Figure 3). The individuals from the Black Sea are either homozygous for the alternative inversion allele (Allele B), or are heterozygous. The Atlantic and North Sea have individuals which carry all three inversion genotypes. For the inversion on chromosome 5, the Atlantic is fixed for one inversion allele (Allele A), and the Black Sea for the other (Allele B, Fig. 2B, Supplementary Figure 4). The samples from the North Sea and western Mediterranean are either homozygous for the same allele as the Atlantic (Allele A), or are heterozygous. Finally, the North Sea and Black Sea are fixed for the same inversion allele on chromosome 9 (Allele B, Fig. 2C, Supplementary Figure 5). Two western Mediterranean individuals also carry this genotype. Two Atlantic individuals are homozygous for the other inversion allele (Allele A), and the remaining western Mediterranean and Atlantic samples are heterozygous.

**Figure 2.**
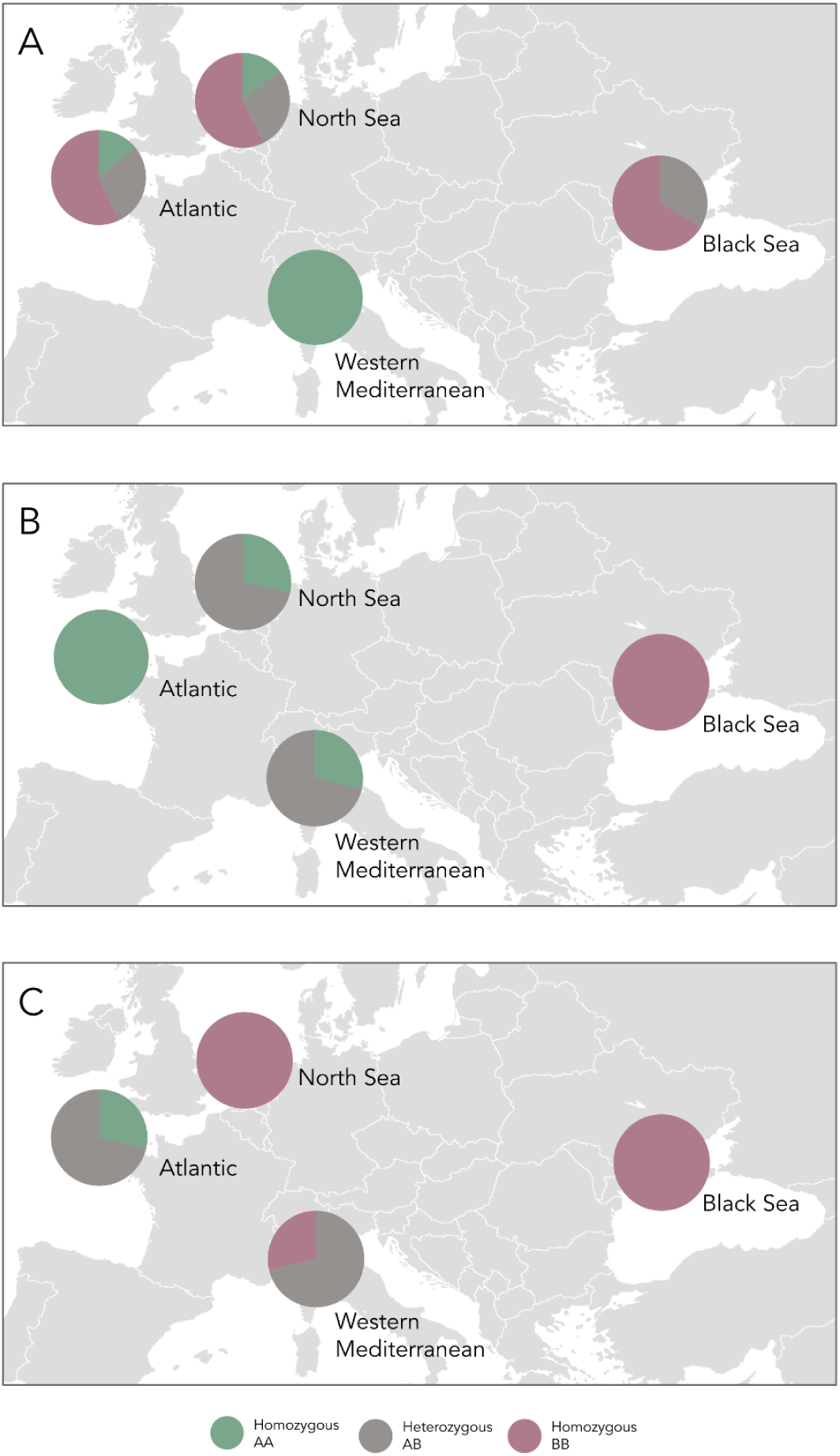
Frequency of inversion homokaryotypes and heterokaryotypes for the inversions present on **A** chromosome 4, **B** chromosome 5, and **C** chromosome 9. Note that A and B inversion alleles are arbitrarily assigned.

Genomewide population-level pairwise F_ST_ scans corroborated these results, with elevated blocks of F_ST_ present at inversion regions between certain population pairs (see Supplementary Figures 6-11). pixy scans also demonstrated elevated F_ST_ and d_XY_ between inversion alleles at the inversion region (Supplementary Figures 12-14).

The average pi for each inversion region did not differ greatly between inversion alleles on any of the three chromosomes (Fig. 3), indicating a very similar amount of genetic diversity carried in each inversion allele. This similarity suggests that the inversions are old, and enough time has passed for the inverted allele to accrue diversity. We note, however, that two of the inversions have a very small inter-allele difference: for the inversion on chromosome 4, one inversion allele (allele A, carried as a homozygous genotype by all individuals from the western Mediterranean, and one individual each from the Atlantic and North Sea) had a slightly higher average pi than the other allele, (allele B, carried as a homozygous genotype by several individuals from the Atlantic, North Sea, and Black Sea) (Fig. 3A). Similarly in chromosome 5, the average pi of one of the inversion alleles (allele A, carried as a homozygous genotype by all individuals from the Atlantic, and some individuals from the western Mediterranean and North Sea) was higher than the other (allele B, carried as a homozygous genotype by all individuals from the Black Sea) (Fig. 3B). The lower pi in these alleles may indicate that they are the derived inversion allele, but alternatively may be the result of sampling bias, or a difference in the effective sizes of the two allele populations. The average pi is very similar between the inversion alleles for the inversion region on chromosome 9 (Fig. 3C). The average pi was also very similar between the North Sea and Black Sea for the inversion region on chromosome 9 (Fig. 3D), suggesting that both populations have had the allele for a long time.

**Figure 3.**
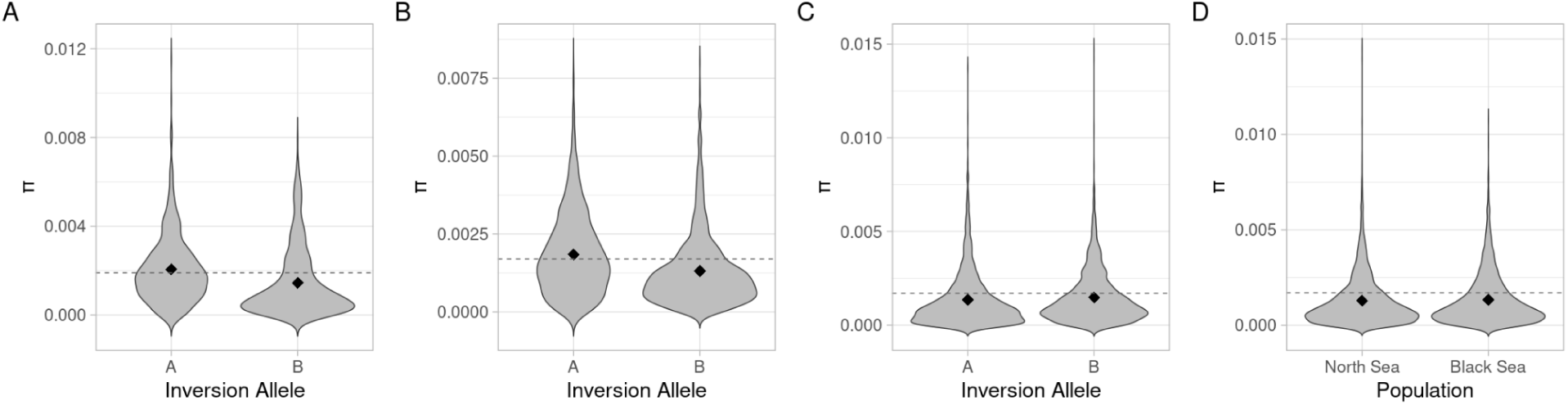
Average pi for each inversion allele in the inversion region on: **A** chromosome 4, **B** chromosome 5, **C** chromosome 9. **D** Average pi for the North Sea and Black Sea in the inversion region on chromosome 9. The dotted line indicates the chromosomal average for pi. (Note that A and B inversion alleles match those of Fig. 2, which were arbitrarily assigned.)

Finally, a neighbor-joining tree on the nucleotide differences (i.e. IBS) displayed the outgroup species (*O. chilensis*) as more closely related to one of the inversion alleles for each inversion (Fig. 4).

**Figure 4.**
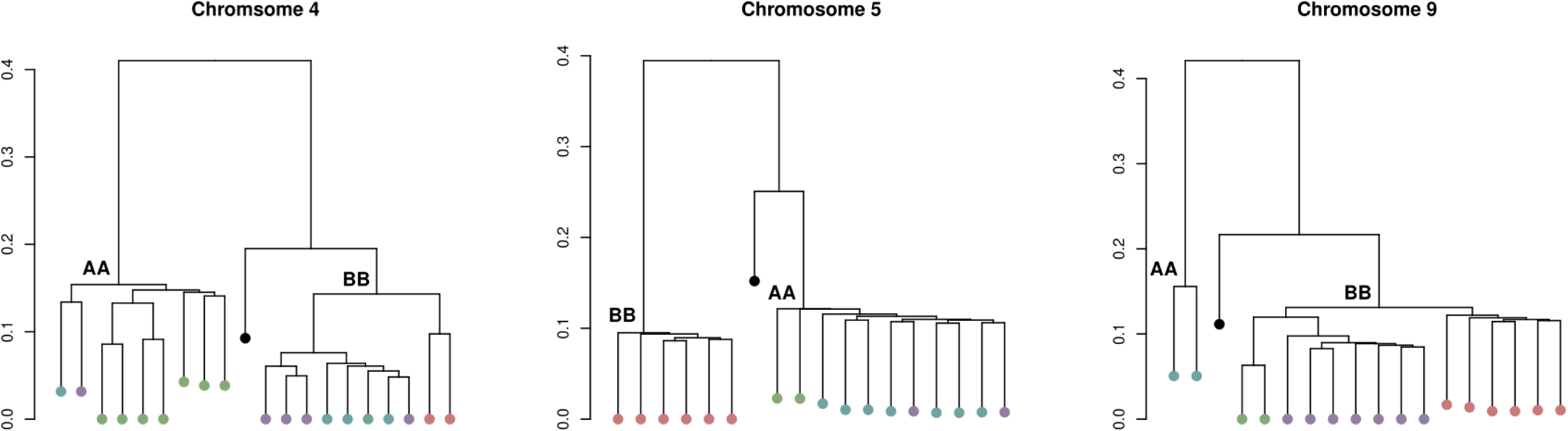
Identical-by-state (IBS) neighbor-joining trees for the three chromosomal inversions in *O. edulis* demonstrating the grouping of the outgroup species with inversion alleles on chromosomes 4, 5, and 9. Population indicated by colour (Blue: Atlantic, Purple: North Sea, Green: western Mediterranean, Red: Black Sea, Black: Outgroup species, *O. chilensis*). Only individuals with homozygous inversion genotypes were included. Homokaryotype indicated as AA or BB (corresponding with Figs. 2 & 3, and arbitrarily assigned).

Inversions had varying levels of gene enrichment (see Supplementary Tables 4-6). No GO terms were found to be significantly enriched in inversion 4 after correcting for false discovery. Two GO terms were significantly enriched in the inversion on chromosome 9: pentose and glucuronate interconversions, and metabolic pathways. The inversion on chromosome 5 demonstrated the most enrichment, with seven GO terms significantly enriched, mainly concerning metabolism.

In all three inversion regions, and the non-inversion region, the majority of SNPs were classified as low impact (59.1-62.3%), and very few SNPs were classified as high impact (0.44-0.83%). However, the inversion regions on chromosomes 4 and 5 had a small, but significant (Fig. 5, and see Supplementary Statistics) difference in proportion of classified variants from the non-inversion region, with slightly less moderate and high impact SNPs, and slightly more low impact SNPs. The proportion of low vs. moderate vs. high impact variants in the inversion region on chromosome 9 was not statistically different from that of the non-inversion region.

**Figure 5.**
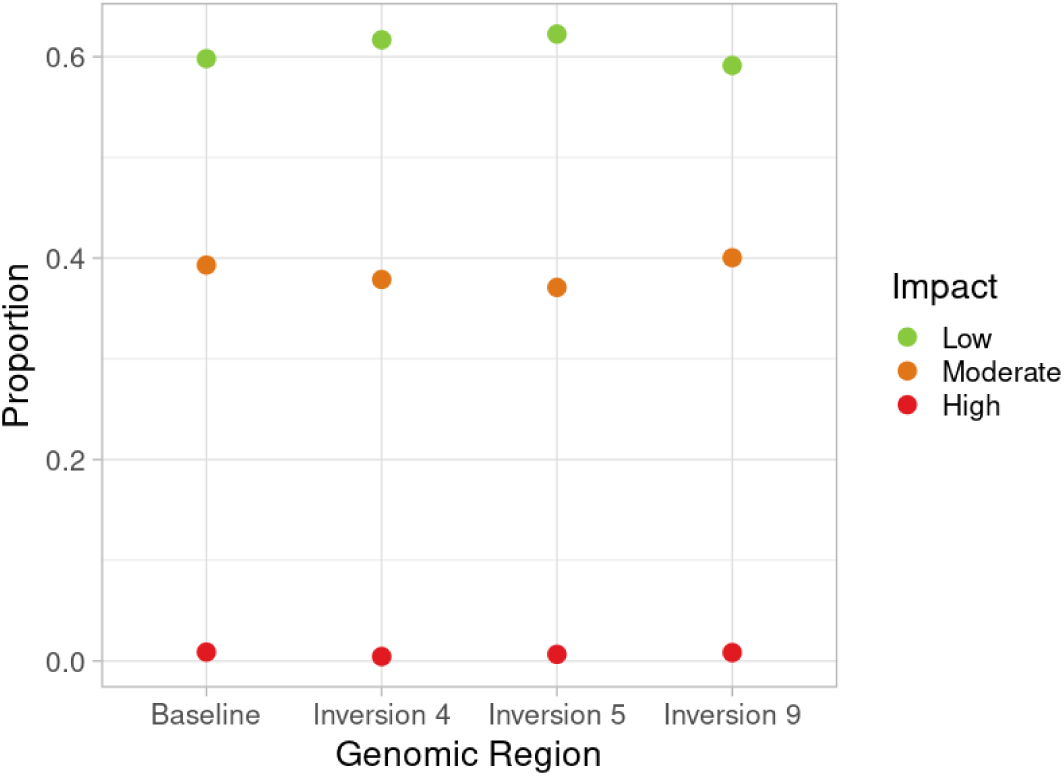
Proportion of variants classified as having low, moderate or high impact in each inversion, and in the baseline regions of these chromosomes. Baseline includes the variants outside of inversion regions on chromosomes 4, 5, and 9, but excluding centromeric regions.

### Signals of parallelism

Across the genome, 2,236 SNPs were classified as having a parallel signal, with just 64 SNPs classified as having an antiparallel signal (Fig. 6). A strong parallel signal can be seen in the inversion region of chromosome 9. A signal of parallelism can also be seen for the inversion region of chromosome 5, with a weaker signal for the inversion region on chromosome 4. However, the strongest parallel signals are found in the collinear regions, in small concentrated ‘islands’ throughout the genome. Islands were identified when a minimum of 3 SNPs in close proximity (within < 25kb) demonstrated a parallel signal. Twelve islands were identified. A single non-synonymous site was also identified with a strong parallel signal (see marking G on Fig. 6).

**Figure 6.**
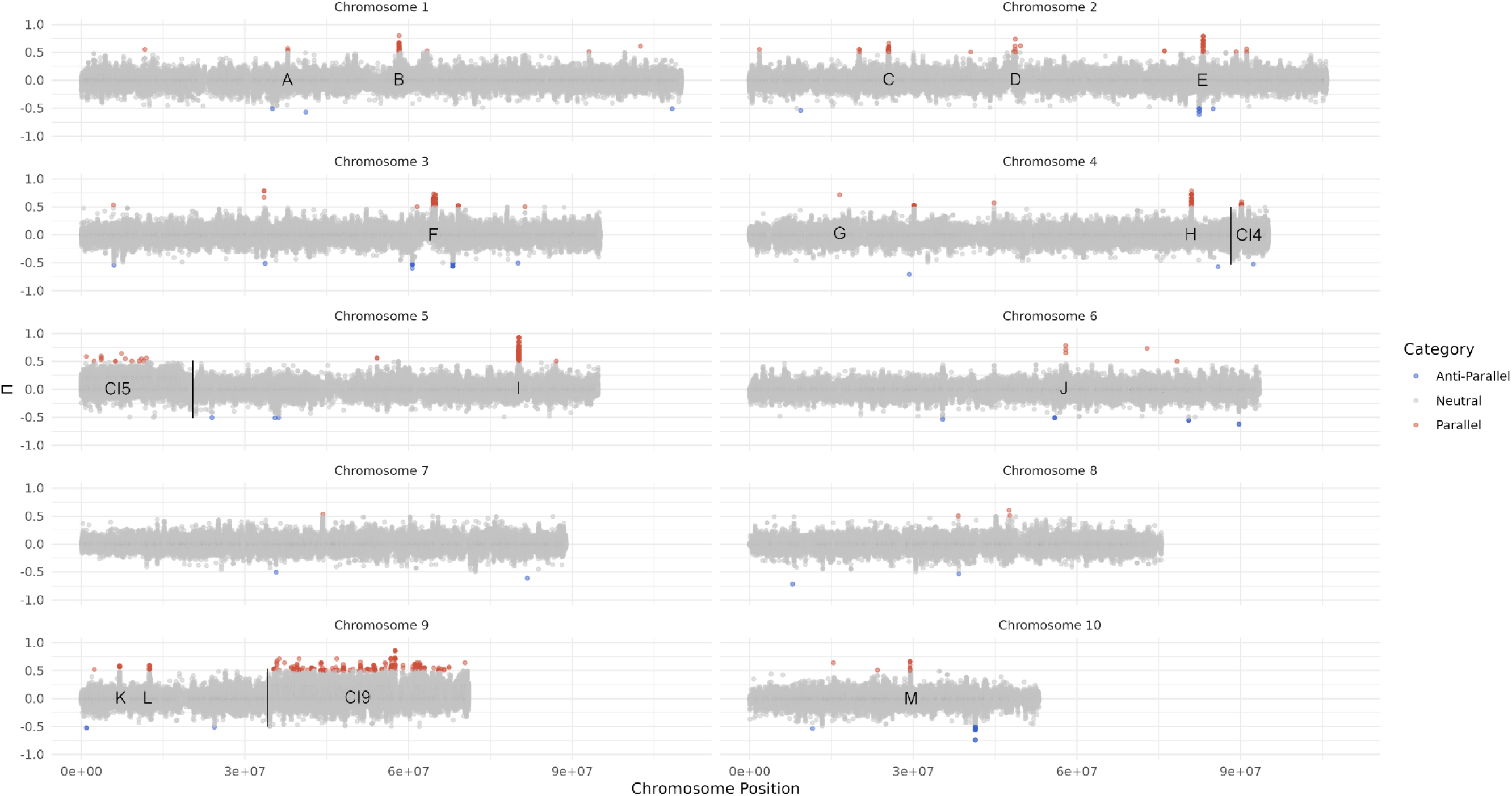
Per SNP Π score, demonstrating the signal of genetic parallelism across all 10 chromosomes. SNPs are classified as parallel (red), non-parallel (grey), or antiparallel (blue). Letters (A-M) indicate parallel regions of interest: **A**- Island 1, **B**- Island 2, **C**- Island 3, **D**- Island 4, **E**- Island 5, **F**- Island 6, **G**- Single non-synonymous site, **H**- Island 7, **I**- Island 8, **J**- Island 9, **K**- Island 10, **L**- Island 11, **M**- Island 12 (see also Supplementary Table 7). Horizontal black lines indicate estimated breakpoint of inversions. Inversions marked for the inversion on **CI4**- chromosome 4, **CI5** - chromosome 5, and **CI9**- chromosome 9.

For each of these 13 parallel regions of interest (Fig. 6 and Supplementary Table 7), the SNP with the highest Π score was identified. Allele frequencies are strongly correlated within genetic lineages, with similar allele frequencies observed between the North Sea and the Black Sea, and between the Atlantic and Mediterranean (Fig. 7A-M). Allele frequencies showed strong differences within geographic regions (i.e. the Northern European Seas and the Southern European Seas). However, in 11 of the 13 SNPs, the difference between allele frequencies in populations was higher in the SES than the NES. In the remaining two SNPs, allele frequency differences were the same between regions (Fig. 7D) or higher in the NES (Fig. 7M). The inversion allele frequencies (Figures 7CI4, 7CI5, and 7CI9) showed much weaker correlations within lineages and differences within regions (except for the differences between populations in the SES region for the inversion on chromosome 4, Fig. 7CI4).

**Figure 7.**
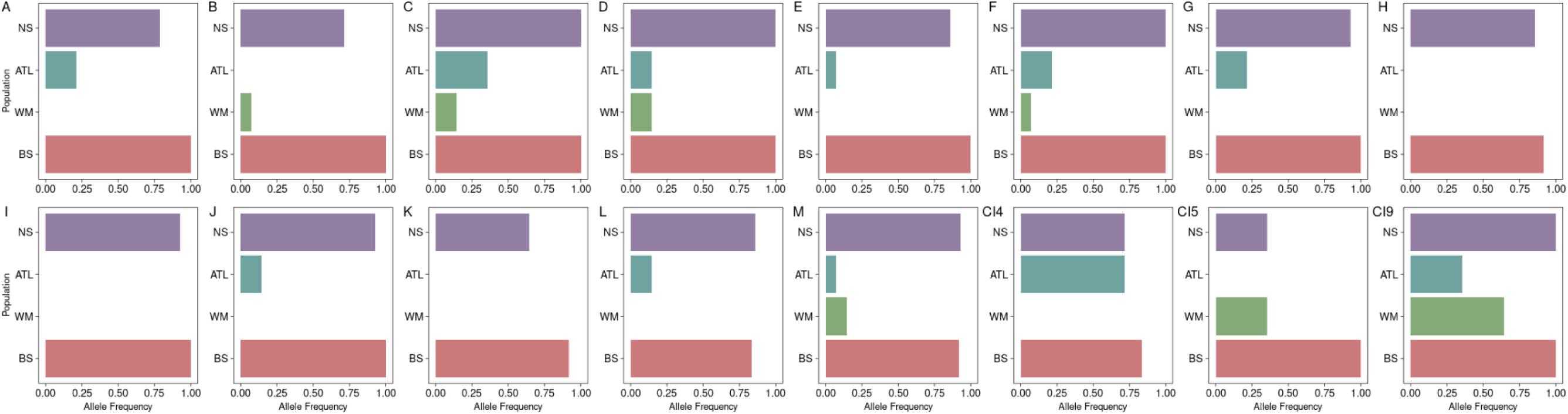
Allele frequencies of each of the four populations for the SNP demonstrating the highest Π score at parallel regions of interest: **A**- Island 1, **B**- Island 2, **C**- Island 3, **D**- Island 4, **E**- Island 5, **F**- Island 6, **G**- Single non-synonymous site, **H**- Island 7, **I**- Island 8, **J**- Island 9, **K**- Island 10, **L**- Island 11, **M**- Island 12. Inversion allele frequencies for each of the four populations for the inversion on: **CI4**- chromosome 4, **CI5** - chromosome 5, **CI9**- chromosome 9. Labels correspond to marked regions of interest on the Π scan (Fig. 6). (NS: North Sea, ATL: Atlantic, WM: western Mediterranean, BS: Black Sea).

The annotation categories for the islands and surrounding sites was plotted, alongside the corresponding scores for pi for all four populations, and pairwise d_XY_ and F_ST_ calculated for the populations in the NES and SES (Figures 8-9 and Supplementary Figures 15-24). Other SNPs that demonstrated parallel signals, but were single outliers, were not further investigated, except for one at the beginning of chromosome 4 that was classified as a nonsynonymous SNP (Fig. 7G, Supplementary Figure 25).

**Figure 8.**
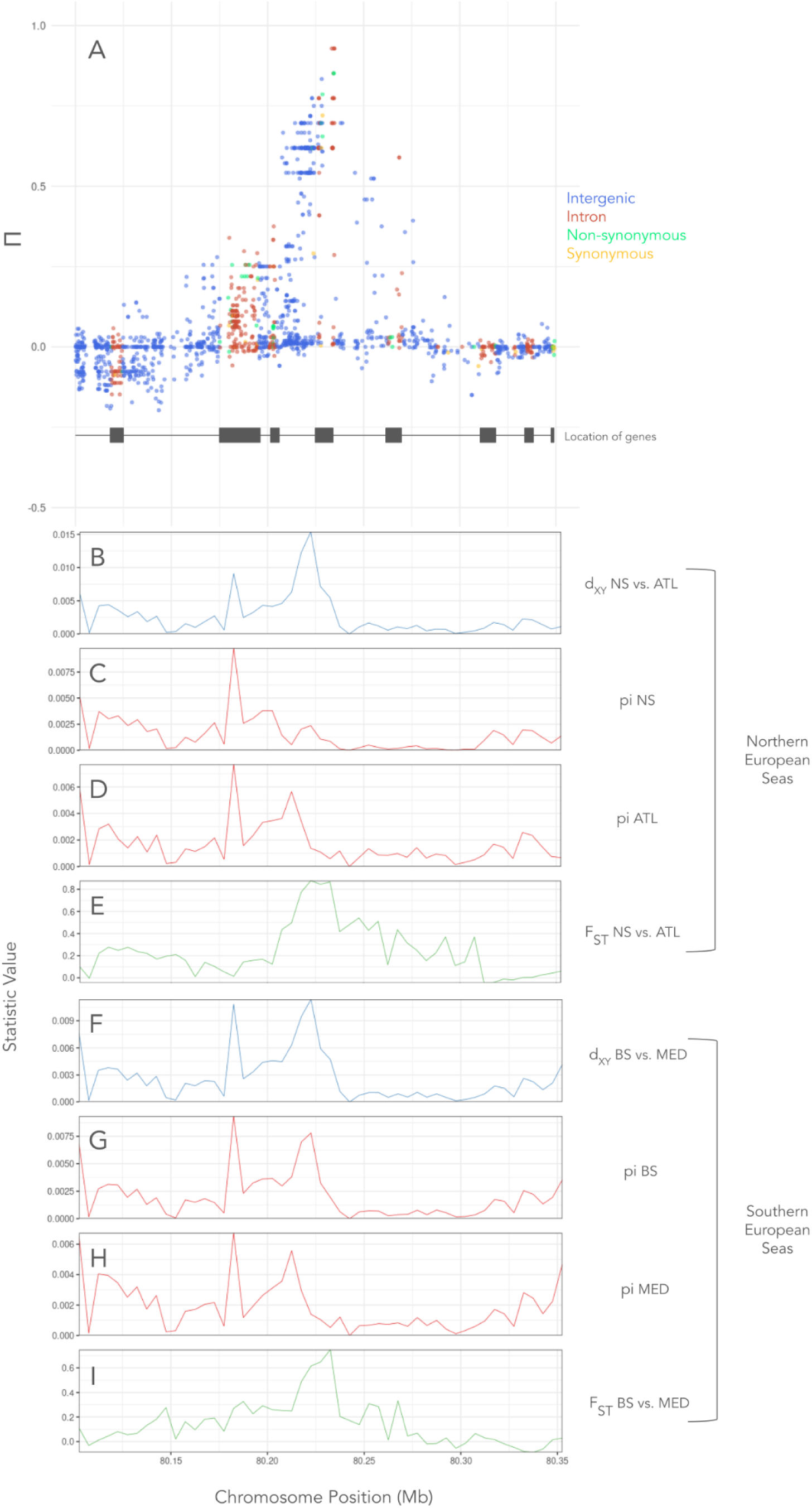
Island of parallelism located on chromosome 5 (Feature I, Island 8). **A** Plot of Π scan with chromosomal position along the x-axis and Π on the y-axis. Points refer to individual SNPs and colours refer to the category of SNP determined by SNPeff (intergenic, intronic, nonsynonymous or synonymous). Grey blocks correspond to the location of genes in the island region. **B** d_XY_ calculated in the NES region between the North Sea and Atlantic populations across the island region. **C** pi calculated for the North Sea population across the island region. **D** pi calculated for the Atlantic population across the island region. **E** Pairwise F_ST_ calculated in the NES region between the North Sea and Atlantic populations across the island region. **F** d_XY_ calculated in the SES region between the Black Sea and western Mediterranean populations across the island region. **G** pi calculated for the Black Sea population across the island region. **H** pi calculated for the western Mediterranean population across the island region. **I** Pairwise F_ST_calculated in the SES region between the Black Sea and Mediterranean populations across the island region.

**Figure 9.**
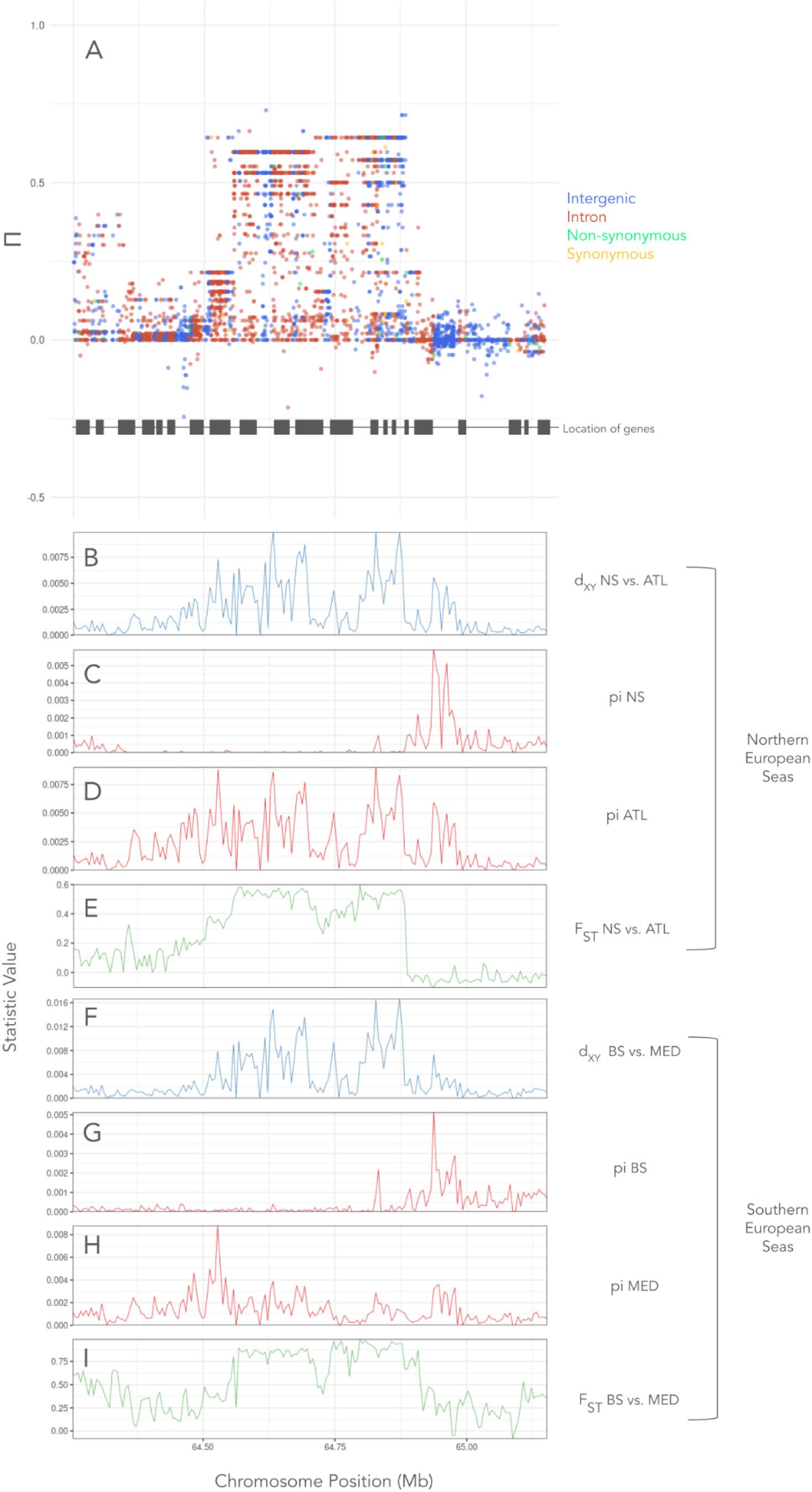
Island of parallelism located on chromosome 3 (Feature F, Island 6). **A** Plot of Π scan with chromosomal position along the x-axis and Π on the y-axis. Points refer to individual SNPs and colours refer to the category of SNP determined by SNPeff (intergenic, intronic, nonsynonymous or synonymous). Grey blocks correspond to the location of genes in the island region. **B** d_XY_ calculated in the NES region between the North Sea and Atlantic populations across the island region. **C** pi calculated for the North Sea population across the island region. **D** pi calculated for the Atlantic population across the island region. **E** Pairwise F_ST_ calculated in the NES region between the North Sea and Atlantic populations across the island region. **F** d_XY_ calculated in the SES region between the Black Sea and western Mediterranean populations across the island region. **G** pi calculated for the Black Sea population across the island region. **H** pi calculated for the western Mediterranean population across the island region. **I** Pairwise F_ST_ calculated in the SES region between the Black Sea and Mediterranean populations across the island region.

Islands ranged in size from approximately 15kb to as large as 300kb. In most islands, the central region of the peak is a genic region (intron, synonymous or nonsynonymous SNPs). We verified that islands were also associated with a peak in pairwise F_ST_ in both geographic regions. In several islands, pairwise d_XY_ was also elevated in both regions (see Fig. 8, and Supplementary Figures 15, 20 & 22), though this was not the case for every island. Estimates of pi showed variation between populations across islands. In some islands, an elevation of pi was present in all populations (Fig. 8 and Supplementary Figures 15 & 22), whereas contrasting signals of pi were seen in many of the islands. For example, in several islands, the Black Sea demonstrated a valley of pi (Supplementary Figures 17 & 21), and in one island a large valley was observed in both the Black Sea and the North Sea populations (Fig. 9). In another island, the North Sea and Black Sea both demonstrated small peaks of pi right at the location of the island peak, despite a valley of pi observed in all populations around the island region (Supplementary Figure 23). The genomic region directly around the single nonsynonymous SNP with a strong parallel signal appeared to have a valley of pi in the Black Sea and western Mediterranean populations, and a decrease in d_XY_ for the same combination of populations (Fig. 7G, Supplementary Figure 25), with the opposite signal in the Atlantic and North Sea populations. The peaks of most islands appear to be narrow, with some exceptions (Supplementary Figure 17), including one where a plateau shaped peak was observed (Figure 9). The size and shape of the island indicates a potential haplotype-block with limited recombination.

Annotated genes associated with the islands of parallelism varied greatly (Supplementary Table 7). However, two of the parallel islands (9 and 12) contained a gene that codes for an ATP synthase subunit, and GO analysis identified one enriched term associated with the island regions, for oxidative phosphorylation, involved in energy metabolism (Supplementary Table 8). To further investigate this, a mitochondrial ML-tree was reconstructed. The tree was split into three clades, one containing the Black Sea individuals, one containing most of the North Sea individuals, and a third containing all individuals from the Atlantic and western Mediterranean, together with a few additional individuals from the North Sea (Fig. 10). We observed amino acid changes at the ATP8 gene only. The Black Sea clade is characterized by two amino-acid changes in ATP8, with a valine-to-isoleucine substitution at position 23, and serine-by-leucine at position 21. The North Sea clade is associated with a valine-to-leucine substitution at position 23. Thus, the Black Sea and North Sea clades show independent amino-acid substitutions at the same position in ATP8, with the ancestral valine replaced by isoleucine and leucine, respectively.

**Figure 10.**
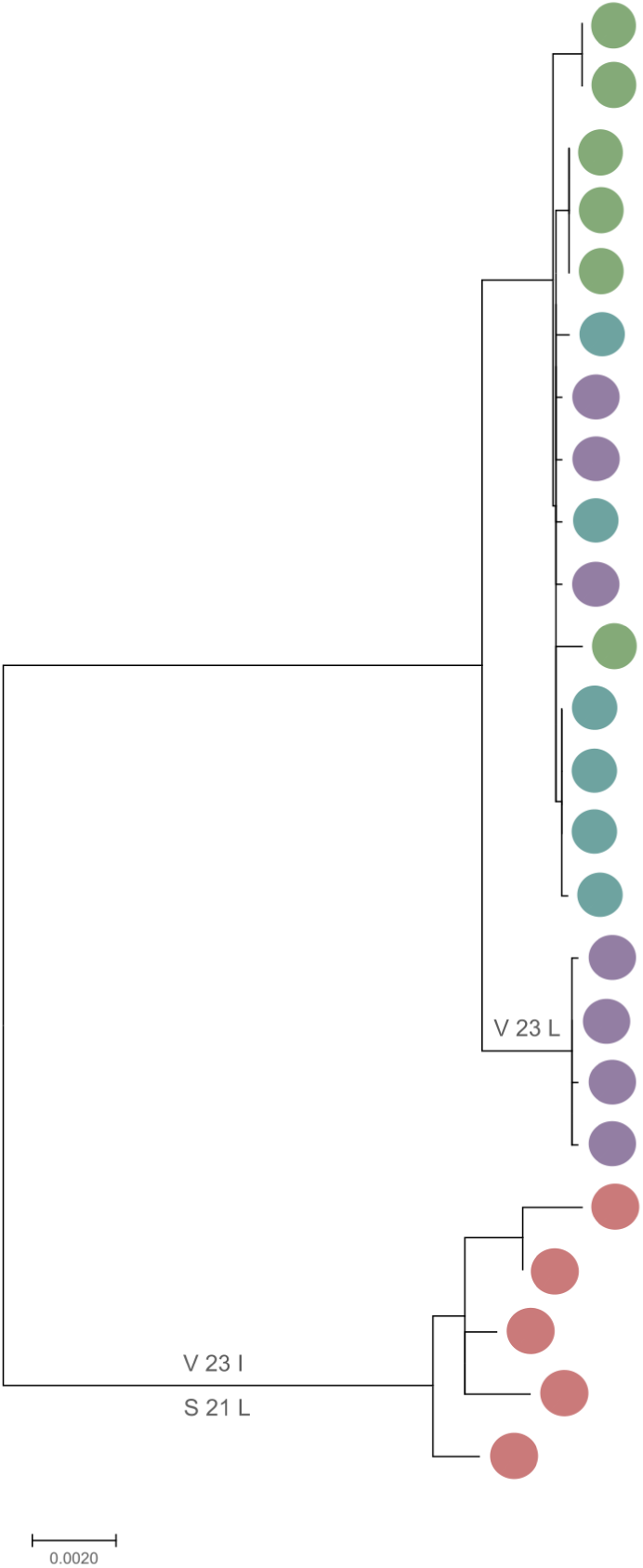
Mitochondrial tree for *Ostrea edulis* samples. Population indicated by colour (Blue: Atlantic, Purple: North Sea, Green: western Mediterranean, Red: Black Sea). Labels indicate ATP8 amino acid changes: valine at position 23 is replaced by leucine in the North Sea group while valine is replaced by isoleucine at position 23 and serine by leucine at position 21 in the Black Sea group.

**Figure 11.**
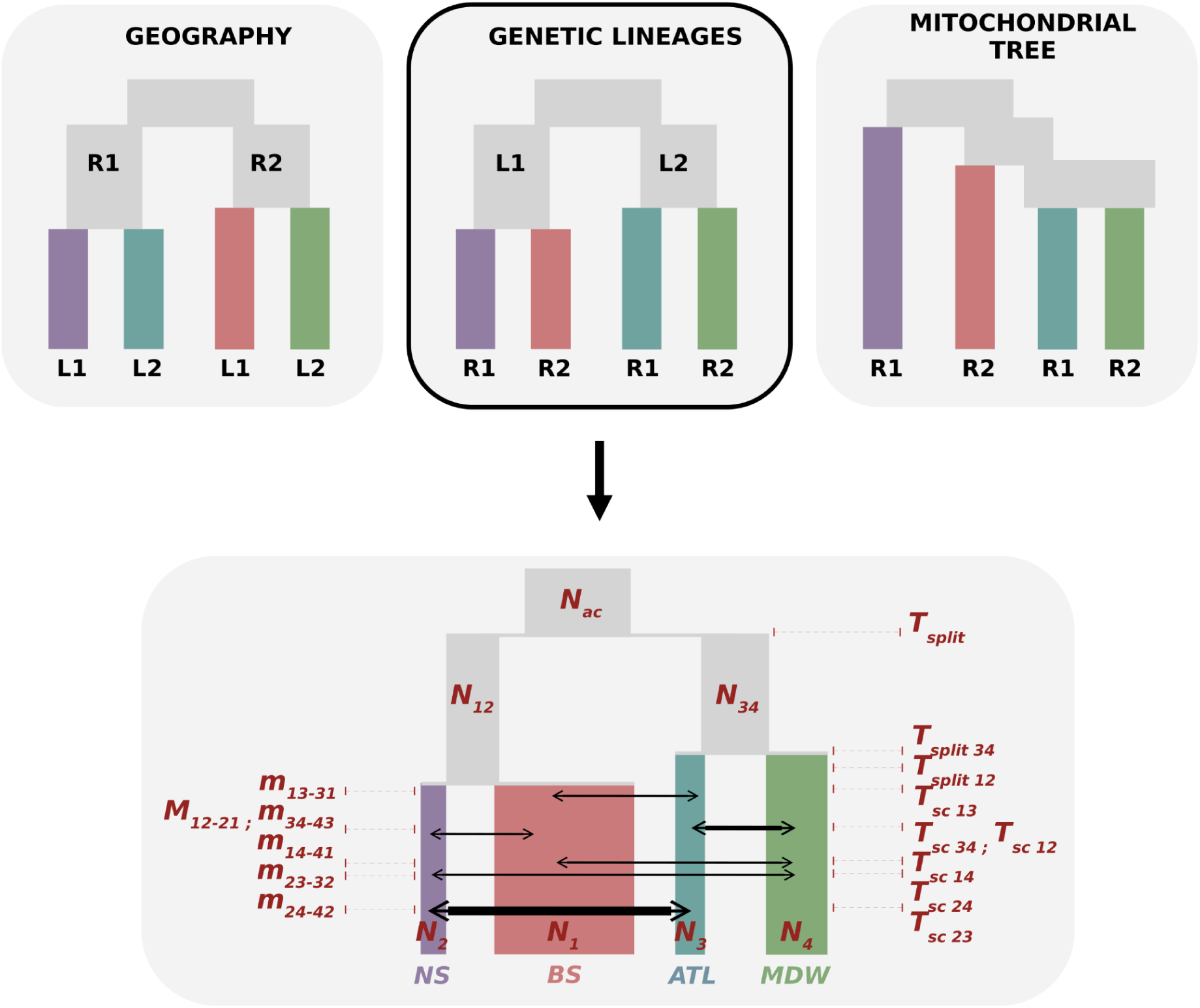
Demographic inferences with DILS for the four-population models. The three tested topologies are shown: geography, genetic lineages, and the mitochondrial tree. These topologies differ in the order of population splits. Additional labels indicate proposed regional (R1, R2) and lineage (L1, L2) assignments. Population indicated by colour (Blue: ATL/Atlantic, Purple: NS/North Sea, Green: MDW/western Mediterranean, Red: BS/Black Sea). For the best-fitting topology (genetic lineages), a graphical representation of the inferred demographic history is provided, together with the associated parameters: N, effective population sizes; T, times of population splits and secondary contacts; and M, migration rates. Estimated parameter values are provided in Supplementary Figure 28.

### Demographic history

We performed pairwise demographic inferences with DILS between populations within geographic regions (the North Sea vs. the Atlantic, the western Mediterranean vs. the Black Sea), and within genetic lineages across regions (the North Sea vs. the Black Sea, the Atlantic vs. the western Mediterranean). DILS showed high posterior probabilities for the best models, and model selection was consistent across replicate runs (Supplementary Table 9). The hierarchical model testing demonstrated strong support for a model of current migration between the populations, with all four pairwise comparisons showing high posterior probabilities for the “current migration” model. All four comparisons also showed consistent posterior probabilities for a model of heterogeneous effective migration across the genome, and homogenous *Ne* across the genome. This is consistent with a substantial effect of barriers to gene flow on patterns of genetic diversity in *O. edulis*, while background selection appears to play a secondary role. All pairwise comparisons demonstrated support for a model of “secondary contact”, with the exception of the comparison between the Black Sea and the western Mediterranean which better fitted the “isolation with continuous migration model”. However, the posterior probabilities for this latter comparison were generally weaker compared to other population comparisons, suggesting that DILS struggled to differentiate between the two models of current migration for this specific pair. Overall, there was strong support for a secondary-contact model. Results of the hierarchical model testing in the pairwise comparisons were used to constrain the parameter space of the four-population inference. The joint evolutionary history of all four *O. edulis* populations better suited the “genetic lineage” topology (Supplementary Table 10), such that an ancestral lineage first split into two clades (L1 = the North Sea + the Black Sea vs. L2 = the Atlantic + the western Mediterranean) which subsequently split into two regions (R1 = Northern European Seas vs. R2 = Southern European Seas). We noted, however, that the posterior probability of the best-supported topology (“genetic lineage”) was only 0.50-0.54, and that this topology was recovered as the best-supported topology in only two of the three replicate runs, with the “mitochondrial tree” being favored in the third. This suggests that the three topologies are difficult to distinguish and that the data provide limited support for a single evolutionary scenario, likely due to the high level of gene flow between *O. edulis* populations at the scale of the European basin. The best-fitting model provides a reasonably good overall fit to the data (Supplementary Figure 26), although diversity levels are consistently overestimated (Supplementary Figure 27). Parameter estimates under the best-fitting model (Supplementary Figure 28) indicate high levels of gene flow between all four population pairs, with migration rates around 40 (in unit *4.N.m*). Gene flow is even higher between the Atlantic and North Sea populations, suggesting that the latter population has been mostly swamped by gene flow from the Atlantic, and potentially explaining the difficulty in distinguishing between the “genetic lineage” and the “mitochondrial tree” topologies. The heterogeneity of effective migration rates was consistently represented by a U-shaped distribution, while the heterogeneity of *Ne* across the genome was best-fitted by a right-skewed exponential distribution. Those results are consistent with the findings in the pairwise comparisons for a minor role of within-population linked selection relative to barriers to gene flow in shaping genetic patterns. Also consistent with the pairwise comparisons is the high levels of gene flow, clear signals of secondary contact and split times between the two genetic lineages (L1 vs. L2) consistent with isolation during the last glacial period (∼200 kya). The ratios of the time of secondary contact to the time of split ranged from 30 to 50%, compatible with isolation during the last glacial period.

## DISCUSSION

In this study, we used short-read whole-genome sequencing on 27 *O. edulis* samples to revisit population structure and three large chromosomal inversions previously identified using limited numbers of SNPs or low-coverage genomic data (Lapègue et al. 2023, Alves-Monterio et al. 2024, Sambade et al. 2025). We further characterized the population-level distribution of these inversions and investigated their age and genomic impact. Building on previous evidence for parallel differentiation across the species range (Lapègue et al. 2023), we identified genomic regions contributing to this parallelism and assessed the relative contribution of chromosomal inversions and smaller genomic islands. Finally, demographic inference supported the presence of two cryptic semi-isolated lineages with genomes largely porous to gene flow, and allowed us to investigate the evolutionary history underlying these contrasting patterns of genomic differentiation.

### Chromosomal inversions support the range-margin versus range-center lineage hypothesis

Combining our results with the broader geographic sampling of Alves-Monteiro et al. (2024) now reveals that inversion allele frequencies are more consistent with a lineage structure separating range-margin from range-center populations than with a simple geographic latitudinal gradient. For chromosome 4, the inversion allele fixed in our western Mediterranean samples is also fixed in the Adriatic, Corsican, and Sardinian samples of Alves-Monteiro et al. (2024). For chromosome 5, the inversion allele fixed in our Black Sea samples is otherwise found at high frequency in northern Europe, mainly in Scandinavia (Alves-Monteiro et al. 2024). Most strikingly, the chromosome 9 inversion allele (chromosome 8 in Alves-Monteiro et al. 2024, who used a different reference genome), fixed in both our Black Sea and North Sea samples, is also fixed or nearly fixed across Scandinavia. Thus, although inversion allele frequencies are strongly geographically structured (Alves-Monteiro et al. 2024; Sambade et al. 2025), the inclusion of the Black Sea reveals a pattern that cannot be readily explained by geographic distance or a simple latitudinal environmental gradient alone. Instead, it supports the two-lineage structure proposed by Lapègue et al. (2023), with Lineage 1 represented by range-margin populations and Lineage 2 by range-center populations. The broader sampling of Alves-Monteiro et al. (2024) suggests that Scandinavian populations predominantly belong to Lineage 1, although their differentiation from the North Sea population deserves further investigation, suggesting that our samples from the Netherlands may represent an admixed edge of Lineage 1 rather than a good reference of the lineage as a whole. Populations from the British Isles to the Adriatic Sea predominantly belong to Lineage 2, with a genetic break around Galicia separating two genetic clusters. The high porosity of genetic barriers may generate gradients of interlineage introgression within each genetic cluster, producing heterogeneous isolation-by-distance patterns across the genome that can mimic local adaptation (Gagnaire et al. 2015; Robinet et al. 2020).

The three inversion polymorphisms also appear to be ancient. Nucleotide diversity differed only slightly between inversion alleles on chromosomes 4 and 5, providing no consistent indication of which allele is ancestral, in agreement with Alves-Monteiro et al. (2024). More importantly, divergence between inversion alleles was high, and our IBS trees consistently grouped *O. chilensis* with one of the two alleles. For all three inversions, divergence between inversion alleles was approximately two to three times greater than divergence between *O. chilensis* and the more closely related *O. edulis* inversion allele, suggesting that the three inversion polymorphisms predate the divergence between *O. edulis* and *O. chilensis*, estimated at approximately 20 million years ago (Li et al. 2021). The inversion allele associated with *O. chilensis* was the less diverse allele on chromosome 4 but the more diverse allele on chromosome 5, further showing that nucleotide diversity provides little indication of which inversion allele is ancestral or derived. An ancient origin was also proposed by Sambade et al. (2025), although diversity patterns led them to suggest that the southern alleles are ancestral, a conclusion that our results do not allow us to confirm.

Finally, we found little functional evidence for divergent selection or relaxed purifying selection acting specifically on the inversions. None contained significantly more high- or moderate-impact variants than collinear regions, and functional enrichment was limited and inconclusive. Sambade et al. (2025) found stronger evidence for functional enrichment and proposed that divergent selection contributes to the geographic distribution of the three inversions, with possible associations with zinc pollution, salinity, and prevalence of the parasite *Bonamia ostreae*. Our results do not exclude a contribution of divergent selection, but suggest that the long evolutionary history of the inversions mostly contribute to their divergence.

Provided that the inversions are ancient and reduced recombination preserves divergence across entire inversions, they are expected to generate strong outlier signals in sliding-window analyses. The sharing of inversion alleles between the Black Sea and North Sea or Scandinavian populations despite their geographic and environmental separation suggests that this strong differentiation partly reflects shared evolutionary history preserved despite gene flow.

### SNP-level analysis reveals parallel differentiation beyond chromosomal inversions

Chromosomal inversions are often expected to generate strong signals of parallel evolution (Westram et al. 2022). In *O. edulis*, however, only the chromosome 9 inversion showed a strong parallel signal, whereas the strongest signals occurred in much smaller genomic islands outside inversions. The long-snouted seahorse provides an interesting comparison because its population structure closely resembles that of *O. edulis*, with two differentiated ecotypes in the Mediterranean Sea and two Atlantic lineages, including a northern Atlantic lineage genetically related to the Mediterranean lagoon ecotype (Riquet et al. 2019). In this system, a genome-wide sliding-window analysis focused primarily on the contribution of large chromosomal inversions to parallel differentiation (Meyer et al. 2024). Although this approach was not designed to identify small differentiated islands, more localized signals outside the major inversions were nevertheless visible in the genome-wide differentiation profiles. This suggests that finer-scale, SNP-based analyses may reveal additional components of parallel differentiation beyond the prominent contribution of inversions. A similar pattern emerges from recent studies in *Littorina*, where inversions contain a large fraction of differentiated SNPs but localized signals of parallel differentiation are also found in collinear regions (Morales et al. 2019; Koch et al. 2022). These studies emphasize complementary scales of a genomic architecture in which large inversions coexist with more localized signals of differentiation. The apparent contribution of inversions may also depend on how genomic differentiation is quantified. Using a SNP-based statistic analogous to ours, Pajot et al. (2026) reported that 96% of highly differentiated SNPs between parallel ecotypes of *Littorina fabalis* occurred within chromosomal inversions. However, large inversions in strong linkage disequilibrium contain many correlated differentiated SNPs and are therefore disproportionately weighted when contributions are measured by counting outlier SNPs. Their scan nevertheless revealed approximately 15 localized peaks of parallel differentiation outside inversions, suggesting a more balanced contribution when distinct genomic signals rather than individual SNPs are considered. Together with our results, these examples suggest that the apparent predominance of inversions in parallel evolution may depend partly on the genomic scale of analysis (sliding-window vs SNP-based) and on how parallel differentiation is quantified (proportion of differentiated SNPs vs number of independent differentiation peaks).

The 12 islands showed different patterns of genetic diversity and divergence, suggesting different evolutionary histories. Islands 3 and 6 showed haploblock-like structures associated with strong linkage disequilibrium, and Island 6 may correspond to a small structural variant. Local reductions in pi and d_XY_ in these and several other islands are compatible with relatively recent selective sweeps rather than long-term barriers to gene flow. In contrast, most islands showed peaks of F_ST_ accompanied by increased d_XY_ and no corresponding valley of within population diversity, consistent with older differentiation maintained despite gene flow (Cruickshank and Hahn 2014; Van Belleghem et al. 2018). Thus, parallel differentiation in *O. edulis* appears to combine signals generated by different evolutionary processes over different timescales, as observed in other species (e.g. sea bass, Duranton et al. 2018).

Interestingly, islands were absent from chromosomes 7 and 8, while the weakest signal was observed on chromosome 6. These three chromosomes were previously found to carry more polymorphic duplications and repetitive sequences than other chromosomes and may also be prone to somatic aneuploidy (Colston-Nepali, Lapègue and Bierne 2026). Although tentative, this pattern suggests that the peculiar genomic properties of these three chromosomes may interfere with the detection of parallel SNP differentiation.

Finally, two islands map nuclear genes encoding ATP synthase subunits, and oxidative phosphorylation was enriched among genes within the islands. The mitochondrial genome does not reproduce the nuclear lineage structure exactly. Atlantic and western Mediterranean mitogenomes cluster together, as expected, but North Sea mitogenomes are heterogeneous, with some clustering with this group and others forming a distinct subclade, yet remaining more closely related to them than to the Black Sea mitogenomes. Nevertheless, ATP8 shows an intriguing pattern of convergent evolution, with the North Sea and Black Sea mitochondrial lineages carrying independent amino-acid substitutions at the same position. Although the North Sea variant is not fixed in our samples, this population may represent an admixed edge of the range-margin lineage, as suggested above, and fixation is rarely observed at any of the islands identified here. Evidence for mitonuclear incompatibility therefore remains tentative. However, the two nuclear ATP synthase subunits identified in independent islands (O and β) are components of the same protein complex, with subunit O physically interacting with the catalytic F1 head containing subunit β. Their parallel differentiation therefore raises the possibility of coevolution and genetic incompatibility between these two nuclear-encoded components of the ATP synthase complex. The convergent amino-acid change in mitochondrial ATP8 further suggests that mitonuclear interactions may also contribute to, or be influenced by, this pattern.

### Evolutionary history underlying the genomic landscape

Our demographic analyses are consistent with historical isolation between two *O. edulis* lineages followed by secondary contact and progressive erosion of differentiation across most of the genome. Pairwise comparisons consistently supported heterogeneous gene flow across the genome, and secondary contact was the best-supported scenario in three of four comparisons. Models with homogeneous *Ne* across the genome were also consistently favoured, suggesting that heterogeneous differentiation is primarily associated with variation in gene flow rather than linked selection within populations. At the four-population level, demographic inference favoured the lineage topology expected under the “swamping hypothesis” (islands stay, background moves), although extensive gene flow, especially between our Atlantic and North Sea samples, reduces confidence in the demographic reconstruction. Following post-glacial contact, expansion of the range-center lineage may have generated and pushed two sister contact zones towards northern Europe and the eastern Mediterranean. Because genetic barriers are highly porous, gene flow can swamp historical differentiation at neutral loci, reorganizing genetic variation according to geography while preserving the original lineage structure at barrier islands (Bierne, Gagnaire and David 2013; Gwee et al. 2025). This history may explain both the geographic distribution of inversion alleles and the islands showing elevated d_XY_, where differentiation appears to have persisted over long timescales despite gene flow.

The lineage topology should not, however, be interpreted as evidence that the two range-margin populations formed a single homogeneous ancestral population. The eastern Mediterranean provides the clearest present-day representative of the range-margin lineage, while the relationship between eastern Mediterranean and northern European populations may partly reflect ancient gene flow within an already structured ancestral range. This interpretation is consistent with the mitochondrial topology, which received only slightly less support than the lineage topology in four-population models and suggests a more complex relationship among range-margin populations. Similar histories have been inferred in other European marine species. In *Mytilus* mussels, alleles introgressed from *M. edulis* (found today from north to southwest France) during ancient contacts with Mediterranean *M. galloprovincialis*, have been identified despite the absence of contemporary contact between these populations (Gosset and Bierne 2013; Fraïsse et al. 2016), with the strongest signatures of ancient *M. edulis* introgression detected as far as the Black Sea (Simon et al. 2021). European sea bass similarly retain complex geographic patterns of ancestry generated by secondary contact and subsequent diffusion of introgressed ancestry across their range (Tine et al. 2014; Duranton et al. 2018). Thus, a shared evolutionary history of the range-margin populations does not imply the absence of ancestral structure and past gene flow between them.

The timing inferred from our different demographic approaches is also broadly compatible with isolation during the last glacial period followed by post-glacial contact. The four-population model yielded time of secondary contact (Tsc) to time of split (Tsplit) ratios of approximately 0.3. Interestingly, similar estimates have been obtained in European sea bass using allele-frequency demographic inference (Tine et al. 2014) and the length of identity-by-state tracts (Duranton et al. 2018). Extensive simulations by Roux et al. (2016) showed that the Tsc/Tsplit ratio can often be estimated more confidently than either Tsplit or Tsc separately. Rather than attempting to infer a precise divergence time, we can therefore use this ratio to constrain the period of isolation. If secondary contact occurred no later than the Last Glacial Maximum, approximately 20 kya, this ratio would place an upper bound on lineage divergence at approximately 67 kya, strikingly close to the preceding extreme glacial phase around 70 kya. This correspondence suggests that the inferred divergence and secondary contact may instead reflect the most recent episode in a longer glacially driven alternation of divergence and contact (Hewitt 2004). If gene flow largely resets genome-wide divergence at each contact, as we suggest here, earlier cycles may leave little detectable trace, and the inferred divergence time may primarily reflect the time elapsed since the end of preceding contact (Fraïsse et al. 2018; Duranton et al. 2018).

The “introgression hypothesis” cannot, however, be excluded for every individual island. Locally adaptive alleles can move between similar selective environments through intervening populations in which they are counter-selected (Schluter and Conte 2009; Bierne, Gagnaire and David 2013; Welch and Jiggins 2014). Alleles favoured in both the North Sea and Black Sea could therefore have moved between these distant environments through Atlantic and Mediterranean populations, especially given that shared polymorphisms are found across the *O. edulis* genome, including within islands of differentiation. This scenario may be particularly relevant to islands showing reduced d_XY_ and pi, including cases where reduced diversity is stronger in the Black Sea than in the North Sea and may reflect a more recent sweep in the former. Similar selective pressures in the North Sea and Black Sea could also contribute to parallel differentiation independently of demographic history. Thus, both processes may have contributed to the present genomic landscape, but demographic reconstruction and the persistence of old divergence at most islands favour porous genetic barriers and genome-wide swamping as the predominant explanation for the parallel structure observed today to be restricted on a few islands only.

## Acknowledgements

The authors are very grateful to Stéphane Pouvreau, GAEC Le Rocher, Camille Saurel, DTU, and Pauline Kamermans for carrying out the sampling of the flat oyster samples used in this study.

Thank you to Pierre Alexandre Gagnaire, Bruno Ernande, and other members of the ISEM BEM team, and LEGPM MARBEC team for valuable discussion.

The authors acknowledge the Pôle de Calcul et de Données Marines (PCDM; https://wwz.ifremer.fr/en/Research-Technology/Research-Infrastructures/Digital-infrastructures/Computation-Centre) for providing DATARMOR computing and storage resources.

## Data availability statement

Raw data from the short-read whole genome sequencing is available at: https://www.ebi.ac.uk/ena/browser/view/PRJEB86116

Code is available at: https://doi.org/10.5281/zenodo.22902114

## Authors contributions

Conceptualization: LCN, CF, NB, SL

Data curation: LCN, CF, CReisser, SL

Formal analysis: LCN, PDV, CReisser, CF, NB, SL

Funding acquisition: NB, SL

Investigation: LCN, PDV CF, NB, SL

Methodology: LCN, CRoux, CF, NB, SL

Project Administration: LCN, NB, SL

Resources: CReisser, FC, SL

Software: BP, CRoux, CF

Supervision: CF, NB, SL

Validation: LCN, CF, NB, SL

Visualization: LCN

Writing (original draft)- LCN, CF, NB, SL

Writing (review and editing)- all authors

## SUPPLEMENTARY MATERIAL

### Supplementary Methods

#### DNA preparation and sequencing

Samples from the Black Sea were extracted using Promega kits (Promega corporation, Madison, Wisconsin, USA) and QIAamp DNA minikits (Qiagen, Hilden, Germany) in 2009 and 2011. All other samples were extracted using Nucleobond HMW DNA kits (Macherey-Nagel, Düren, Germany) in 2020. Extracted DNA was sequenced using an Illumina Novaseq 6000 (paired-end, 2 x 150bp) by Integragen (Evry, France).

#### Preliminary genome assembly investigation

In 2023, there were four publicly available *O. edulis* reference genomes: the ‘Roscoff’ assembly (Boutet et al. 2022, GenBank assembly: GCA_024362745.1), sampled in Brittany, France, the ‘Roslin’ assembly (Gundappa et al. 2022, GenBank assembly: GCF_023158985.2), sampled in Whitstable, UK, the ‘Qingdao’ assembly (Li et al. 2023, GenBank assembly: GCA_032173915.1), sampled from farmed oysters in Grevelingen and Oosterschelde, the Netherlands and the ‘Sanger’ assembly, (Adkins and Mrowicki 2023, GenBank assembly: GCA_947568905.1), sampled in Devon, UK. In order to choose the most appropriate genome for this project, we made thorough comparisons (see Table 1 for assembly statistics). Pairwise comparisons of genome assemblies were visualized using the program D-GENIES (Cabanettes and Klopp, 2018, available at: https://dgenies.toulouse.inra.fr/). The Roscoff and Roslin genomes were both found to contain missing sections of assembly. Due to this, and the fact that at the time of investigation (February 2023) the Sanger genome was not yet annotated, we chose to proceed with the Qingdao genome.

**Table 1.** Assembly statistics gathered from NCBI and associated papers available in February 2023. Missing values indicate a lack of reporting.

|  | Roscoff | Roslin | Qingdao | Sanger |
| --- | --- | --- | --- | --- |
| Genome size | 1 Gb | 935.1 Mb |  | 894.8 Mb |
| Total ungapped length | 1 Gb | 935 Mb | 946 Mb | 894.7 Mb |
| Number of chromosomes | 10 | 10 | 10 | 10 |
| Number of scaffolds | 563* | 1363 | 618 | 52 |
| Complete BUSCOs | 0.95 | 0.94 | 0.95 |  |
| Scaffold N50 | 98.8 Mb | 95.6 Mb | 94.82 Mb | 94.3 Mb |
| Scaffold L50 | 6 | 5 | 5 | 5 |
| Number of contigs | 7754 | 2897 |  | 657 |
| Contig N50 | 95.8 Mb** | 1.8 Mb | 3.23 Mb | 2.4 Mb |
| Contig L50 | 6 | 129 | 86 | 112 |
| GC percent | 35 | 35 | 35 | 35 |
| Protein coding genes | 35962 | 35699 | 34495 |  |
| Fully annotated genes | 24302 | 23109 | 32238 |  |
| Assembly | Chromosome | Chromosome | Chromosome | Chromosome |

*\* discrepancy between numbers reported in paper (563) vs. NCBI (7754). The NCBI number seems more likely to be an error based on the range of numbers reported for the other genomes. \*\* seems likely to be an error (reported on NCBI) in comparison to the range of numbers reported for the other genomes.*

#### Data processing

Raw data was uploaded onto the Ifremer supercomputer, Datarmor (https://www.ifremer.fr/fr/infrastructures-de-recherche/le-supercalculateur-datarmor), and is publicly available at https://www.ebi.ac.uk/ena/browser/view/PRJEB86116. Reads were assessed for quality using the program FastQC (Andrews 2010), and then trimmed using the fastp program (Chen et al. 2018), filtering for an average quality of 28, and removing polyG tails and Illumina adapter sequencing.

#### Variant calling and filtering

Samples were aligned to the Qingdao genome (Li et al. 2023) using the program BWA-MEM (Li and Durbin 2010) version 0.7.17. Mapped reads were then filtered using the program Sambamba (Tarasov et al. 2015) version 0.8.0 to remove reads that did not pair properly, and secondary alignments.

Mapped reads were processed and variant calling completed using the Genome Analysis Toolkit (GATK, McKenna et al. 2010) version 4.4.0.0. Duplicates were marked using Picard Tools (https://broadinstitute.github.io/picard/). Variants were called for each individual sample using the GATK HaplotypeCaller with a minimum base quality score of 30, and an expected heterozygosity value of 0.01 used to compute prior probability that a locus is non-reference. GenomicsDBImport and GenotypeGVCFs were used to perform joint genotyping of all samples. Single nucleotide polymorphisms (SNPs) only were selected using SelectVariants, and variants were hard filtered with VariantFiltration using the recommended parameters (QD < 2.0, QUAL < 30.0, SOR > 3.0, FS > 60.0, MQ < 40.0, MQRankSum < −12.5, ReadPosRankSum < −8.0).

*Ostrea edulis* has a high incidence of polymorphic duplications (Colston-Nepali, Lapègue and Bierne 2026), which may violate the assumptions of downstream analyses for population structure and demographic history. For this reason, additional filtering steps were designed for this project (outlined in the Data Processing Appendix). Final filtering parameters are described here, and two datasets were created in order to best meet the assumptions of different downstream analyses (as recommended by Hemstrom et al. 2024), the “Genome Scan Dataset” and the “Demographic History Dataset”.

##### The Genome Scan Dataset

The objective for this dataset was to maintain a high SNP density, so less stringent filtering was applied. Though some less reliable SNPs (sequencing errors, mapping errors, paralogs etc.) may remain in this dataset, information about neighbouring SNPs should lend confidence to the observed signal. Only biallelic SNPs were included. The program ngsParalog (Linderoth 2018) was used to identify ‘deviant’ SNPs due to sequence differences between paralogous sequences, sometimes called paralogous sequence variants (PSVs). Briefly, ngsParalog uses the coverage, the allele coverage fraction, and the excess heterozygosity in the sample in a likelihood framework to identify SNPs that deviate from the expectation of mendelian diploid genotypes in single copy orthologs (see Appendix Data Processing section 1). VCFtools (Danecek et al. 2011) v.0.1.16 was used to remove deviant SNPs, and sites identified as potentially problematic (Colston-Nepali, Lapègue and Bierne 2026; see Appendix Data Processing section 2). All sites with any missing data were removed, and sites were filtered for a minimum minor allele frequency of 5%. A second Genome Scan Dataset including invariant sites in addition to biallelic SNPs was created for some downstream analyses. The final dataset consisted of 6,150,684 variant sites and 838,424,559 invariant sites.

##### The Demographic History Dataset

In this dataset the objective was to obtain a large number of reliable SNPs, while also seeking to gain an idea of the number of bases actually sequenced by calling invariants, without which we would risk introducing bias into the estimation of nucleotide diversity (Korunes and Samuk 2021). GenomicsDBImport and GenotypeGVCFs in GATK was re-run to include a sample of the species *O. chilensis* as an outgroup. Chromosomes 6, 7 and 8 were excluded as these chromosomes have a significantly higher density of polymorphic duplications (Colston-Nepali, Lapègue and Bierne 2026). Site depth of invariant sites was assessed, and minimum mean depth per site was filtered to the 25th percentile (6), and a maximum of 10 times the mean (117). For the variant site filtration, only biallelic SNPs were selected. ngsParalog (Linderoth 2018) was used to mark regions estimated to be problematic (see Appendix Data Processing). VCFtools (Danecek et al. 2011) v.0.1.16 was used to remove these sites. Site depth was further filtered to two standard deviations below and above the mean (minimum mean depth 7, and maximum mean depth 23). Inversion regions (identified in analyses using the Genome Scan Dataset) were excluded as their special genetic features are not explicitly modelled in the demographic analyses. The final dataset consisted of 2,941,404 variant sites and 439,271,217 invariant sites. The removed sites were encoded as missing data using a custom python script.

### Supplementary Population Structure Analysis

#### METHODS

Following the processing of mapped reads, and variant calling completed using the Genome Analysis Toolkit (GATK, McKenna et al. 2010) version 4.4.0.0, a population structure dataset was created. The objective for this dataset was to ensure accurate analyses of population structure, so stringent filtering was applied to retain only the most reliable SNPs and eliminate the effect of linkage disequilibrium. Only variant sites and biallelic SNPs were included. Chromosomes 6, 7 and 8 were excluded as these chromosomes have a significantly higher density of polymorphic duplications (Colston-Nepali, Lapègue and Bierne 2026). The program ngsParalog (Linderoth 2018) was used to identify ‘deviant’ SNPs due to sequence differences between paralogous sequences, sometimes called paralogous sequence variants (PSVs). Briefly, ngsParalog uses the coverage, the allele coverage fraction, and the excess heterozygosity in the sample in a likelihood framework to identify SNPs that deviate from the expectation of mendelian diploid genotypes in single copy orthologs (see Appendix Data Processing section 1). VCFtools (Danecek et al. 2011) was used to remove deviant SNPs, and sites identified as problematic in investigations using high coverage individuals (see Appendix Data Processing section 2). Of the remaining sites, only those that passed individual coverage filtering (see Appendix Data Processing section 3) were retained. Site depth was further filtered to two standard deviations below and above the mean (minimum mean depth 19, and maximum mean depth 24). Sites were filtered for a minimum minor allele frequency of 5%. All sites with any missing data were removed and BCFtools (Danecek et al. 2021) was used to prune sites for linkage disequilibrium, using a minimum r2 value of 0.6 and a window size of 1000 SNPs. The final dataset consisted of 423,929 variant sites.

To validate the structure previously identified with few SNPs by Lapègue et al. (2023), a Principal Component Analysis (PCA) was run on the Population Structure Dataset using the R package SNPRelate (R Core Team 2024, Zheng et al. 2012).

#### RESULTS

Principal components analysis corroborated population designations by Lapègue et al. (2023), with samples from the Atlantic, North Sea, western Mediterranean, and Black Sea forming distinct clusters (Fig. 1). The first principal component seems driven by geography, with the Atlantic and North Sea populations falling in the same place, the Black Sea the most differentiated, and the western Mediterranean falling in the middle. The second principal component seems to differentiate populations within regions: the North Sea and the Black Sea, both belonging to the genetic lineage at the edges of the range (here numbered lineage 1), sit higher up on the second axis than their regional counterparts, which belong to the other lineage (here numbered lineage 2).

**Figure 1.**
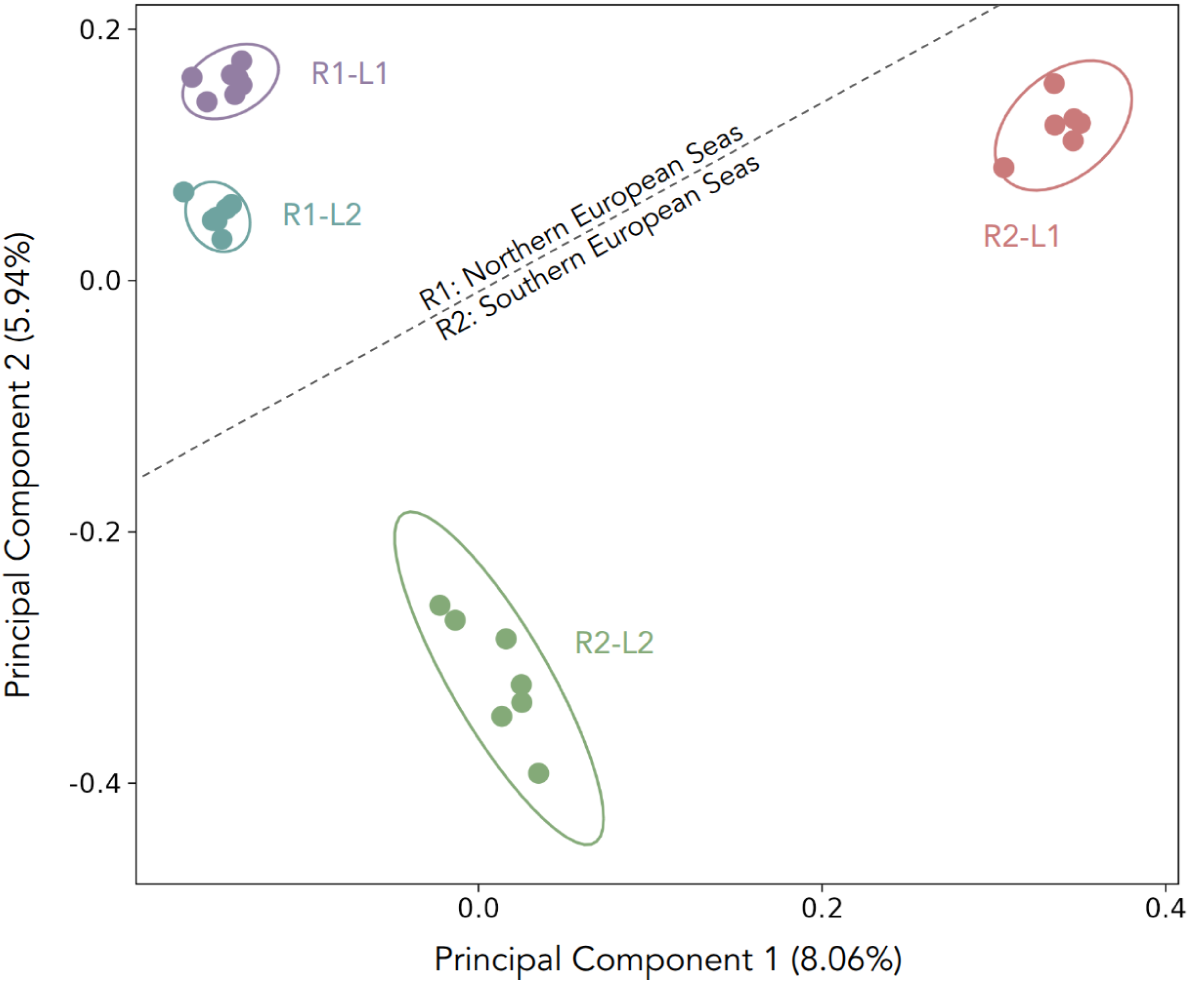
Principal components analysis of the newly sequenced individuals using 423,929 SNPs. Population indicated by colour (Blue: Atlantic, Purple: North Sea, Green: western Mediterranean, Red: Black Sea). Dashed line indicating proposed regional split between Northern European Seas and Southern European Seas. Additional labels indicating proposed regional assignments (R1, R2), and proposed lineage assignments (L1, L2).

##### Data Processing Appendix

SNP lists of possibly problematic SNPs were created to aid filtering.

###### 1. ngsParalog

The program ngsParalog (Linderoth 2018) was used to identify potentially problematic genomic regions that may be caused by repetitive sequence or paralogy using coverage, excess heterozygosity, and likelihood modeling. Including these SNPs can have a large effect on metrics of population genetics (Dallaire et al. 2023). The program samtools mpileup (Danecek et al. 2021) was used to create the input files for ngsParalog, and then the function calcR was used to calculate the likelihood of deviancy for SNPs. The Bonferroni method was used with a p-value of 0.05 to categorize SNPs as ‘deviant’. A list of ∼2.4 million SNPs classified as ‘deviant’ was created.

###### 2. Using the five high coverage individuals

VCF files of the five high coverage individuals used for reference based analyses in Colston-Nepali, Lapègue and Bierne (2026) were used to identify SNPs with more than 10% discordance from expected 0.5 allele coverage fraction, and all SNPs with coverage of less than 1X or more than 3X were selected for each individual. These SNP lists were combined and the final SNP list was saved as ‘highcoverage_SNPs.txt’.

###### 3. Filtering by individual coverage

Analyses by Colston-Nepali, Lapègue and Bierne (2026) indicate potentially problematic SNPs at a coverage that deviates from expected 2X. Coverage filtering was applied globally on the full dataset, but individual level coverage filtering may further reduce problematic SNPs. Average coverage across individuals was 18, but standard deviation was high (+/− 7). Depth per site was calculated for each individual using VCFtools (Danecek et al. 2011), and histograms plotted in R (R Core Team 2016). The mode of the histograms was used to estimate the 2X coverage. Each individual was then filtered to include only sites that fell between 1.5X and 3X coverage. As a result, individuals lost between 30-49% of SNPs. The individual datasets were re-merged and filtered for a maximum of 20% missing data. The ∼5million SNPs that remained had therefore passed coverage filtering in at least 80% of the individuals. A SNP list was created for these remaining SNPs, and saved as ‘individualcoverage_SNPs.txt’.

### Demographic Inference Appendix

#### DILS configuration parameters used

region: type of genomic region sequenced [coding or noncoding].
lightMode: TRUE for a light analysis (on a laptop), FALSE for a better exploration of the parameters space [TRUE or FALSE]
useSFS: use the SFS in the 2-population inference [1]; do not use it [0].
population growth: temporal variation in the population size [constant or variable].
modeBarrier: type of modelling for the genomically heterogeneous Me [bimodal or beta].
max N tolerated: maximum proportion of N/gaps allowed in the sequence of a locus
Lmin: minimum number of treatable sites allowed in the sequence of a locus
nMin: minimum number of sequences allowed per locus and per population/species *If the number of sequences is higher, then DILS performs a draw of nMin sequences (without replacement) from the set of available sequences*.
mu: mutation rate per site per generation
rho over theta: the ratio of recombination over mutation
N min: minimum prior number of diploid individuals
N max: maximum prior number of diploid individuals
Tsplit min: minimum prior number of generations for time of split
Tsplit max: maximum prior number of generations for time of split
M min: minimum prior number of migrants per generation
M max: maximum prior number of migrants per generation

#### Final configuration files for each of the pairwise comparisons

Black Sea vs. western Mediterranean:

region: noncoding
lightMode: FALSE
useSFS: 1
population growth: constant
modeBarrier: beta
max N tolerated: 0.4
Lmin: 500
nMin: 8
mu: 0.00000002763
rho over theta: 0.01
N min: 100
N max: 100000
Tsplit min: 100
Tsplit max: 250000
M min: 0.4
M max: 100

North Sea vs. Atlantic

region: noncoding
lightMode: FALSE
useSFS: 1
population growth: constant
modeBarrier: beta
max N tolerated: 0.4
Lmin: 500
nMin: 8
mu: 0.00000002763
rho over theta: 0.01
N min: 100
N max: 100000
Tsplit min: 100
Tsplit max: 250000
M min: 0.4
M max: 100

North Sea vs. Black Sea

region: noncoding
lightMode: FALSE
useSFS: 1
population growth: constant
modeBarrier: beta
max N tolerated: 0.4
Lmin: 500
nMin: 8
mu: 0.00000002763
rho over theta: 0.01
N min: 100
N max: 100000
Tsplit min: 100
Tsplit max: 250000
M min: 0.4
M max: 100

Atlantic vs. western Mediterranean

region: noncoding
lightMode: FALSE
useSFS: 1
population growth: constant
modeBarrier: beta
max N tolerated: 0.4
Lmin: 500
nMin: 8
mu: 0.00000002763
rho over theta: 0.01
N min: 100
N max: 100000
Tsplit min: 100
Tsplit max: 250000
M min: 0.4
M max: 100

#### Final configuration file for the four-population comparison

region: noncoding
nameA: Black Sea
nameB: North Sea
nameC: Atlantic
nameD: western Mediterranean
maxN: 0.4
nMin: 8
Lmin: 500
mu: 0.00000002763
rec: 0.0000000002763
N_bound_min: 0.01 #i.e. 0.01 * Nref(100000) = 1000
N_bound_max: 0.5 #i.e. 0.5 * Nref(100000) = 50000
T_bound_min: 0.01 #i.e. 0.01 * Nref(100000) = 1000
T_bound_max: 2.0 #i.e. 2.0 * Nref(100000) = 200000
M_bound_min: 0.4
M_bound_max: 100
shape_bound_min: 0.01 #min bound of shape parameters (beta distribution)
shape_bound_max: 1 #max bound of shape parameters (beta distribution)
topologies: topo1,topo2,topo3 #statistical comparison of the three topologies
migAB: 1 #migration rate between A and B will be estimated
migAC: 1 #migration rate between A and C will be estimated
migAD: 1 #migration rate between A and D will be estimated
migBC: 1 #migration rate between B and C will be estimated
migBD: 1 #migration rate between B and D will be estimated
migCD: 1 #migration rate between C and D will be estimated

### Supplementary Tables

**Supplementary Table 1.** Details of sampling locations for the 27 newly sequenced *O. edulis* samples, and one newly sequenced *O. chilensis* sample used in this study. NES= Northern European Seas, SES= Southern European Seas.

| Sample ID | Species | Collection Date | Sampling Location | Population | Region |
| --- | --- | --- | --- | --- | --- |
| BZH_02 | *Ostrea edulis* | 2020 | France: Banc du Roz | North Atlantic | NES |
| BZH_03 | *Ostrea edulis* | 2020 | France: Brittany, Banc du Roz | North Atlantic | NES |
| BZH_04 | *Ostrea edulis* | 2020 | France: Brittany, Banc du Roz | North Atlantic | NES |
| BZH_05 | *Ostrea edulis* | 2020 | France: Brittany, Banc du Roz | North Atlantic | NES |
| BZH_06 | *Ostrea edulis* | 2020 | France: Brittany, Banc du Roz | North Atlantic | NES |
| BZH_07 | *Ostrea edulis* | 2020 | France: Brittany, Banc du Roz | Atlantic | NES |
| BZH_08 | *Ostrea edulis* | 2020 | France: Brittany, Banc du Roz | Atlantic | NES |
| COR_01 | *Ostrea edulis* | 2020 | France: Corsica, Diana Lagoon | Western Mediterranean | SES |
| COR_02 | *Ostrea edulis* | 2020 | France: Corsica, Diana Lagoon | Western Mediterranean | SES |
| COR_03 | *Ostrea edulis* | 2020 | France: Corsica, Diana Lagoon | Western Mediterranean | SES |
| COR_04 | *Ostrea edulis* | 2020 | France: Corsica, Diana Lagoon | Western Mediterranean | SES |
| COR_05 | *Ostrea edulis* | 2020 | France: Corsica, Diana Lagoon | Western Mediterranean | SES |
| COR_06 | *Ostrea edulis* | 2020 | France: Corsica, Diana Lagoon | Western Mediterranean | SES |
| COR_07 | *Ostrea edulis* | 2020 | France: Corsica, Diana Lagoon | Western Mediterranean | SES |
| DEN_01 | *Ostrea edulis* | 2020 | Denmark: Limfjord | North Sea | NES |
| NET_01 | *Ostrea edulis* | 2020 | Netherlands: Lake Grevelingen | North Sea | NES |
| NET_02 | *Ostrea edulis* | 2020 | Netherlands: Lake Grevelingen | North Sea | NES |
| NET_03 | *Ostrea edulis* | 2020 | Netherlands: Lake Grevelingen | North Sea | NES |
| NET_04 | *Ostrea edulis* | 2020 | Netherlands: Lake Grevelingen | North Sea | NES |
| NET_05 | *Ostrea edulis* | 2020 | Netherlands: Lake Grevelingen | North Sea | NES |
| NET_06 | *Ostrea edulis* | 2020 | Netherlands: Lake Grevelingen | North Sea | NES |
| D9_08 | *Ostrea edulis* | 1999 | Crimea: Ozero Donuzlav | Black Sea | SES |
| D9_20 | *Ostrea edulis* | 1999 | Crimea: Ozero Donuzlav | Black Sea | SES |
| S11_01 | *Ostrea edulis* | 1999 | Crimea: Sevastopol | Black Sea | SES |
| S11_12 | *Ostrea edulis* | 1999 | Crimea: Sevastopol | Black Sea | SES |
| S11_22 | *Ostrea edulis* | 1999 | Crimea: Ozero Donuzlav | Black Sea | SES |
| S9_07 | *Ostrea edulis* | 1999 | Crimea: Sevastopol | Black Sea | SES |
| S9_12 | *Ostrea edulis* | 1999 | Crimea: Sevastopol | Black Sea | SES |
| OCME M1B | *Ostrea chilensis* | 2010 | Wales: Menai Strait | Not Applicable-Outgroup | Not Applicable-Outgroup |

**Supplementary Table 2.** Variant categories assigned by SNPeff and the general category assigned for use in this study, as used by Colston-Nepali, Lapègue and Bierne (2026).

| **SNPeff category** | **General category** |
| --- | --- |
| upstream_gene_variant | Intergenic variant |
| downstream_gene_variant | Intergenic variant |
| intergenic_region | Intergenic variant |
| intron_variant | Intron variant |
| 3_prime_UTR_variant | Intron variant |
| 5_prime_UTR_variant | Intron variant |
| 5_prime_UTR_premature_start_codon_gain_variant | Intron variant |
| splice_region_variant&intron_variant | Intron variant |
| splice_acceptor_variant&intron_variant | Intron variant |
| splice_donor_variant&intron_variant | Intron variant |
| synonymous_variant | Synonymous variant |
| stop_retained_variant | Synonymous variant |
| splice_region_variant&synonymous_variant | Synonymous variant |
| splice_region_variant&stop_retained_variant | Synonymous variant |
| missense_variant | Nonsynonymous variant |
| stop_gained | Nonsynonymous variant |
| missense_variant&splice_region_variant | Nonsynonymous variant |
| splice_region_variant | Nonsynonymous variant |
| stop_lost&splice_region_variant | Nonsynonymous variant |
| stop_lost | Nonsynonymous variant |
| start_lost | Nonsynonymous variant |
| stop_gained&splice_region_variant | Nonsynonymous variant |
| initiator_codon_variant | Nonsynonymous variant |
| start_lost&splice_region_variant | Nonsynonymous variant |

**Supplementary Table 3.** Chromosome numbering correspondence between the four available *O. edulis* reference genomes (modified from Colston-Nepali, Lapègue and Bierne 2026). The three chromosomes with large inversions are marked in bold.

| The Qingdao assembly (Li et al. 2023), used in this study) | The Roscoff assembly, (Boutet et al. 2022), used by Lapègue et al. 2022, and Alves Monterio et al. 2024 | The Roslin assembly (Gundappa et al. 2022), used by Sambade et al. 2025 | The Sanger assembly (Adkins and Mrowicki 2022) |
| --- | --- | --- | --- |
| 1 | 1 | 1 | 1 |
| 2 | 2 | 4 | 2 |
| 3 | 3 | 6 | 5 |
| **4** | **4** | **5** | **6** |
| **5** | **5** | **2** | **4** |
| 6 | 6 | 3 | 3 |
| 7 | 7 | 7 | 7 |
| 8 | 9 | 9 | 8 |
| **9** | **8** | **8** | **9** |
| 10 | 10 | 10 | 10 |

**Supplementary Table 4.** GO terms significantly enriched in the inversion on chromosome 4 at p < 0.05. None of these are significant after FDR correction.

| GO Term | Database | ID | Input number | Background number | Enrich Ratio | P-Value | Corrected P-Value |
| --- | --- | --- | --- | --- | --- | --- | --- |
| Steroid biosynthesis | KEGG PATHWAY | crg00100 | 1 | 10 | 0.1 | 0.07780755773 | 0.4720945069 |
| Phenylalanine metabolism | KEGG PATHWAY | crg00360 | 2 | 23 | 0.08695652174 | 0.01438741379 | 0.4720945069 |

**Supplementary Table 5.** GO terms significantly enriched in the inversion on chromosome 5 at p < 0.05. Seven of these are significant after FDR correction (indicated with a *)

| GO Term | Database | ID | Input number | Background number | Enrich Ratio | P-Value | Corrected P-Value |
| --- | --- | --- | --- | --- | --- | --- | --- |
| Metabolic pathways* | KEGG PATHWAY | crg01100 | 57 | 1509 | 3.78E-02 | 6.17E-06 | 0.0002861919009 |
| Amino sugar and nucleotide sugar metabolism* | KEGG PATHWAY | crg00520 | 11 | 95 | 1.16E-01 | 7.73E-06 | 0.0002861919009 |
| Drug metabolism - other enzymes* | KEGG PATHWAY | crg00983 | 7 | 77 | 9.09E-02 | 0.001318124675 | 0.02696007659 |
| RNA trNESport* | KEGG PATHWAY | crg03013 | 10 | 158 | 6.33E-02 | 0.001839080196 | 0.02696007659 |
| Neuroactive ligand-receptor interaction* | KEGG PATHWAY | crg04080 | 14 | 276 | 5.07E-02 | 0.001904244834 | 0.02696007659 |
| Ubiquitin mediated proteolysis* | KEGG PATHWAY | crg04120 | 10 | 162 | 6.17E-02 | 0.002185952156 | 0.02696007659 |
| Fatty acid metabolism* | KEGG PATHWAY | crg01212 | 6 | 70 | 8.57E-02 | 0.003783012072 | 0.03999184191 |
| Pyrimidine metabolism | KEGG PATHWAY | crg00240 | 5 | 57 | 8.77E-02 | 0.007333435387 | 0.06783427733 |
| N-Glycan biosynthesis | KEGG PATHWAY | crg00510 | 5 | 59 | 8.47E-02 | 0.008370787635 | 0.06882647611 |
| Fatty acid biosynthesis | KEGG PATHWAY | crg00061 | 3 | 23 | 1.30E-01 | 0.01390915685 | 0.1029277607 |
| Glycine, serine and threonine metabolism | KEGG PATHWAY | crg00260 | 4 | 46 | 8.70E-02 | 0.01649346068 | 0.1065752605 |
| Nicotinate and nicotinamide metabolism | KEGG PATHWAY | crg00760 | 3 | 26 | 1.15E-01 | 0.01872268089 | 0.1065752605 |
| Fatty acid elongation | KEGG PATHWAY | crg00062 | 3 | 26 | 1.15E-01 | 0.01872268089 | 0.1065752605 |
| Ribosome biogenesis in eukaryotes | KEGG PATHWAY | crg03008 | 5 | 78 | 6.41E-02 | 0.02362297325 | 0.1248642872 |
| RNA degradation | KEGG PATHWAY | crg03018 | 5 | 85 | 5.88E-02 | 0.0320481636 | 0.1581042738 |
| Spliceosome | KEGG PATHWAY | crg03040 | 7 | 150 | 4.67E-02 | 0.03550632766 | 0.1642167654 |
| Endocytosis | KEGG PATHWAY | crg04144 | 8 | 191 | 4.19E-02 | 0.04308938375 | 0.1875655528 |
| Glycosaminoglycan biosynthesis - heparan sulfate / heparin | KEGG PATHWAY | crg00534 | 2 | 16 | 1.25E-01 | 0.04764192582 | 0.1958612506 |

**Supplementary Table 6.** GO terms significantly enriched in the inversion on chromosome 9 at p < 0.05. Two of these are significant after FDR correction (indicated with a *)

| Term | Database | ID | Input number | Background number | Enrich Ratio | P-Value | Corrected P-Value |
| --- | --- | --- | --- | --- | --- | --- | --- |
| Pentose and glucuronate interconversions* | KEGG PATHWAY | crg00040 | 5 | 20 | 0.25 | 0.001096318297 | 0.03291721958 |
| Metabolic pathways* | KEGG PATHWAY | crg01100 | 74 | 1509 | 0.04903909874 | 0.001097240653 | 0.03291721958 |
| Starch and sucrose metabolism | KEGG PATHWAY | crg00500 | 6 | 38 | 0.1578947368 | 0.002817501437 | 0.06339378234 |
| Galactose metabolism | KEGG PATHWAY | crg00052 | 6 | 41 | 0.1463414634 | 0.003948421036 | 0.07107157864 |
| Phagosome | KEGG PATHWAY | crg04145 | 10 | 127 | 0.07874015748 | 0.01396522294 | 0.1819699548 |
| Arginine and proline metabolism | KEGG PATHWAY | crg00330 | 7 | 72 | 0.09722222222 | 0.0141532187 | 0.1819699548 |
| Glycolysis / Gluconeogenesis | KEGG PATHWAY | crg00010 | 6 | 59 | 0.1016949153 | 0.01864582128 | 0.2097654894 |
| Autophagy - animal | KEGG PATHWAY | crg04140 | 8 | 103 | 0.07766990291 | 0.02811165392 | 0.2811165392 |
| Fructose and mannose metabolism | KEGG PATHWAY | crg00051 | 4 | 39 | 0.1025641026 | 0.04992186115 | 0.4151226232 |

**Supplementary Table 7.**
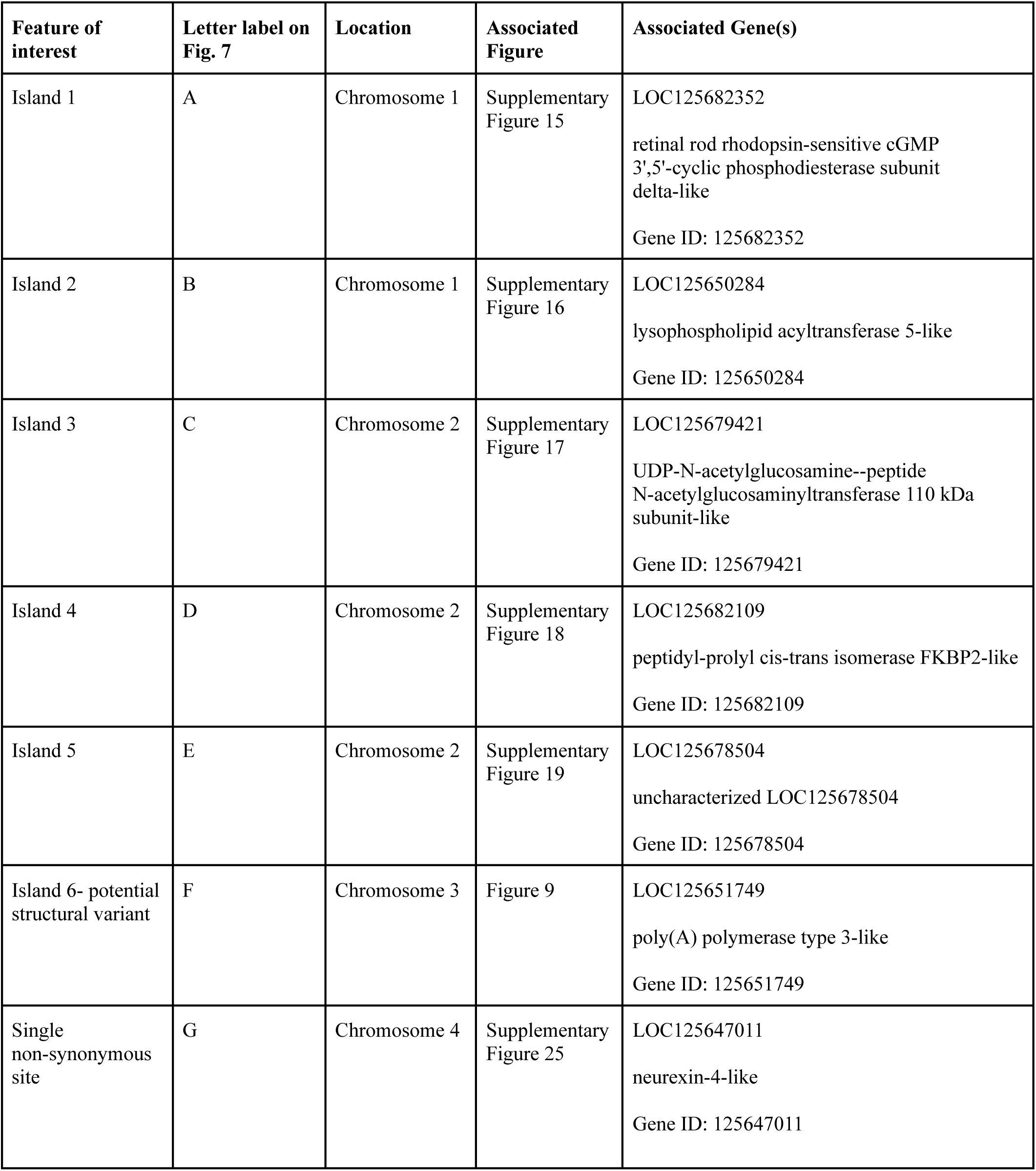
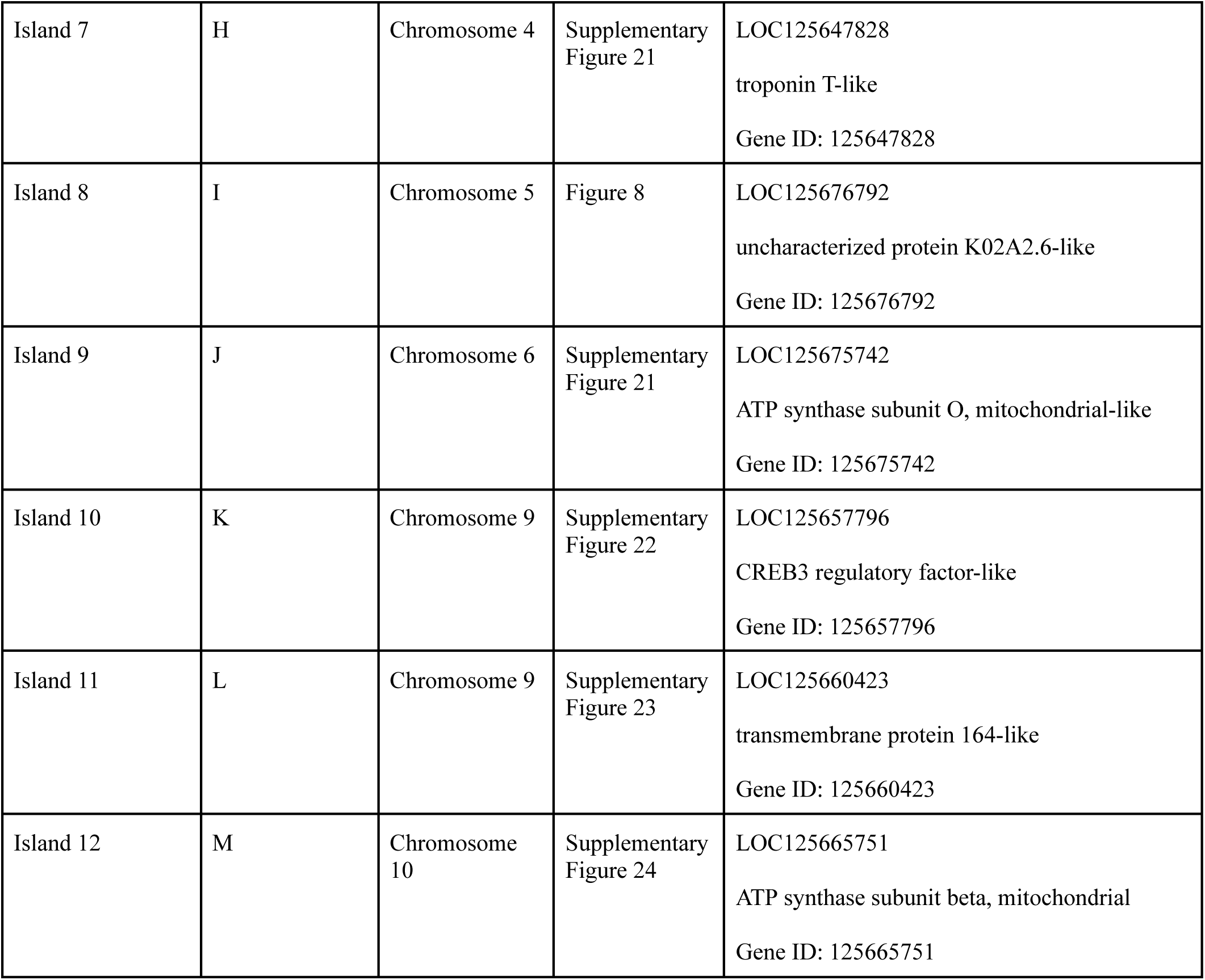
Features of interest identified by the Π scan (islands, potential structural variant, and single nonsynonymous site), corresponding label on Fig. 7, their location, associated figure and potential associated gene(s).

| Feature of interest | Letter label on Fig. 7 | Location | Associated Figure | Associated Gene(s) |
| --- | --- | --- | --- | --- |
| Island 1 | A | Chromosome 1 | Supplementary Figure 15 | LOC125682352<br>retinal rod rhodopsin-sensitive cGMP 3',5'-cyclic phosphodiesterase subunit delta-like<br>Gene ID: 125682352 |
| Island 2 | B | Chromosome 1 | Supplementary Figure 16 | LOC125650284<br>lysophospholipid acyltransferase 5-like<br>Gene ID: 125650284 |
| Island 3 | C | Chromosome 2 | Supplementary Figure 17 | LOC125679421<br>UDP-N-acetylglucosamine--peptide N-acetylglucosaminyltransferase 110 kDa subunit-like<br>Gene ID: 125679421 |
| Island 4 | D | Chromosome 2 | Supplementary Figure 18 | LOC125682109<br>peptidyl-prolyl cis-trans isomerase FKBP2-like<br>Gene ID: 125682109 |
| Island 5 | E | Chromosome 2 | Supplementary Figure 19 | LOC125678504<br>uncharacterized LOC125678504<br>Gene ID: 125678504 |
| Island 6- potential structural variant | F | Chromosome 3 | Figure 9 | LOC125651749<br>poly(A) polymerase type 3-like<br>Gene ID: 125651749 |
| Single non-synonymous site | G | Chromosome 4 | Supplementary Figure 25 | LOC125647011<br>neurexin-4-like<br>Gene ID: 125647011 |
| Island 7 | H | Chromosome 4 | Supplementary<br>Figure 21 | LOC125647828<br>troponin T-like<br>Gene ID: 125647828 |
| Island 8 | I | Chromosome 5 | Figure 8 | LOC125676792<br>uncharacterized protein K02A2.6-like<br>Gene ID: 125676792 |
| Island 9 | J | Chromosome 6 | Supplementary<br>Figure 21 | LOC125675742<br>ATP synthase subunit O, mitochondrial-like<br>Gene ID: 125675742 |
| Island 10 | K | Chromosome 9 | Supplementary<br>Figure 22 | LOC125657796<br>CREB3 regulatory factor-like<br>Gene ID: 125657796 |
| Island 11 | L | Chromosome 9 | Supplementary<br>Figure 23 | LOC125660423<br>transmembrane protein 164-like<br>Gene ID: 125660423 |
| Island 12 | M | Chromosome<br>10 | Supplementary<br>Figure 24 | LOC125665751<br>ATP synthase subunit beta, mitochondrial<br>Gene ID: 125665751 |

**Supplementary Table 8.**
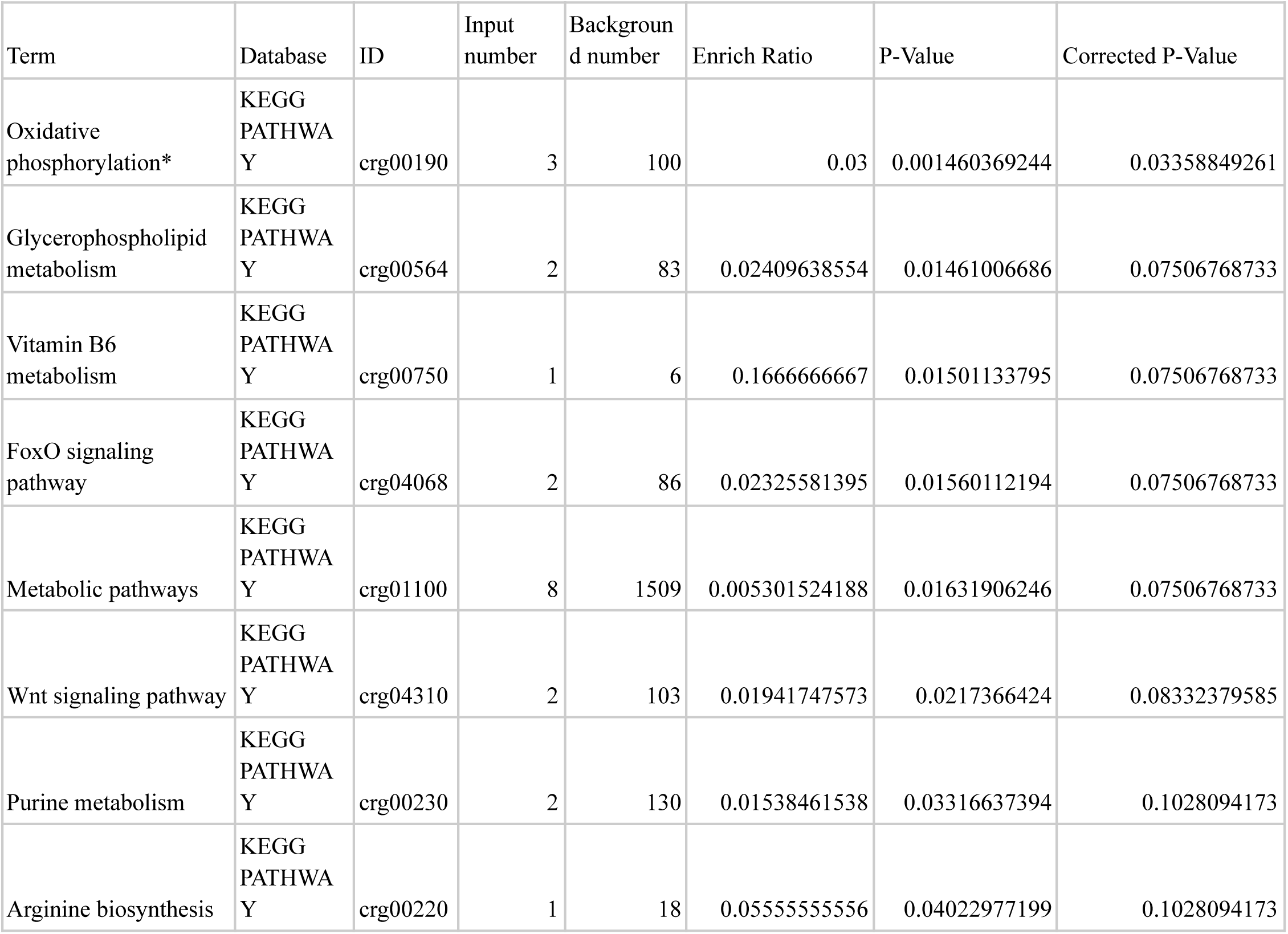
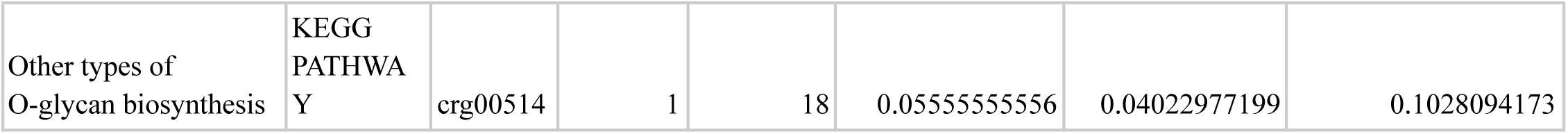
GO terms significantly enriched in the islands at p < 0.05. One of these is significant after FDR correction (indicated with a *)

**Supplementary Table 9.**
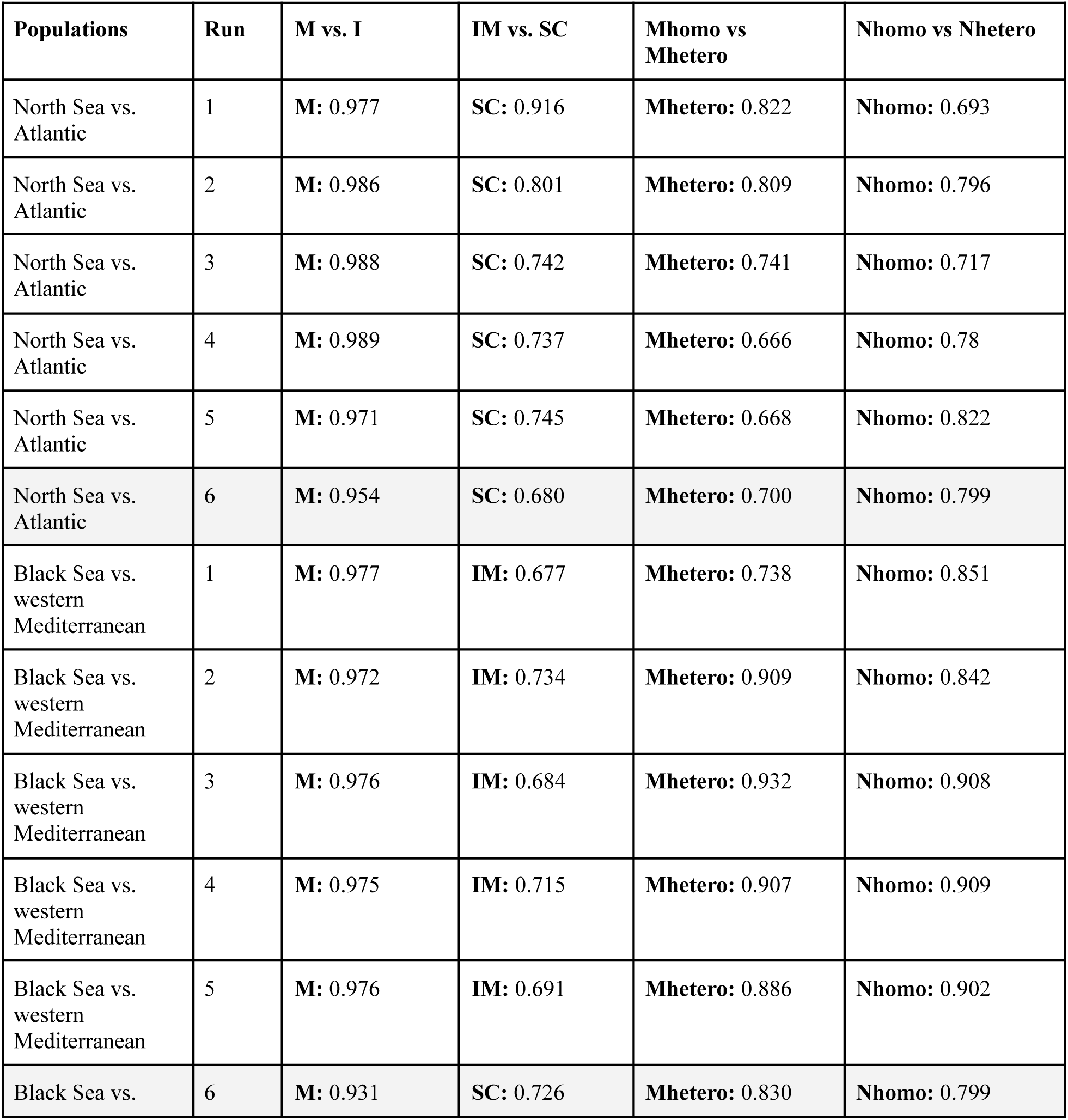
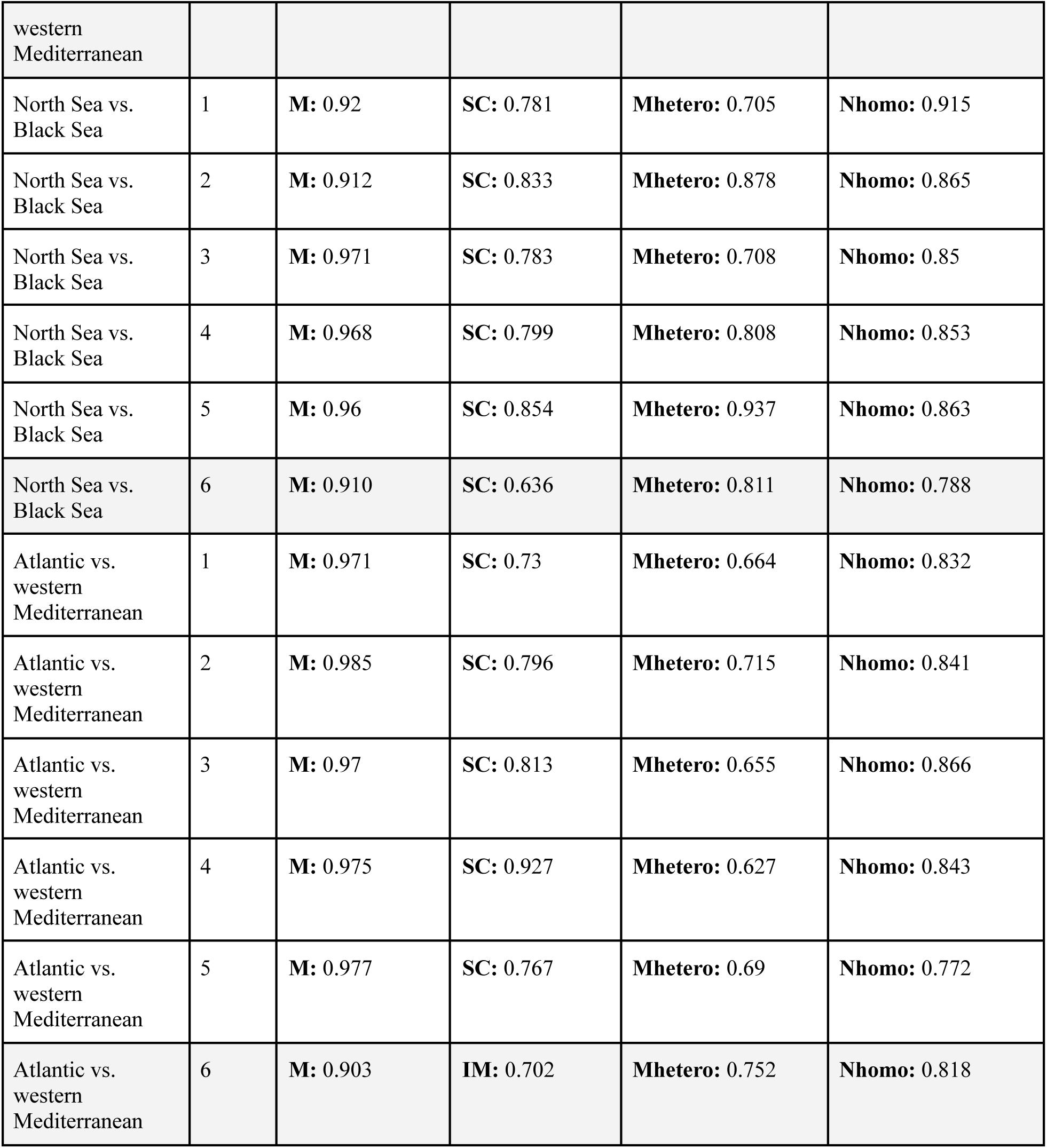
Demographic inferences with DILS: model comparisons for the two-population models. Four pairwise population comparisons were performed in six replicates: i) North Sea vs. Atlantic, ii) Black Sea vs. western Mediterranean, iii) North Sea vs. Black Sea, iv) Atlantic vs. western Mediterranean. “Run” is the replicated run number. Posterior probabilities from hierarchical model testing are shown for each run: i) “M vs. I (current migration vs. current isolation), ii) “IM vs SC” (isolation with continuous migration vs. secondary contact), iii) “Mhomo vs. Mhetero” (homogenous vs heterogenous effective migration across the genome), and iv) “Nhomo vs. Nhetero” (homogenous vs heterogenous effective size across the genome).

**Supplementary Table 10.** Demographic inferences with DILS for the four-population models. “Run” is the replicated run number. Posterior probability from topology testing with random forest is shown for each run. “topo1” is (A,((B,C),D)) and it corresponds to geography. “topo2” is (A,(B,(C, D))) and it corresponds to the mitochondrial tree. “topo3” is ((A, B),(C, D)) and it corresponds to the genetic lineages. A is the Black Sea, B is the North Sea, C is the Atlantic and D is the western Mediterranean.

| **Populations** | **Run** | **topo1 vs. topo2 vs. topo3** |
| --- | --- | --- |
| North Sea vs. Atlantic vs.<br>western Mediterranean vs. Black Sea | 1 | **topo3:** 0.54 |
| North Sea vs. Atlantic vs.<br>western Mediterranean vs. Black Sea | 2 | **topo2:** 0.50 |
| North Sea vs. Atlantic vs.<br>western Mediterranean vs. Black Sea | 3 | **topo3:** 0.50 |

### Supplementary Figures

**Supplementary Figure 1.**
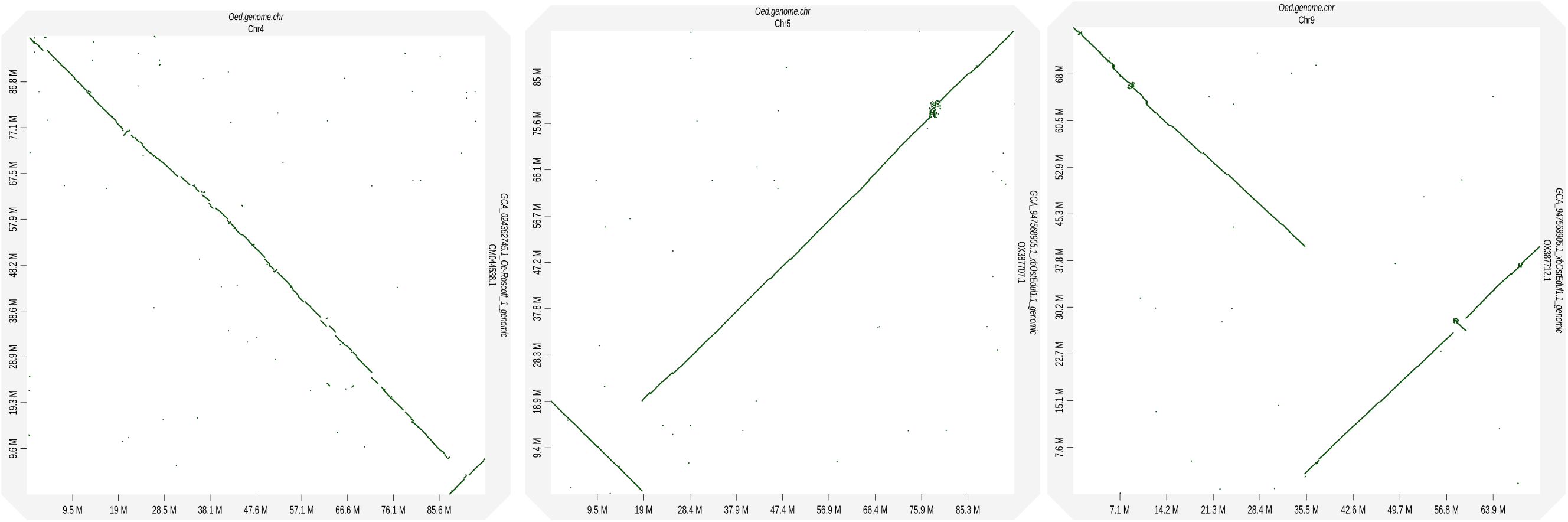
D-GENIES results for chromosomes 4, 5, and 9. For chromosome 4: the Qingdao assembly on the *x* axis and the Roslin assembly on the *y* axis. For chromosome 5: the Qingdao assembly on the *x* axis and the Sanger assembly on the *y* axis. For chromosome 9: the Qingdao assembly on the *x* axis and the Sanger assembly on the *y* axis.

**Supplementary Figure 2.**
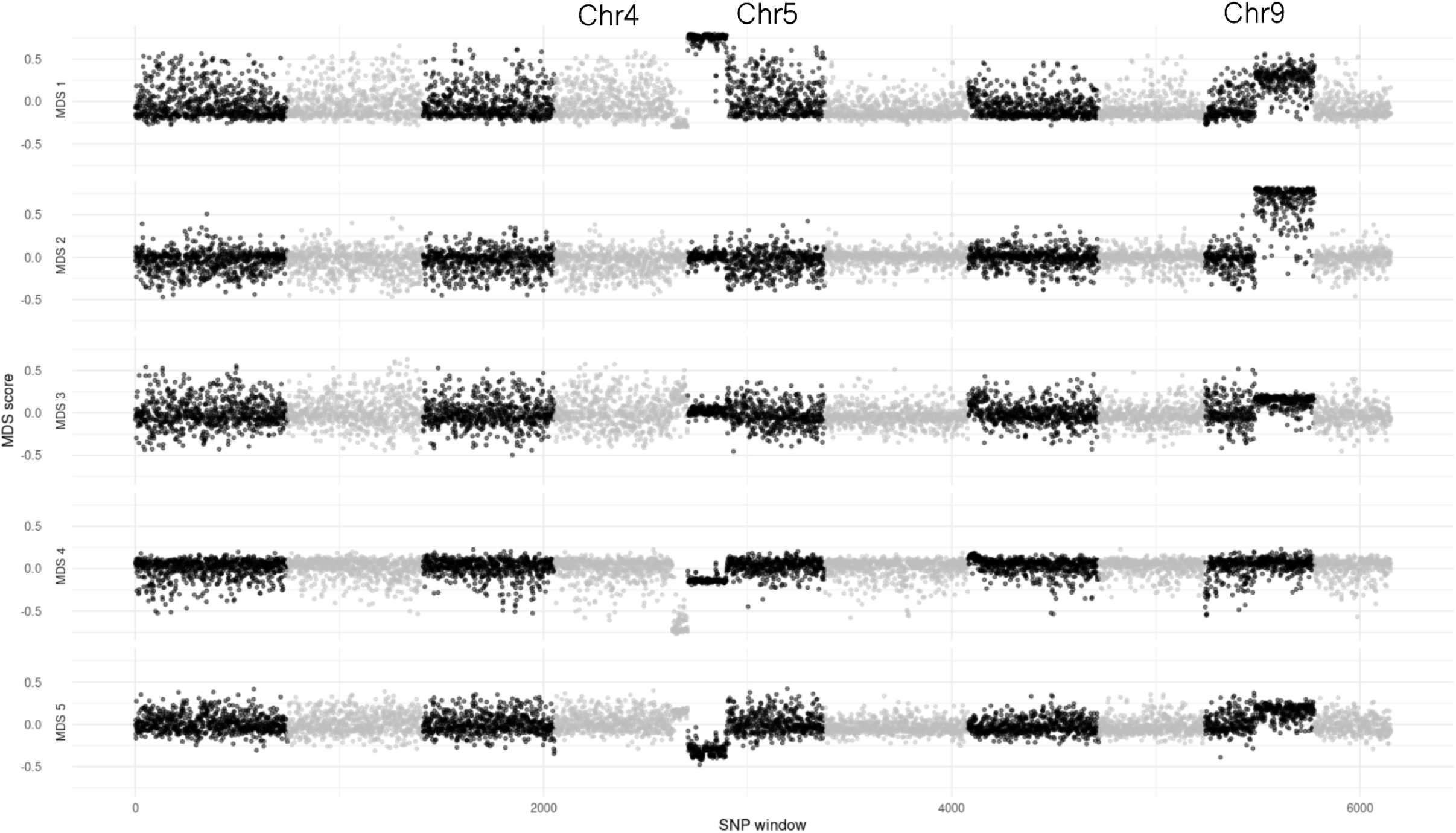
Multi-dimensional scaling plot of the first five principal components of a local principal component analysis of the whole flat oyster genome. Chromosomes are differentiated by alternating grey or black coloured points. Chromosomes that have inversions (4, 5, and 9) are marked. The presence of a structural variant on chromosome 4 can be seen on MDS 1, 3, and 5, on chromosome 5 on MDS 1, 4, and 5, and on chromosome 9 on MDS 1, 2, and 3.

**Supplementary Figure 3.**
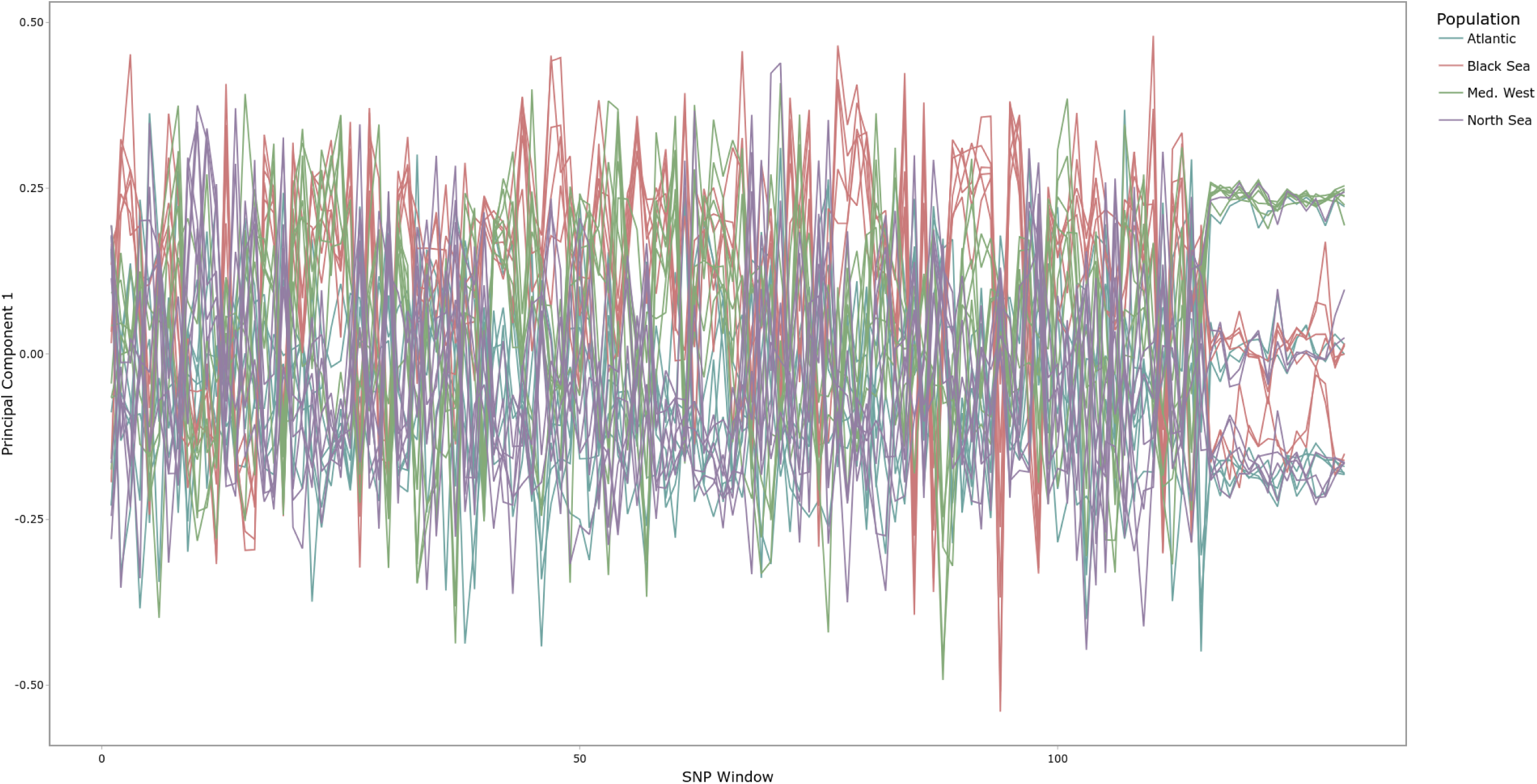
Local PCA for chromosome 4. Scores for the first principal component are plotted on the x-axis. Corresponding SNP windows of 5kb are indicated on the y-axis. Colours indicate the population of the individual (Blue: Atlantic, Purple: North Sea, Green: western Mediterranean, Red: Black Sea).

**Supplementary Figure 4.**
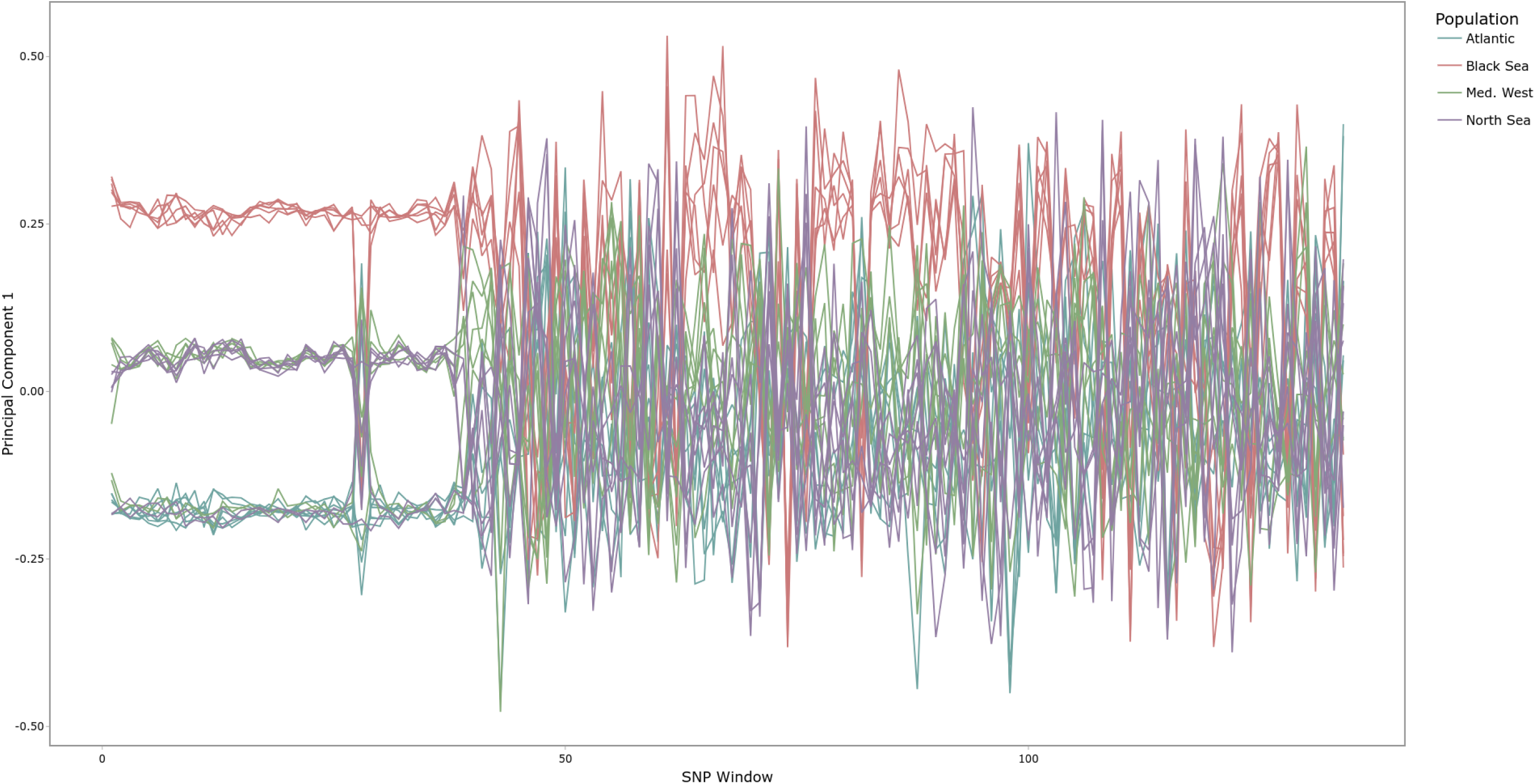
Local PCA for chromosome 5. Scores for the first principal component are plotted on the x-axis. Corresponding SNP windows of 5kb are indicated on the y-axis. Colours indicate the population of the individual (Blue: Atlantic, Purple: North Sea, Green: western Mediterranean, Red: Black Sea).

**Supplementary Figure 5.**
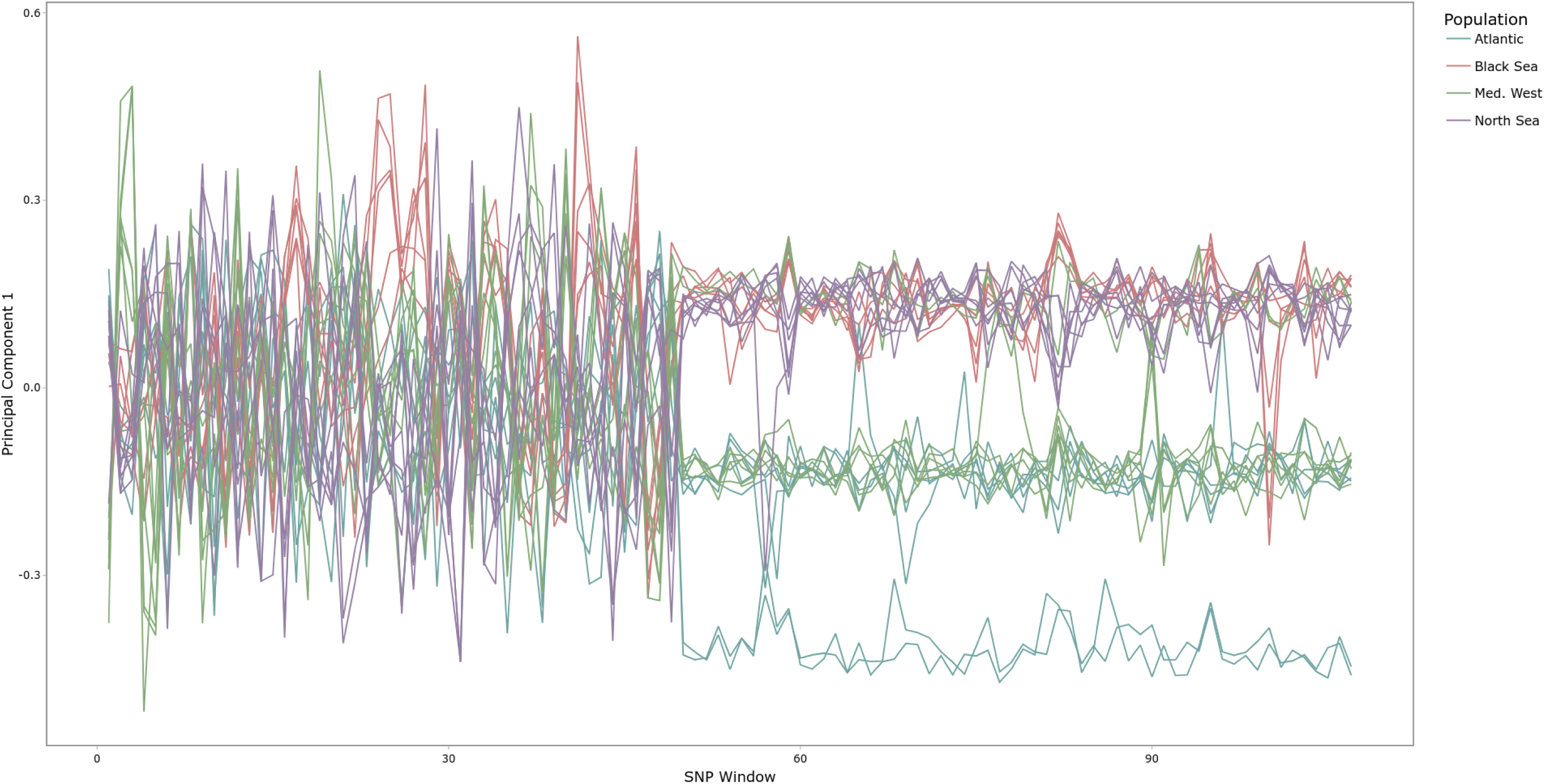
Local PCA for chromosome 9. Scores for the first principal component are plotted on the x-axis. Corresponding SNP windows of 5kb are indicated on the y-axis. Colours indicate the population of the individual (Blue: Atlantic, Purple: North Sea, Green: western Mediterranean, Red: Black Sea).

**Supplementary Figure 6.**
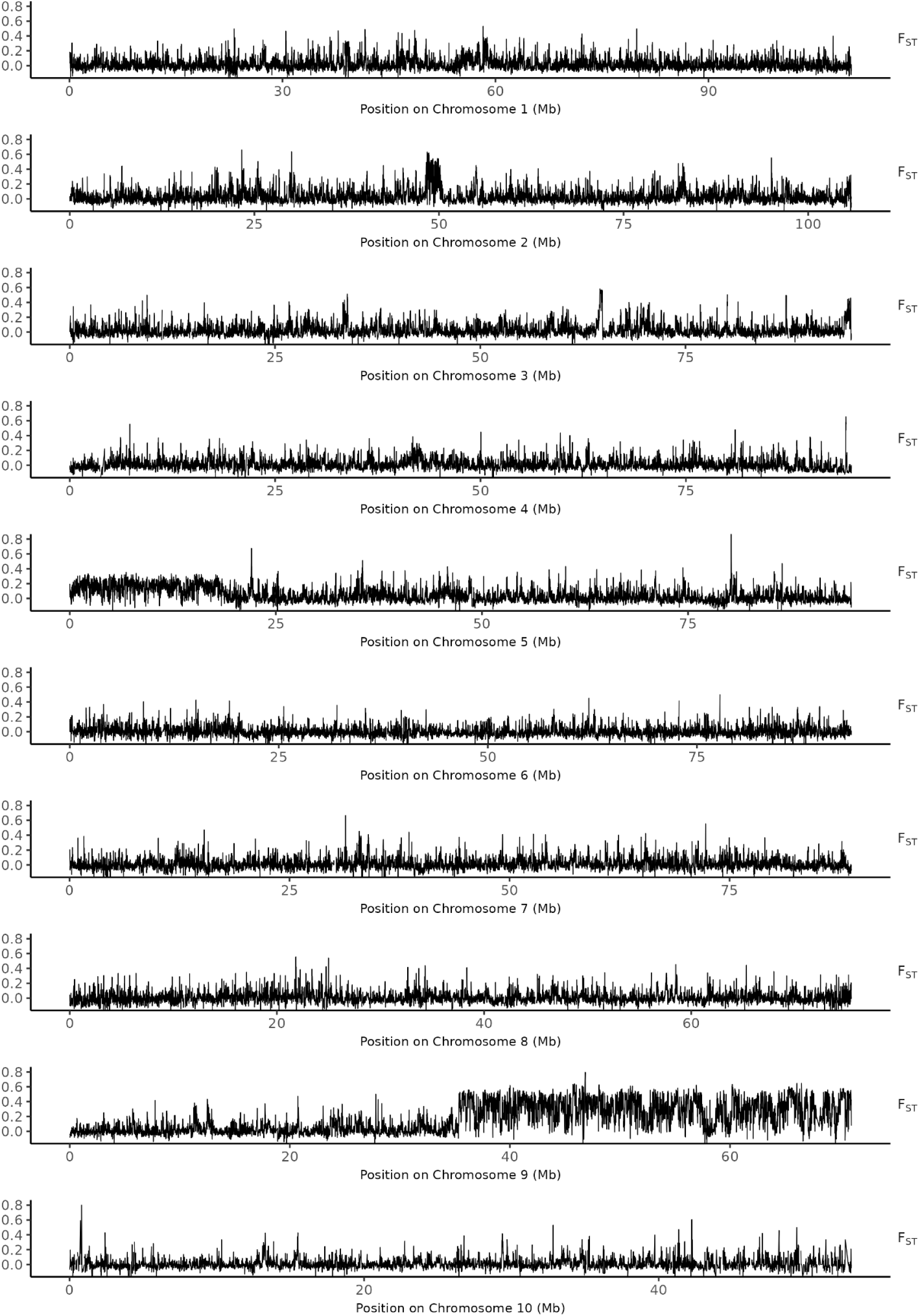
Genomewide pairwise F_ST_ scan between the Atlantic and North Sea populations

**Supplementary Figure 7.**
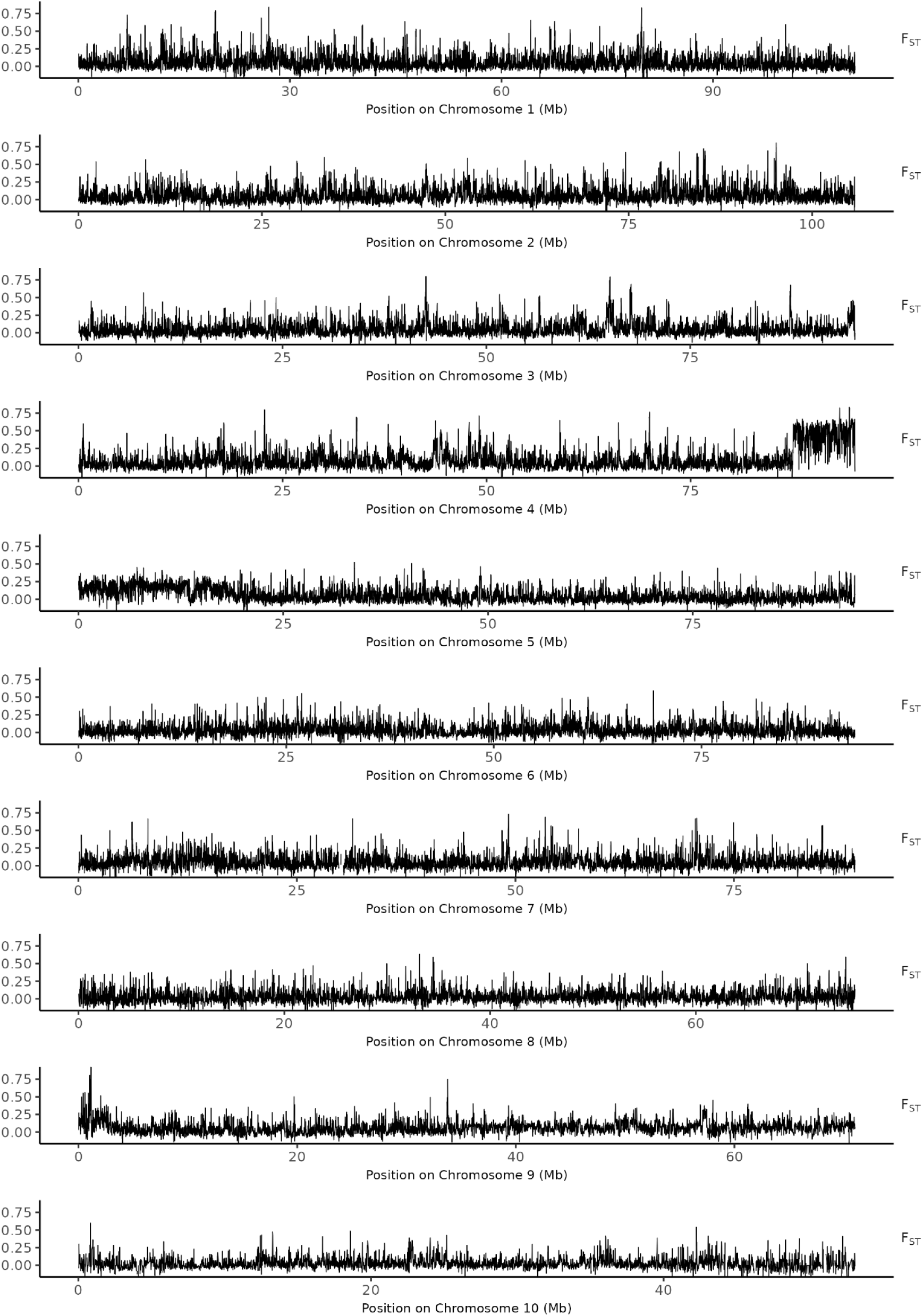
Genomewide pairwise F_ST_ scan between the Atlantic and western Mediterranean populations

**Supplementary Figure 8.**
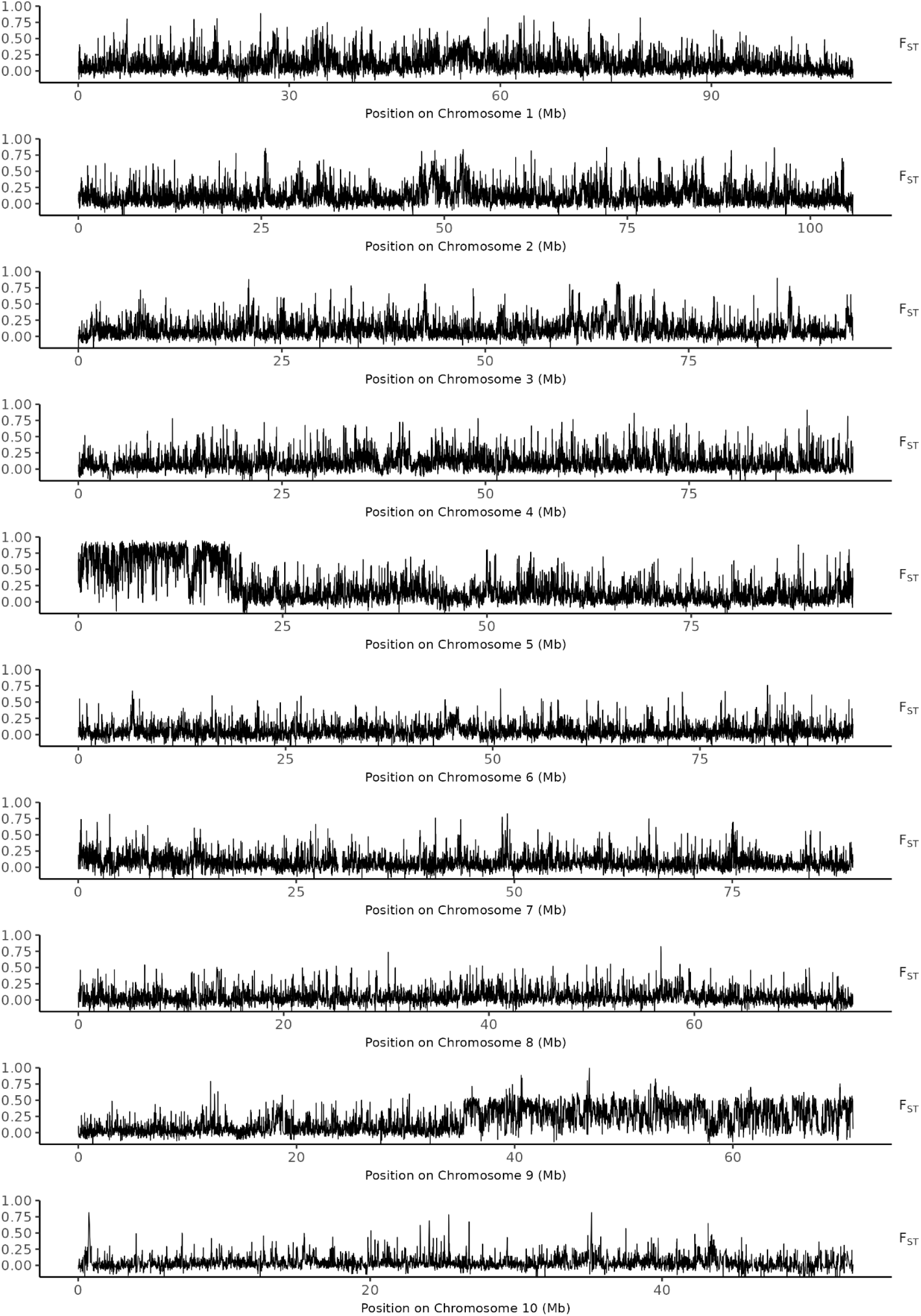
Genomewide pairwise F_ST_ scan between the Atlantic and Black Sea populations

**Supplementary Figure 9.**
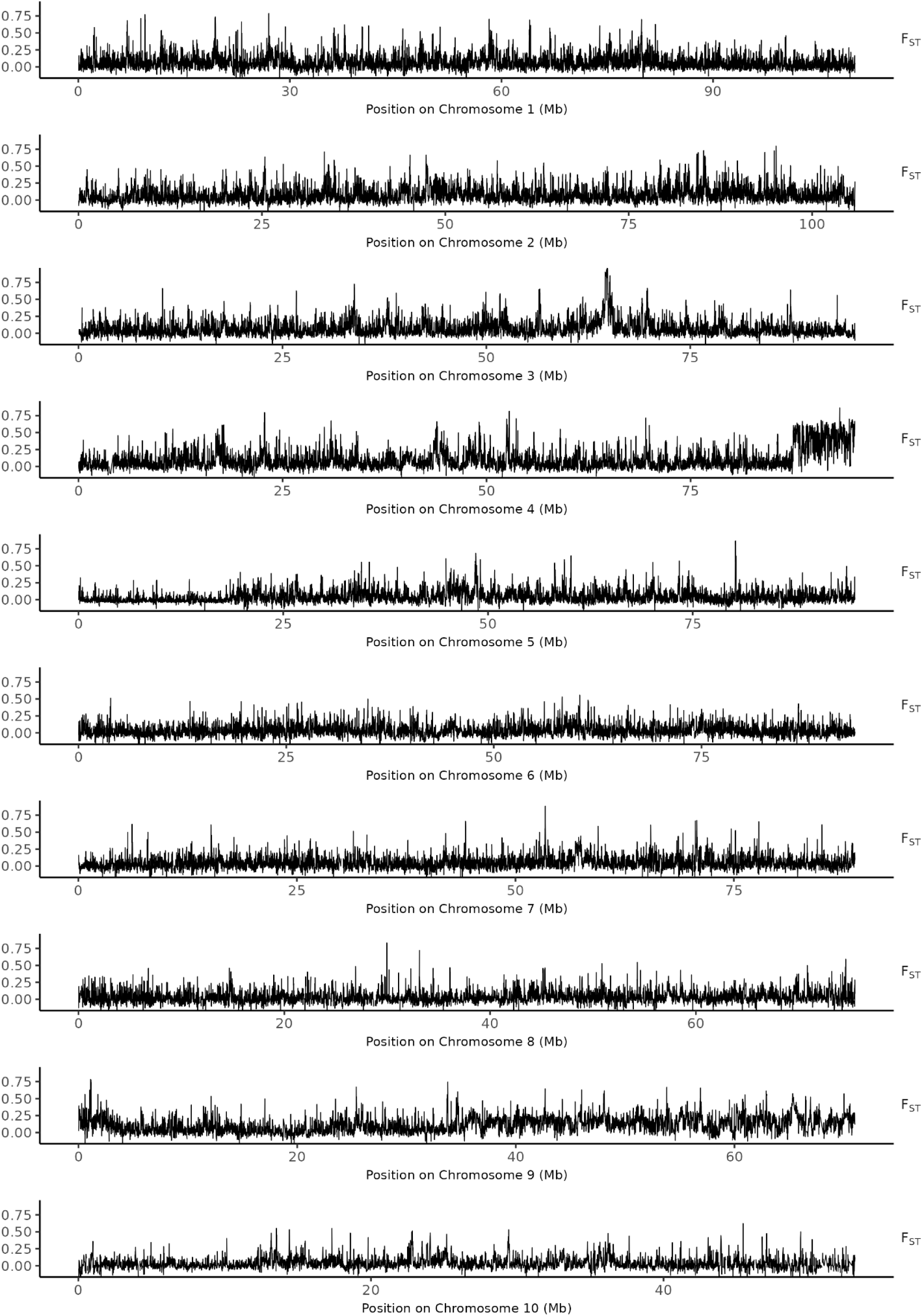
Genomewide pairwise F_ST_ scan between the North Sea and western Mediterranean populations

**Supplementary Figure 10.**
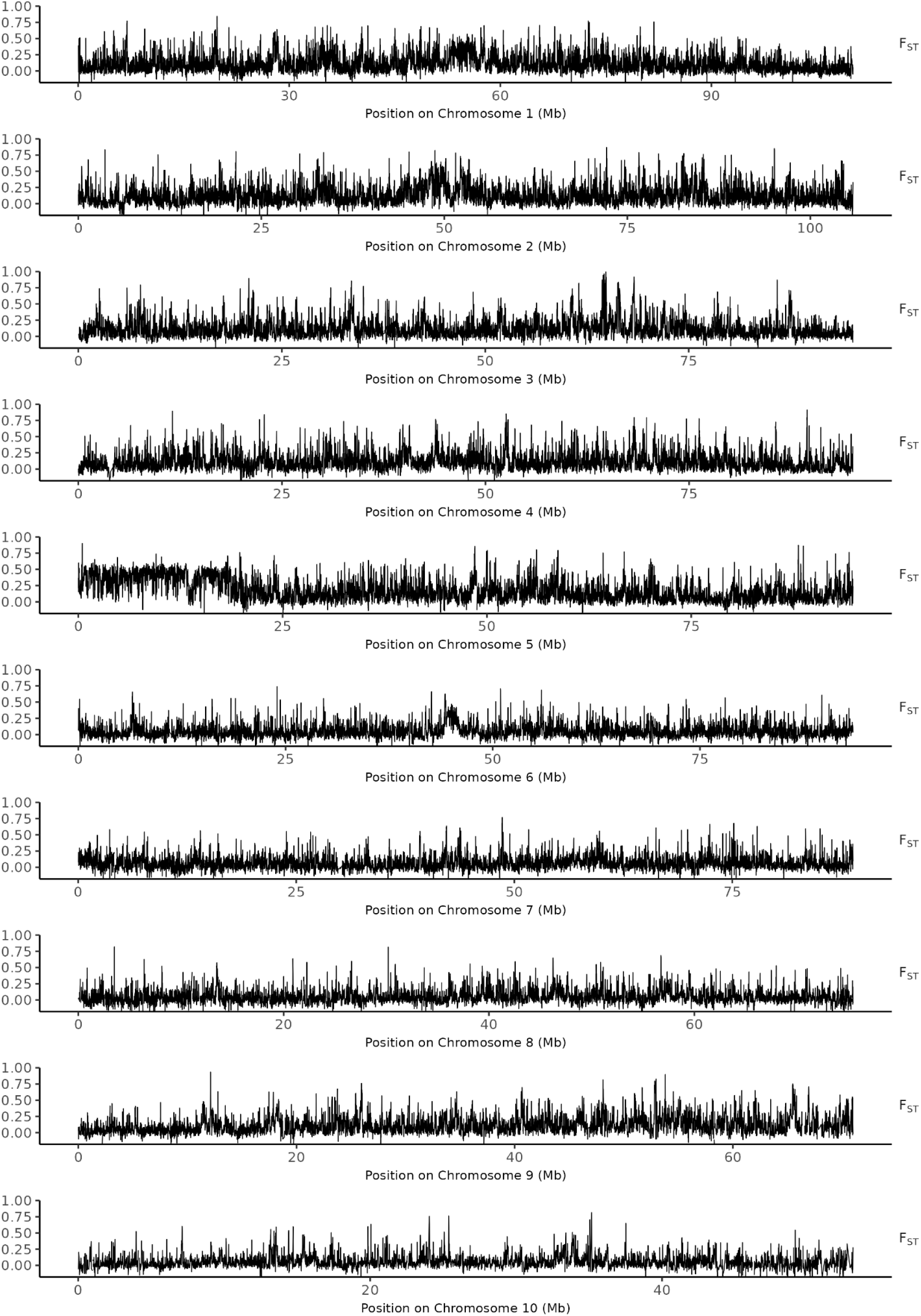
Genomewide pairwise F_ST_ scan between the North Sea and Black Sea populations

**Supplementary Figure 11.**
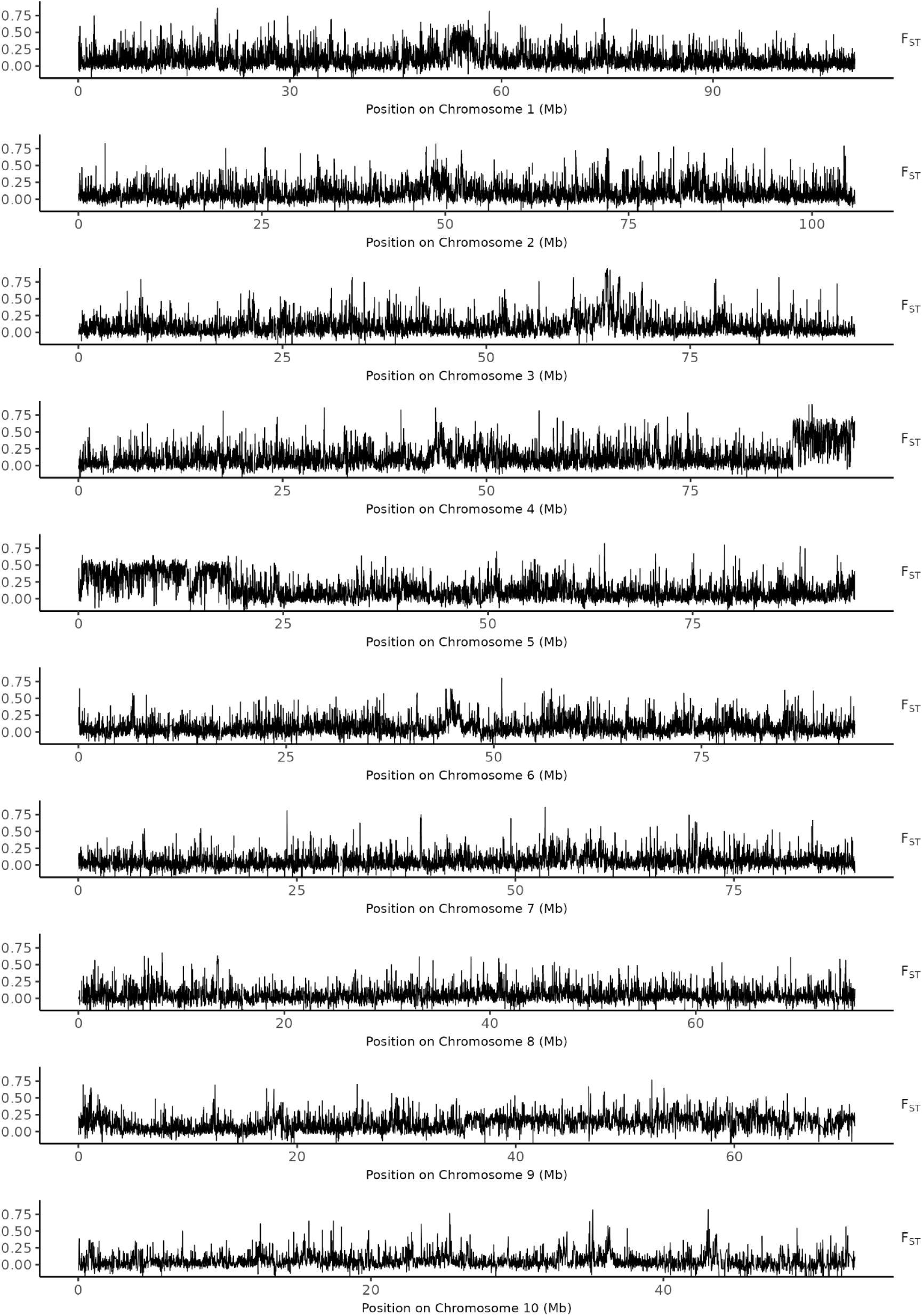
Genomewide pairwise F_ST_ scan between the western Mediterranean and Black Sea

**Supplementary Figure 12.**
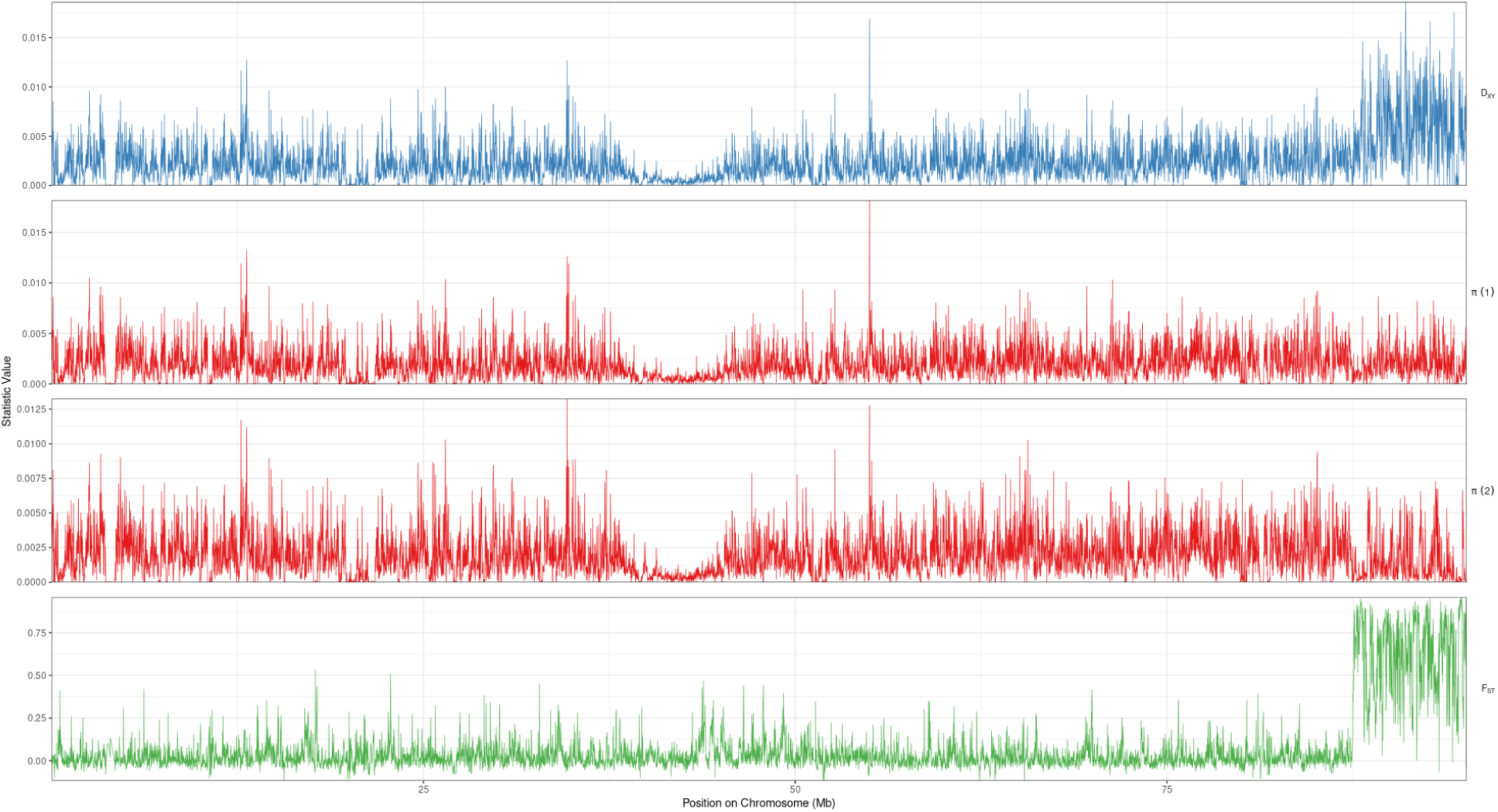
Pairwise F_ST_ (green), and pairwise Dxy (blue) between inversion alleles, and pi for each inversion allele (red) across chromosome 4.

**Supplementary Figure 13.**
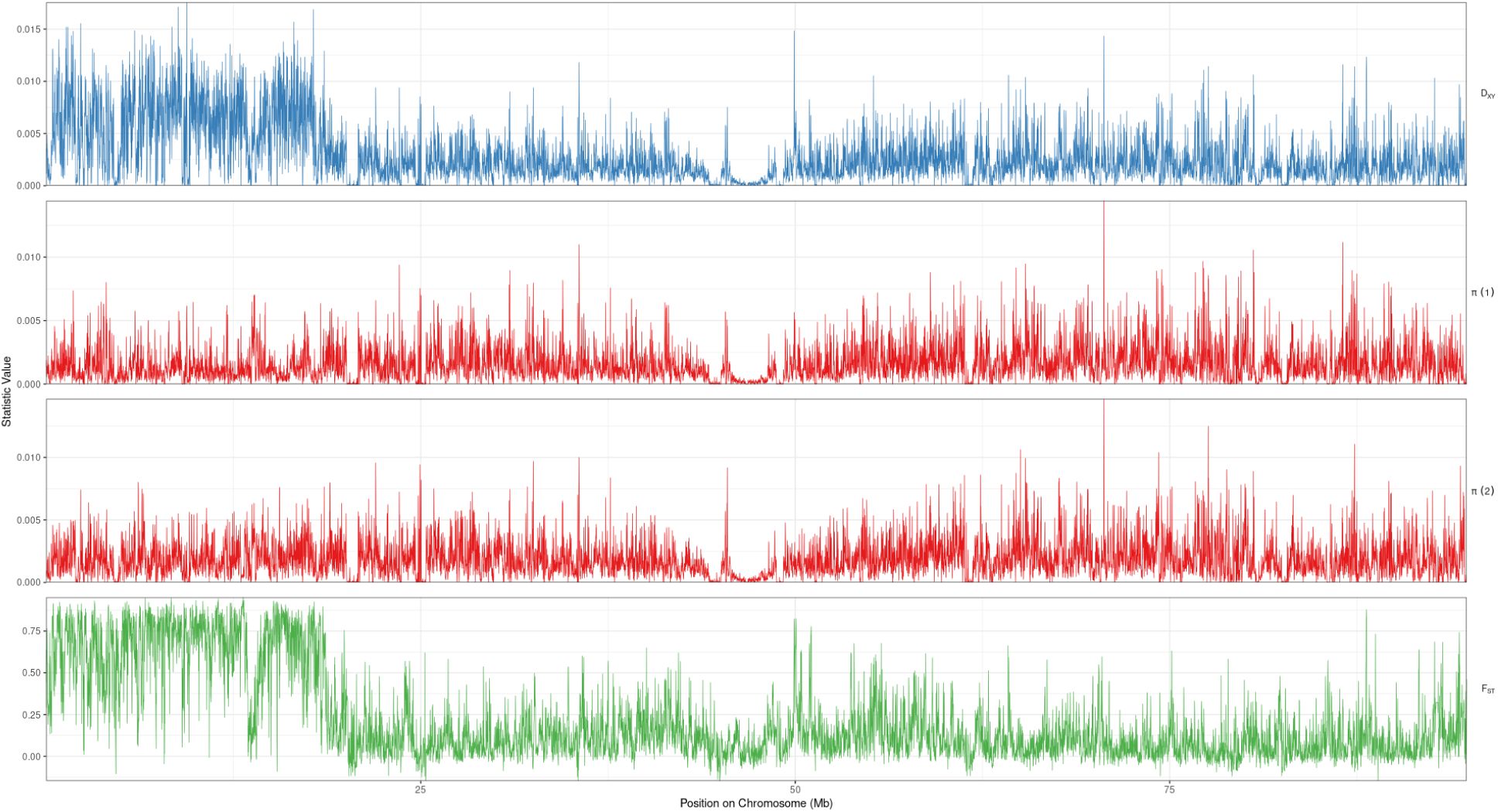
Pairwise F_ST_ (green), and pairwise Dxy (blue) between inversion alleles, and pi for each inversion allele (red) across chromosome 5.

**Supplementary Figure 14.**
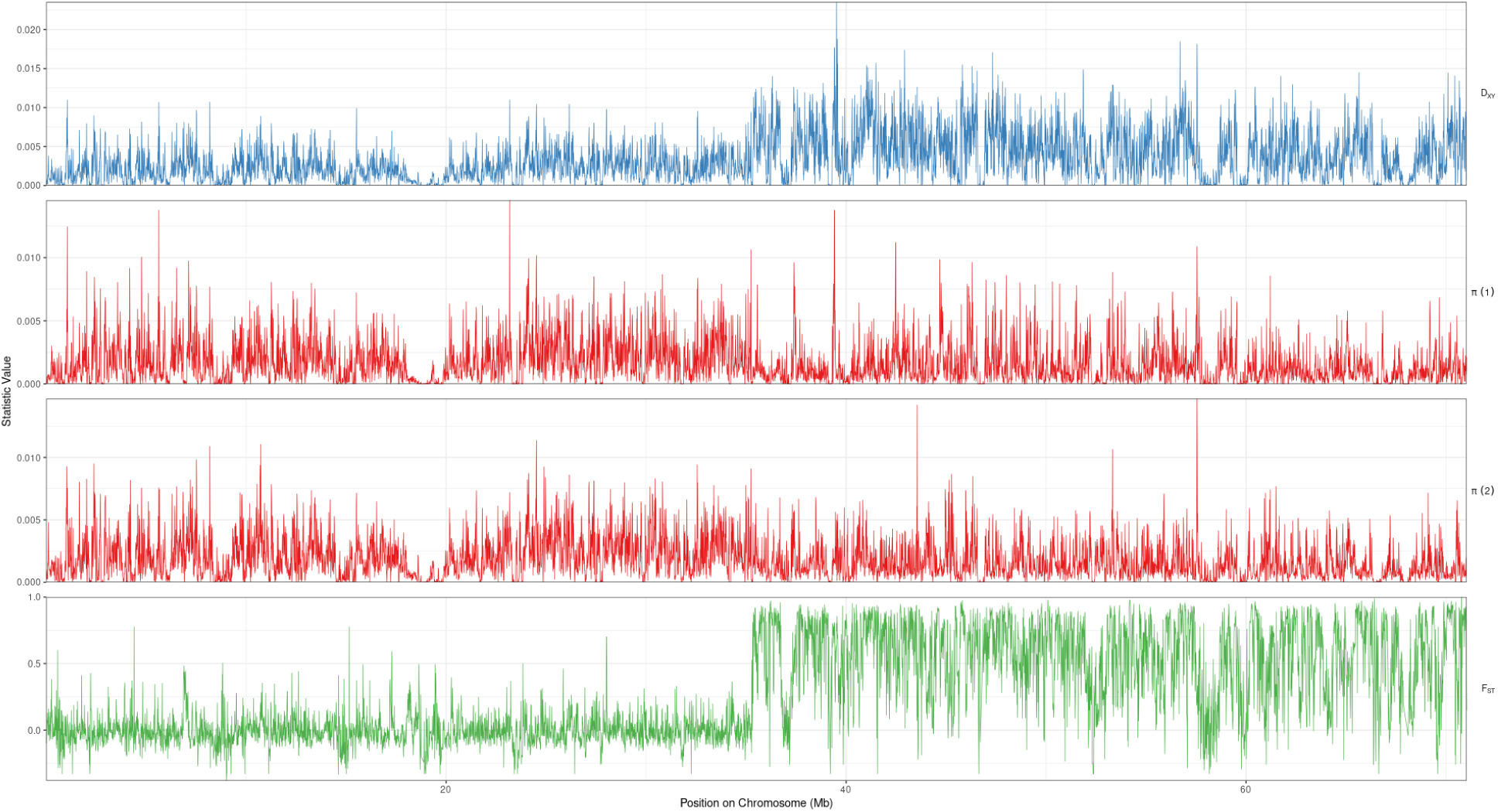
Pairwise F_ST_ (green), and pairwise Dxy (blue) between inversion alleles, and pi for each inversion allele (red) across chromosome 9.

**Supplementary Figure 15.**
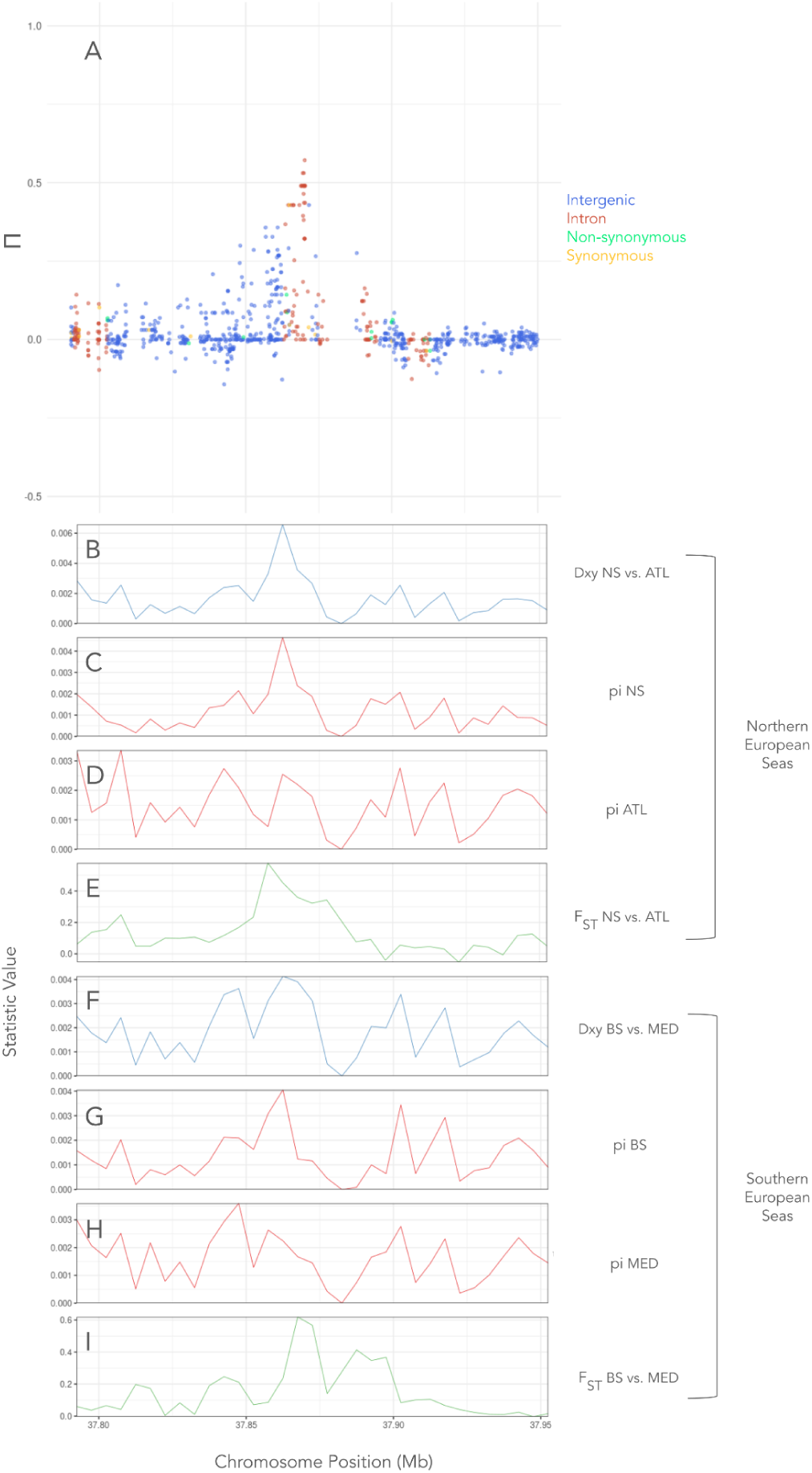
Island of parallelism located on chromosome 1 (Feature A, Island 1). **A** Plot of Π scan with chromosomal position along the x-axis and Π on the y-axis. Points refer to individual SNPs and colours refer to the category of SNP determined by SNPeff (intergenic, intronic, nonsynonymous or synonymous). **B** Dxy calculated in the NES region between the North Sea and Atlantic populations across the island region. **C** pi calculated for the North Sea population across the island region. **D** pi calculated for the Atlantic population across the island region. **E** Pairwise F_ST_ calculated in the NES region between the North Sea and Atlantic populations across the island region. **F** Dxy calculated in the SES region between the Black Sea and western Mediterranean populations across the island region. **G** pi calculated for the Black Sea population across the island region. **H** pi calculated for the western Mediterranean population across the island region. **I** Pairwise F_ST_ calculated in the SES region between the Black Sea and Mediterranean populations across the island region.

**Supplementary Figure 16.**
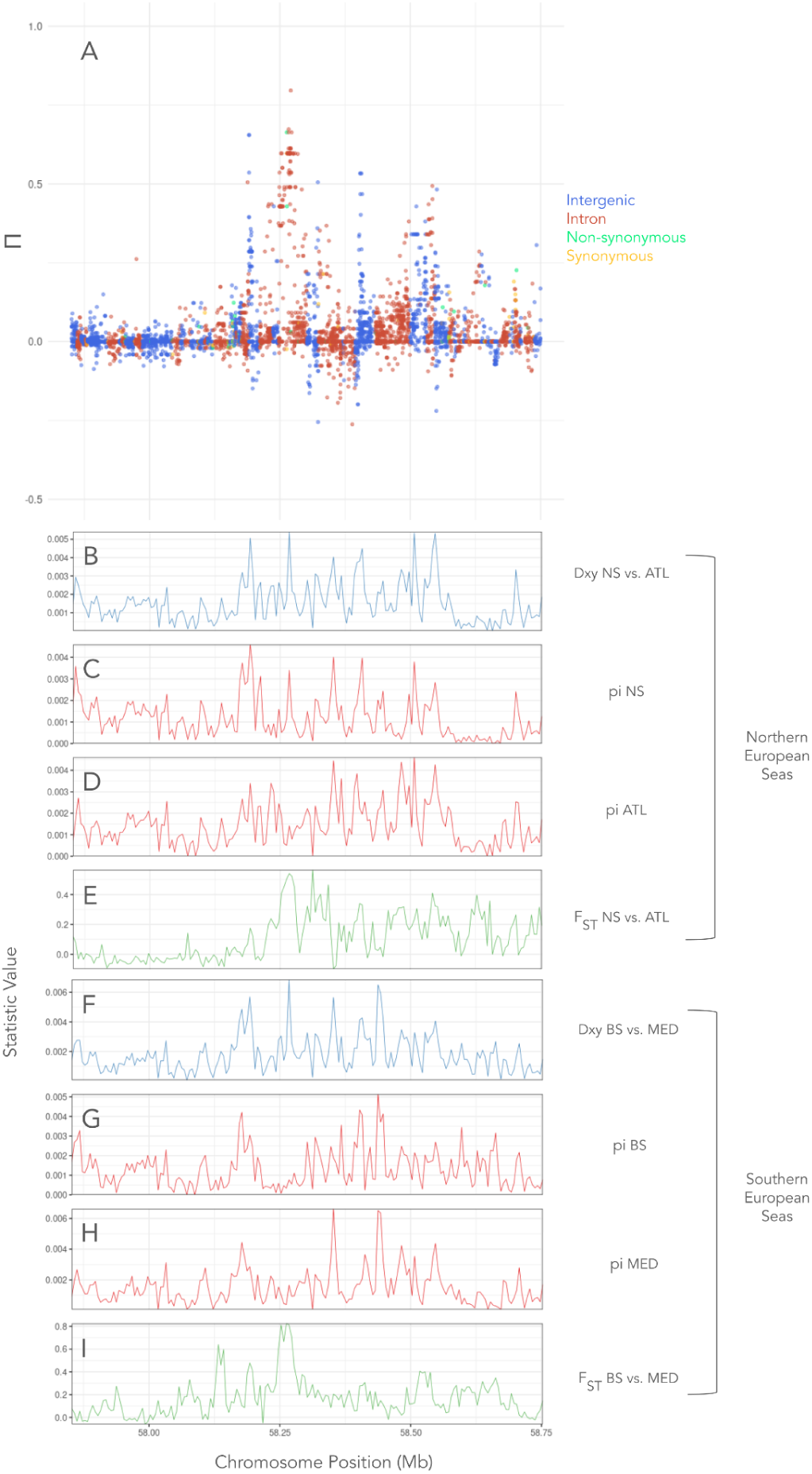
Island of parallelism located on chromosome 1 (Feature B, Island 2). **A** Plot of Π scan with chromosomal position along the x-axis and Π on the y-axis. Points refer to individual SNPs and colours refer to the category of SNP determined by SNPeff (intergenic, intronic, nonsynonymous or synonymous). **B** Dxy calculated in the NES region between the North Sea and Atlantic populations across the island region. **C** pi calculated for the North Sea population across the island region. **D** pi calculated for the Atlantic population across the island region. **E** Pairwise F_ST_ calculated in the NES region between the North Sea and Atlantic populations across the island region. **F** Dxy calculated in the SES region between the Black Sea and western Mediterranean populations across the island region. **G** pi calculated for the Black Sea population across the island region. **H** pi calculated for the western Mediterranean population across the island region. **I** Pairwise F_ST_ calculated in the SES region between the Black Sea and Mediterranean populations across the island region.

**Supplementary Figure 17.**
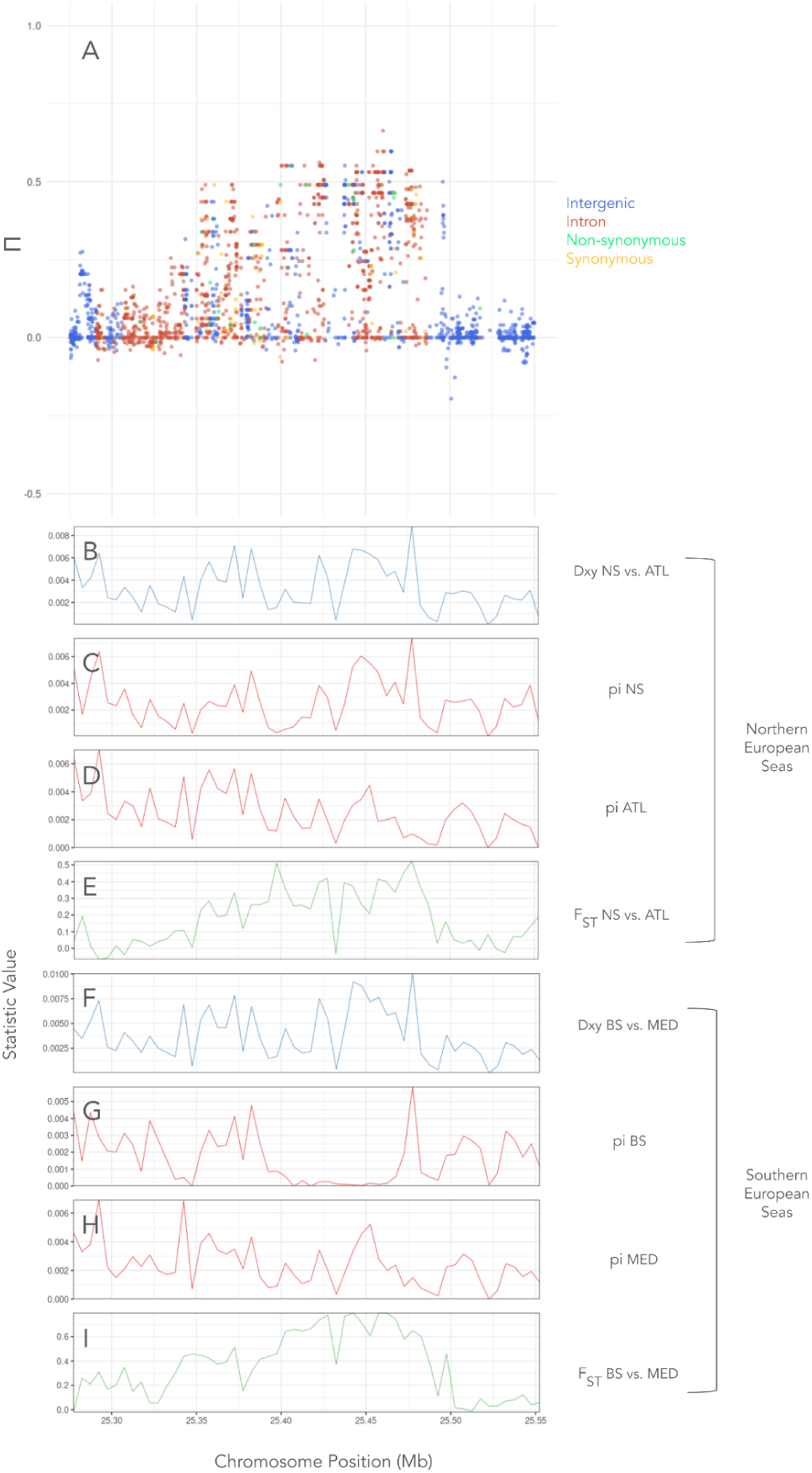
Island of parallelism located on chromosome 2 (Feature C, Island 3). **A** Plot of Π scan with chromosomal position along the x-axis and Π on the y-axis. Points refer to individual SNPs and colours refer to the category of SNP determined by SNPeff (intergenic, intronic, nonsynonymous or synonymous). **B** Dxy calculated in the NES region between the North Sea and Atlantic populations across the island region. **C** pi calculated for the North Sea population across the island region. **D** pi calculated for the Atlantic population across the island region. **E** Pairwise F_ST_ calculated in the NES region between the North Sea and Atlantic populations across the island region. **F** Dxy calculated in the SES region between the Black Sea and western Mediterranean populations across the island region. **G** pi calculated for the Black Sea population across the island region. **H** pi calculated for the western Mediterranean population across the island region. **I** Pairwise F_ST_ calculated in the SES region between the Black Sea and Mediterranean populations across the island region.

**Supplementary Figure 18.**
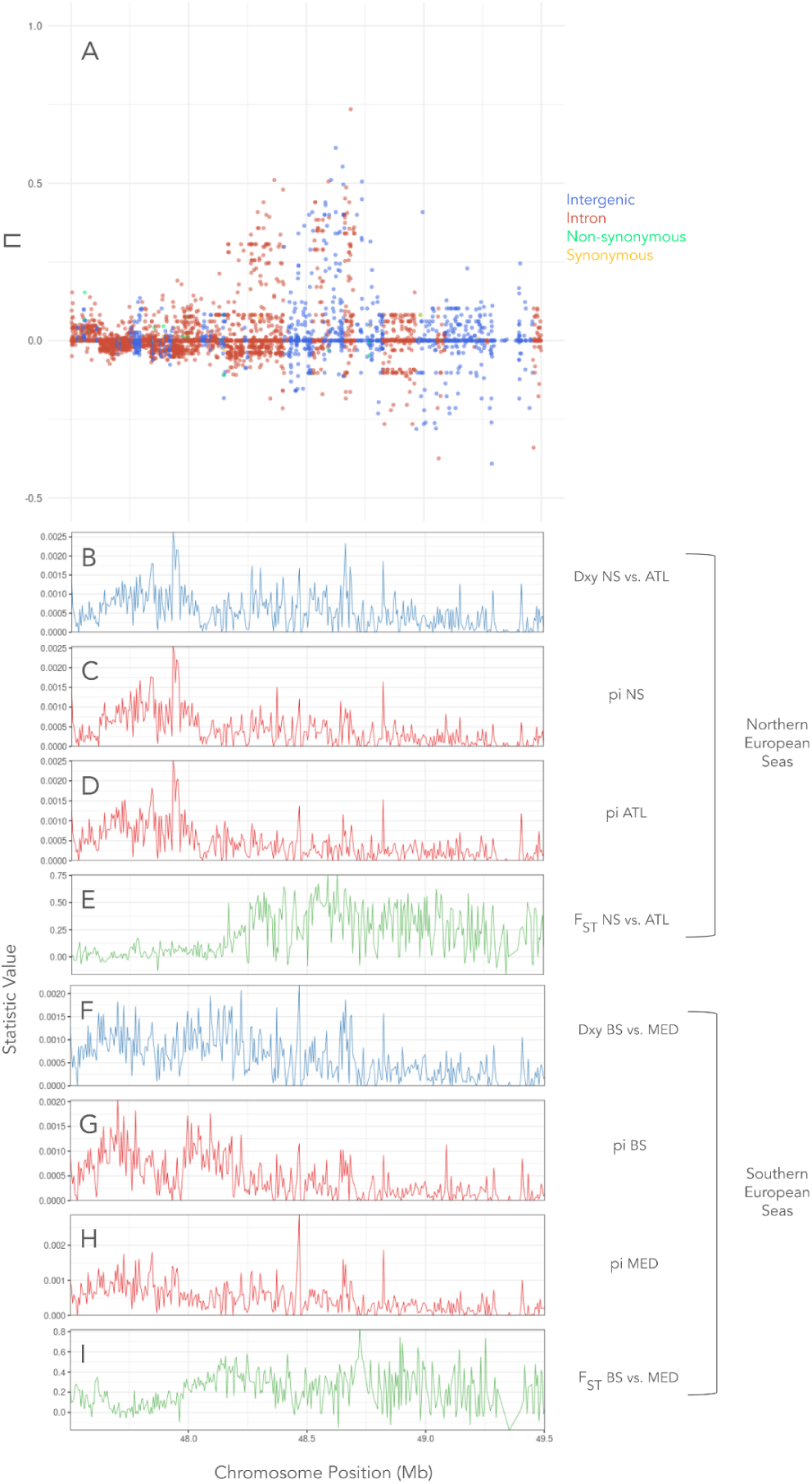
Island of parallelism located on chromosome 2 (Feature D, Island 4). **A** Plot of Π scan with chromosomal position along the x-axis and Π on the y-axis. Points refer to individual SNPs and colours refer to the category of SNP determined by SNPeff (intergenic, intronic, nonsynonymous or synonymous). **B** Dxy calculated in the NES region between the North Sea and Atlantic populations across the island region. **C** pi calculated for the North Sea population across the island region. **D** pi calculated for the Atlantic population across the island region. **E** Pairwise F_ST_ calculated in the NES region between the North Sea and Atlantic populations across the island region. **F** Dxy calculated in the SES region between the Black Sea and western Mediterranean populations across the island region. **G** pi calculated for the Black Sea population across the island region. **H** pi calculated for the western Mediterranean population across the island region. **I** Pairwise F_ST_ calculated in the SES region between the Black Sea and Mediterranean populations across the island region.

**Supplementary Figure 19.**
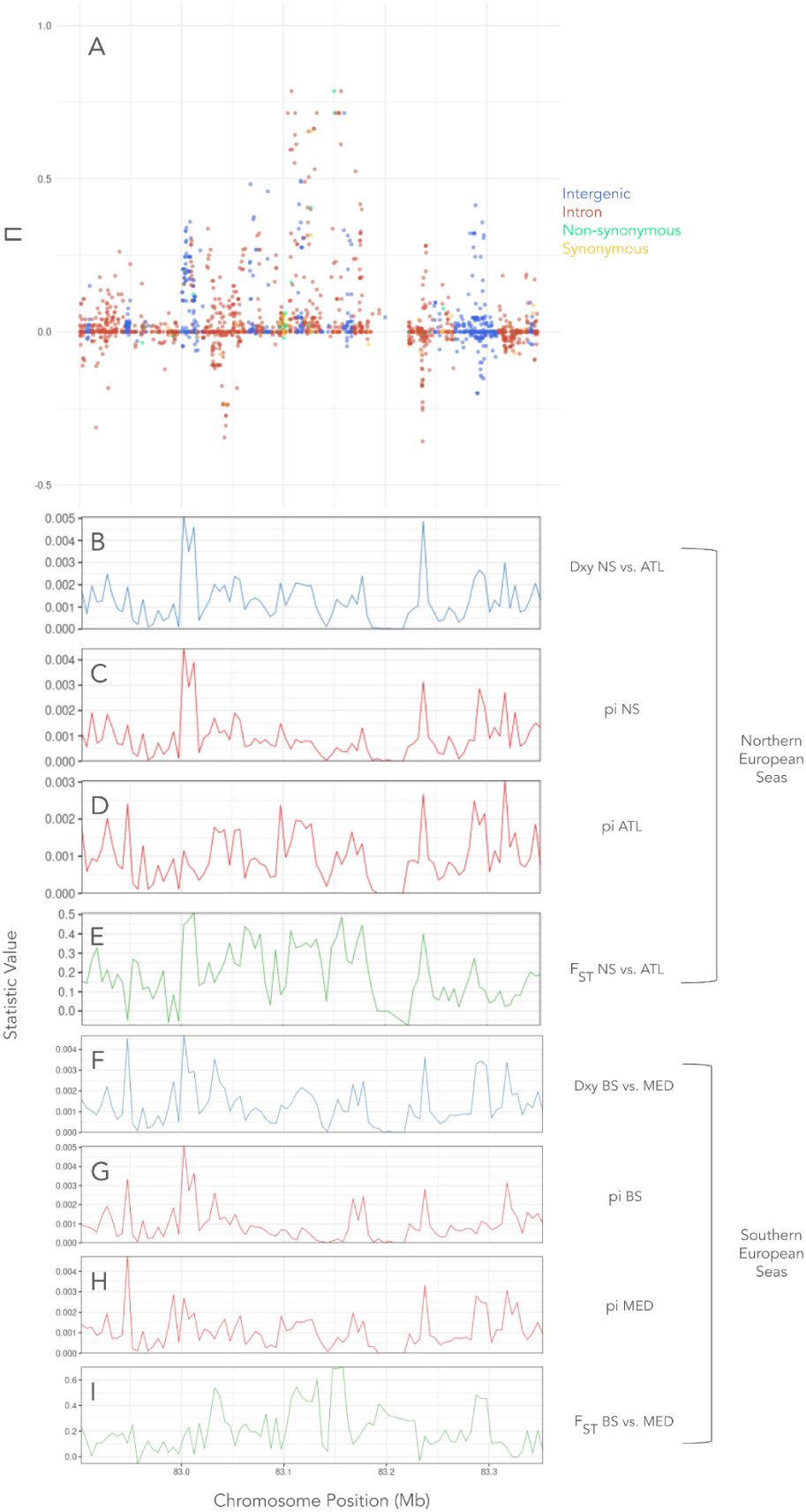
Island of parallelism located on chromosome 2 (Feature E, Island 5). **A** Plot of Π scan with chromosomal position along the x-axis and Π on the y-axis. Points refer to individual SNPs and colours refer to the category of SNP determined by SNPeff (intergenic, intronic, nonsynonymous or synonymous). **B** Dxy calculated in the NES region between the North Sea and Atlantic populations across the island region. **C** pi calculated for the North Sea population across the island region. **D** pi calculated for the Atlantic population across the island region. **E** Pairwise F_ST_ calculated in the NES region between the North Sea and Atlantic populations across the island region. **F** Dxy calculated in the SES region between the Black Sea and western Mediterranean populations across the island region. **G** pi calculated for the Black Sea population across the island region. **H** pi calculated for the western Mediterranean population across the island region. **I** Pairwise F_ST_ calculated in the SES region between the Black Sea and Mediterranean populations across the island region.

**Supplementary Figure 20.**
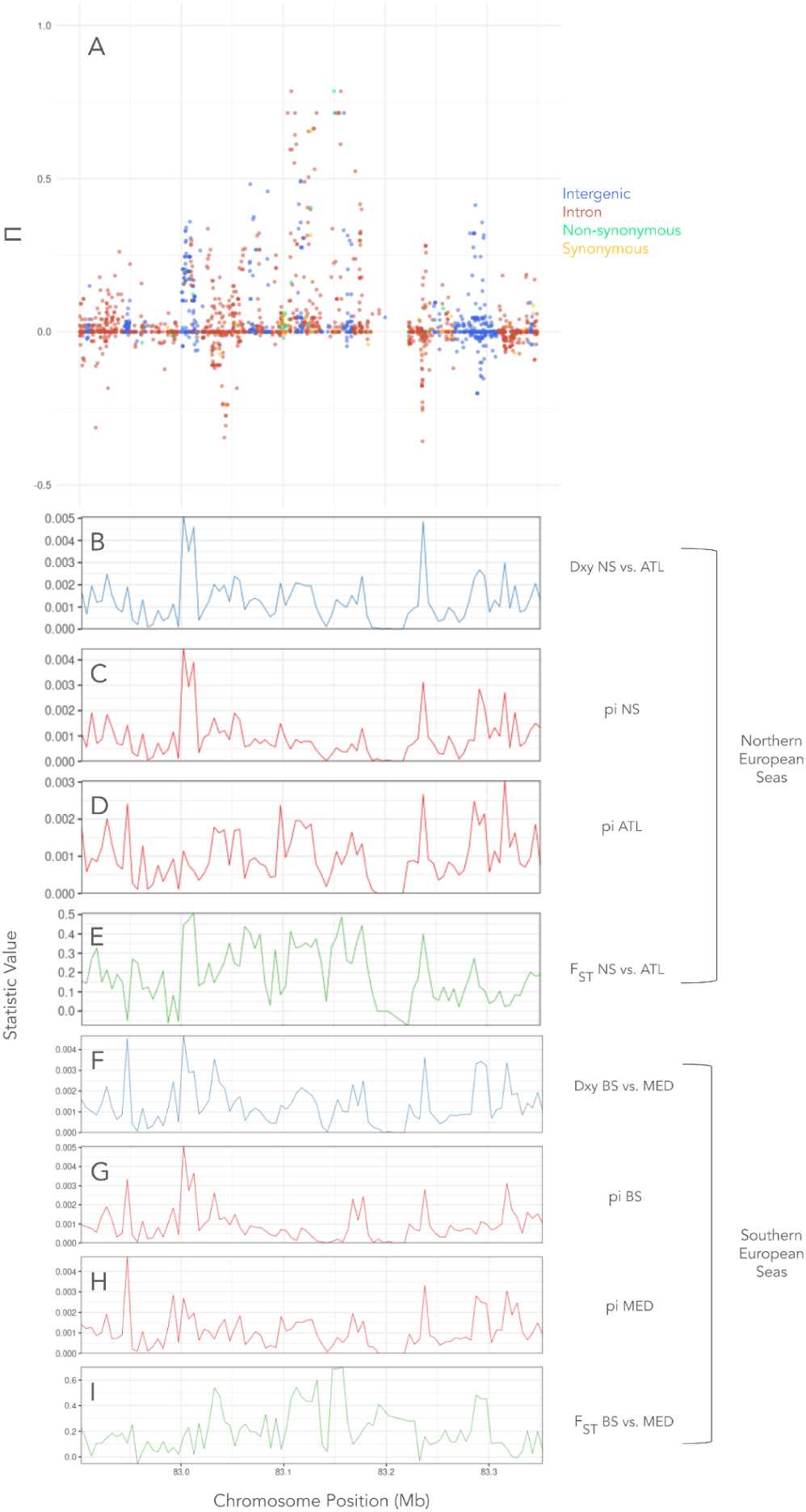
Island of parallelism located on chromosome 4 (Feature H, Island 7). **A** Plot of Π scan with chromosomal position along the x-axis and Π on the y-axis. Points refer to individual SNPs and colours refer to the category of SNP determined by SNPeff (intergenic, intronic, nonsynonymous or synonymous). **B** Dxy calculated in the NES region between the North Sea and Atlantic populations across the island region. **C** pi calculated for the North Sea population across the island region. **D** pi calculated for the Atlantic population across the island region. **E** Pairwise F_ST_ calculated in the NES region between the North Sea and Atlantic populations across the island region. **F** Dxy calculated in the SES region between the Black Sea and western Mediterranean populations across the island region. **G** pi calculated for the Black Sea population across the island region. **H** pi calculated for the western Mediterranean population across the island region. **I** Pairwise F_ST_ calculated in the SES region between the Black Sea and Mediterranean populations across the island region.

**Supplementary Figure 21.**
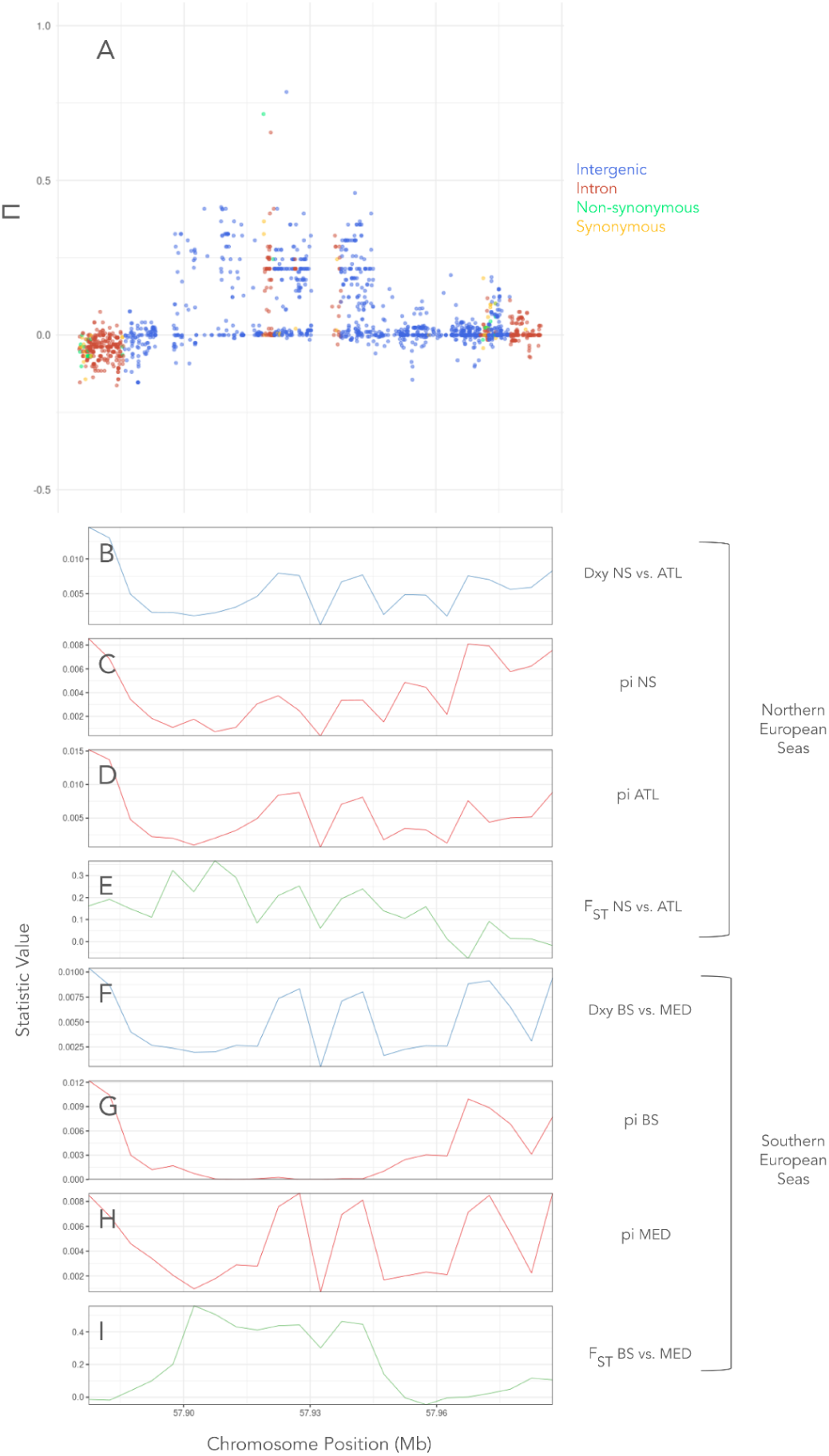
Island of parallelism located on chromosome 6 (Feature J, Island 9). **A** Plot of Π scan with chromosomal position along the x-axis and Π on the y-axis. Points refer to individual SNPs and colours refer to the category of SNP determined by SNPeff (intergenic, intronic, nonsynonymous or synonymous). **B** Dxy calculated in the NES region between the North Sea and Atlantic populations across the island region. **C** pi calculated for the North Sea population across the island region. **D** pi calculated for the Atlantic population across the island region. **E** Pairwise F_ST_ calculated in the NES region between the North Sea and Atlantic populations across the island region. **F** Dxy calculated in the SES region between the Black Sea and western Mediterranean populations across the island region. **G** pi calculated for the Black Sea population across the island region. **H** pi calculated for the western Mediterranean population across the island region. **I** Pairwise F_ST_ calculated in the SES region between the Black Sea and Mediterranean populations across the island region.

**Supplementary Figure 22.**
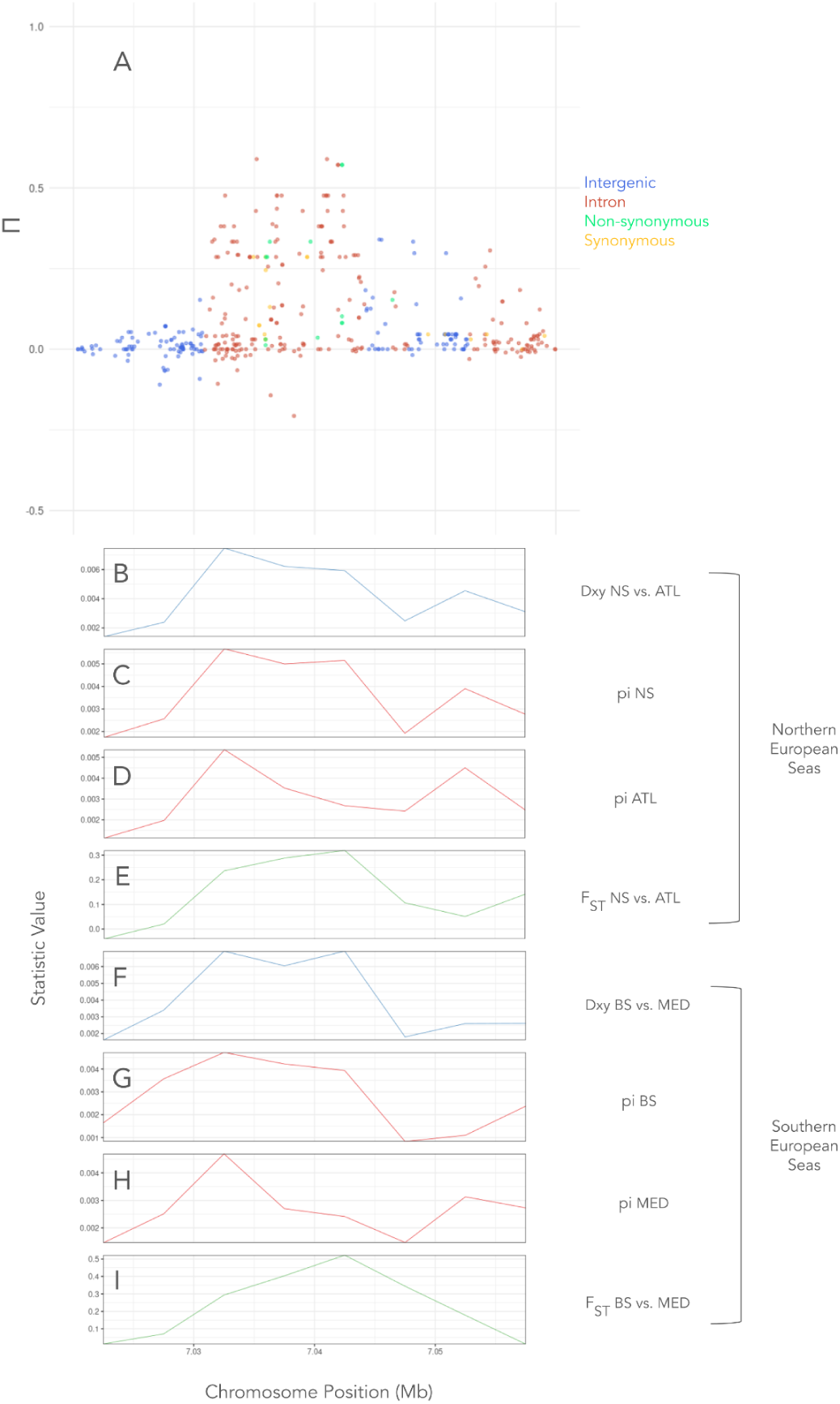
Island of parallelism located on chromosome 9 (Feature K, Island 10). **A** Plot of Π scan with chromosomal position along the x-axis and Π on the y-axis. Points refer to individual SNPs and colours refer to the category of SNP determined by SNPeff (intergenic, intronic, nonsynonymous or synonymous). **B** Dxy calculated in the NES region between the North Sea and Atlantic populations across the island region. **C** pi calculated for the North Sea population across the island region. **D** pi calculated for the Atlantic population across the island region. **E** Pairwise F_ST_ calculated in the NES region between the North Sea and Atlantic populations across the island region. **F** Dxy calculated in the SES region between the Black Sea and western Mediterranean populations across the island region. **G** pi calculated for the Black Sea population across the island region. **H** pi calculated for the western Mediterranean population across the island region. **I** Pairwise F_ST_ calculated in the SES region between the Black Sea and Mediterranean populations across the island region.

**Supplementary Figure 23.**
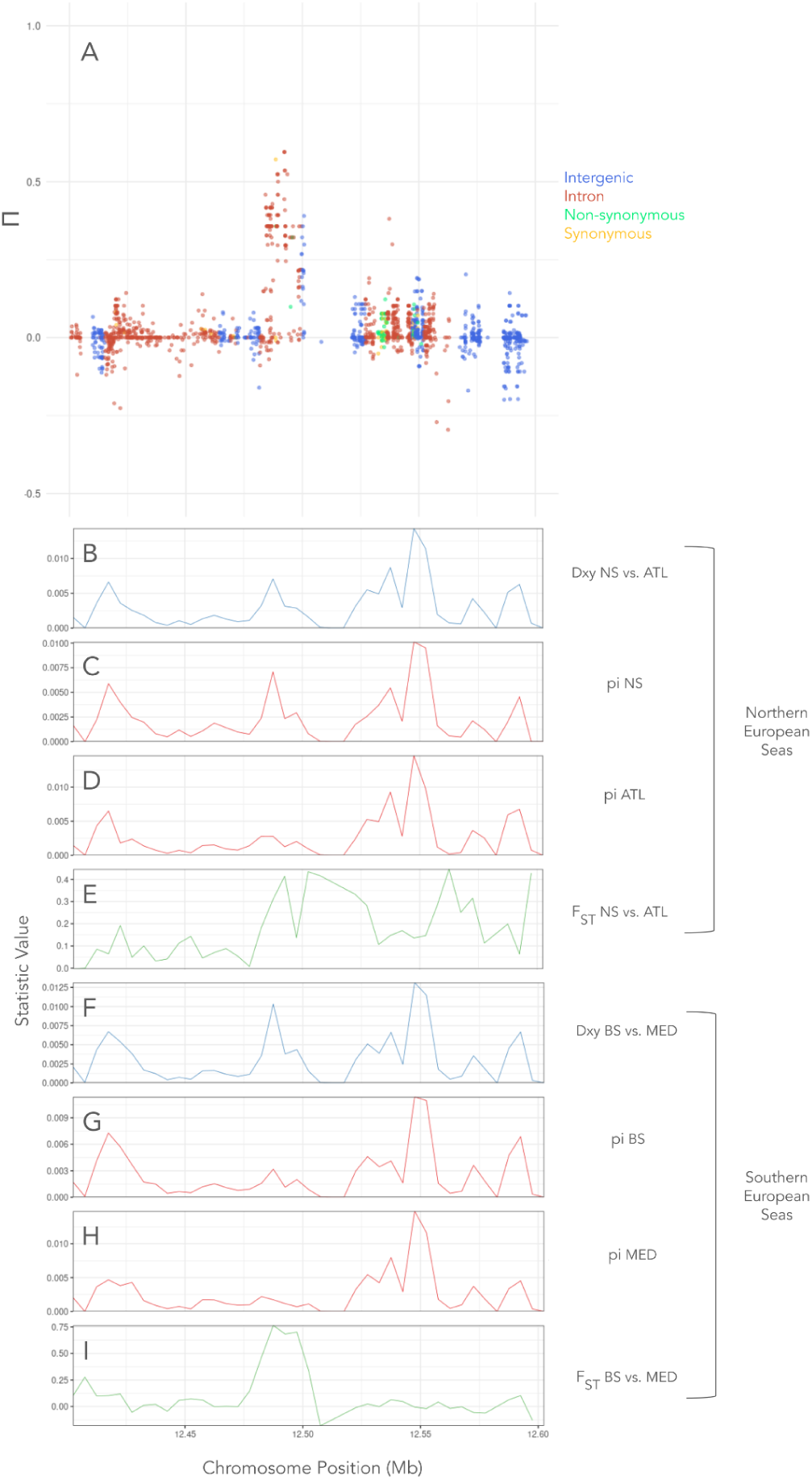
Island of parallelism located on chromosome 9 (Feature L, Island 11). **A** Plot of Π scan with chromosomal position along the x-axis and Π on the y-axis. Points refer to individual SNPs and colours refer to the category of SNP determined by SNPeff (intergenic, intronic, nonsynonymous or synonymous). **B** Dxy calculated in the NES region between the North Sea and Atlantic populations across the island region. **C** pi calculated for the North Sea population across the island region. **D** pi calculated for the Atlantic population across the island region. **E** Pairwise F_ST_ calculated in the NES region between the North Sea and Atlantic populations across the island region. **F** Dxy calculated in the SES region between the Black Sea and western Mediterranean populations across the island region. **G** pi calculated for the Black Sea population across the island region. **H** pi calculated for the western Mediterranean population across the island region. **I** Pairwise F_ST_ calculated in the SES region between the Black Sea and Mediterranean populations across the island region.

**Supplementary Figure 24.**
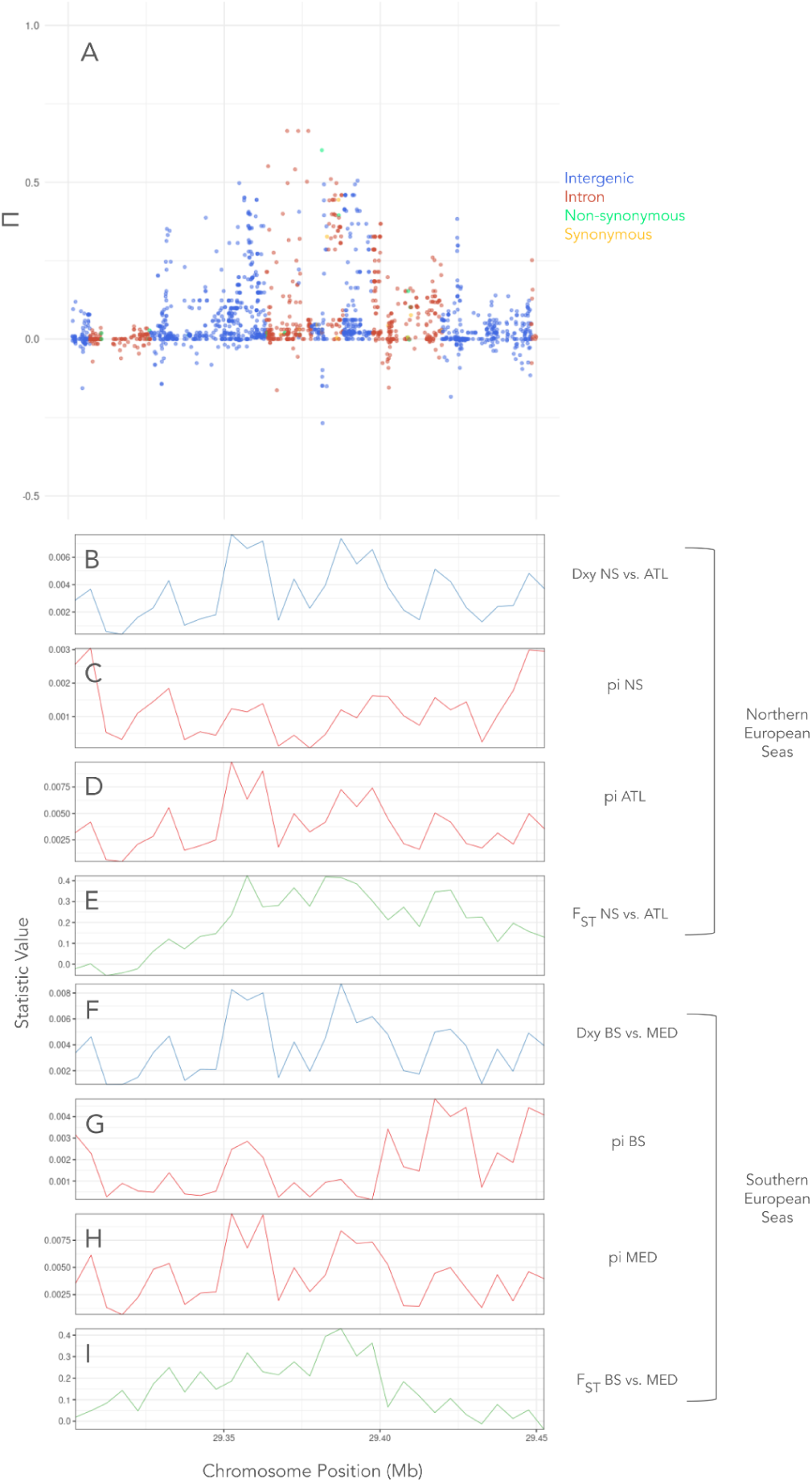
Island of parallelism located on chromosome 10 (Feature M, Island 12). **A** Plot of Π scan with chromosomal position along the x-axis and Π on the y-axis. Points refer to individual SNPs and colours refer to the category of SNP determined by SNPeff (intergenic, intronic, nonsynonymous or synonymous). **B** Dxy calculated in the NES region between the North Sea and Atlantic populations across the island region. **C** pi calculated for the North Sea population across the island region. **D** pi calculated for the Atlantic population across the island region. **E** Pairwise F_ST_ calculated in the NES region between the North Sea and Atlantic populations across the island region. **F** Dxy calculated in the SES region between the Black Sea and western Mediterranean populations across the island region. **G** pi calculated for the Black Sea population across the island region. **H** pi calculated for the western Mediterranean population across the island region. **I** Pairwise F_ST_ calculated in the SES region between the Black Sea and Mediterranean populations across the island region.

**Supplementary Figure 25.**
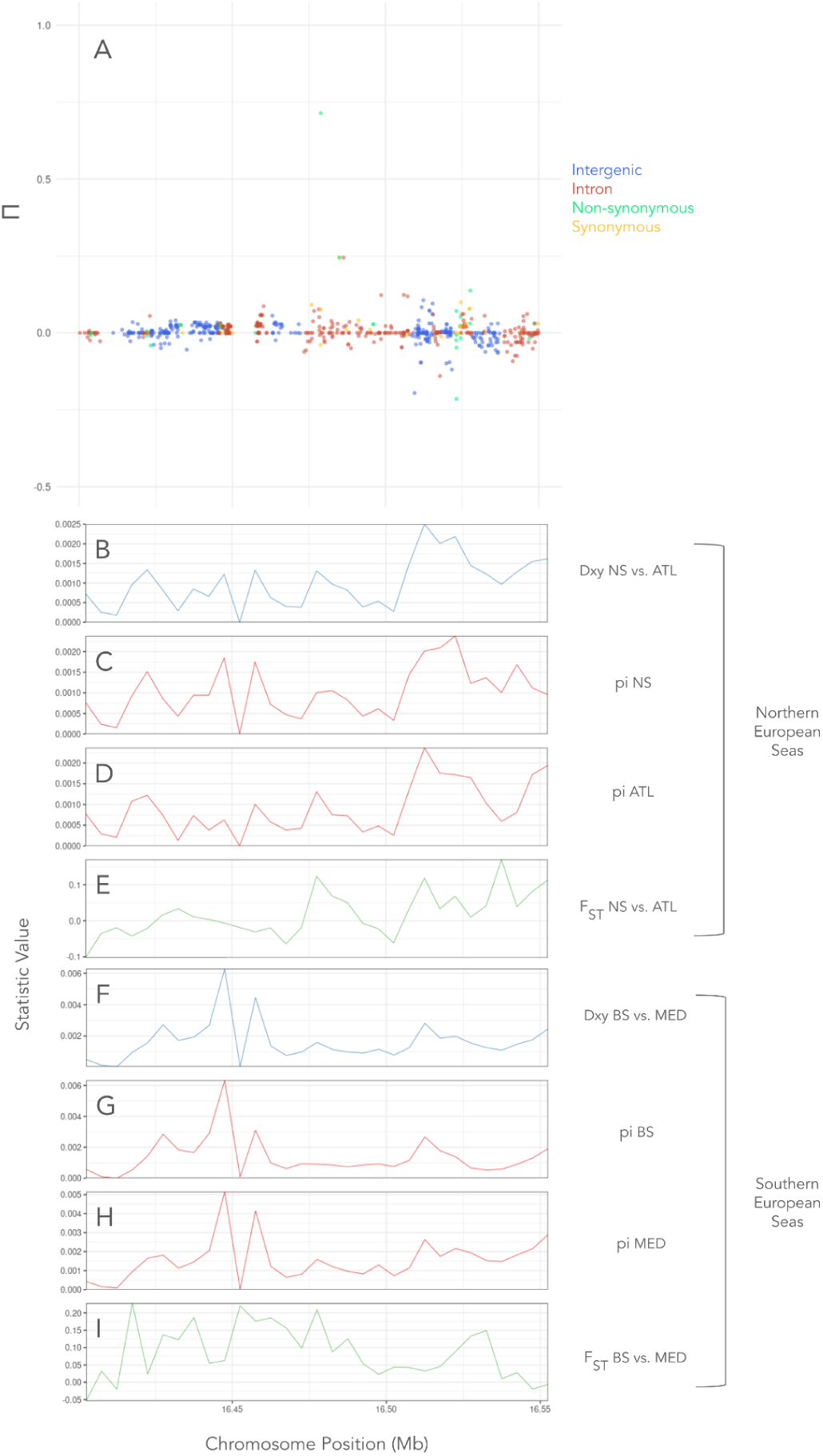
Single nonsynonymous SNP demonstrating an elevated Π score on chromosome 4 (Feature G). **A** Plot of Π scan with chromosomal position along the x-axis and Π on the y-axis. Points refer to individual SNPs and colours refer to the category of SNP determined by SNPeff (intergenic, intronic, nonsynonymous or synonymous). **B** Dxy calculated in the NES region between the North Sea and Atlantic populations across the island region. **C** pi calculated for the North Sea population across the island region. **D** pi calculated for the Atlantic population across the island region. **E** Pairwise F_ST_ calculated in the NES region between the North Sea and Atlantic populations across the island region. **F** Dxy calculated in the SES region between the Black Sea and western Mediterranean populations across the island region. **G** pi calculated for the Black Sea population across the island region. **H** pi calculated for the western Mediterranean population across the island region. **I** Pairwise F_ST_ calculated in the SES region between the Black Sea and Mediterranean populations across the island region

**Supplementary Figure 26.**
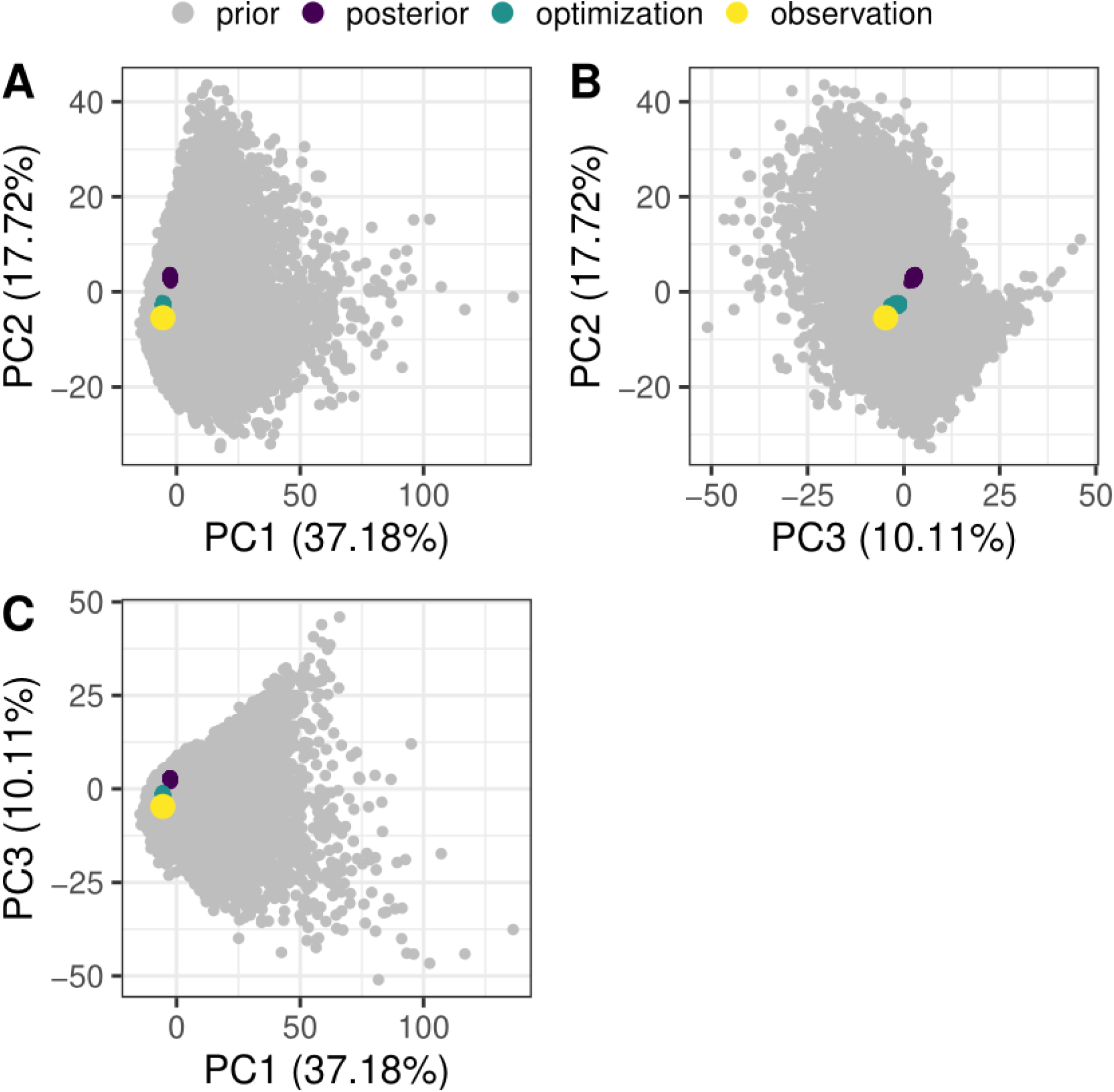
Global goodness-of-fit for the best model of the four-population inference (“topo3” i.e. the “genetic lineages” topology). The three first axes of a principal component analysis computed on the summary statistics of the Approximate Bayesian Computation implemented in DILS are shown. Each dot corresponds to a dataset: i) prior simulations in grey, ii) posterior simulations in purple, iii) optimized posterior simulations in green and iv) observed dataset in yellow. The proportion of variance explained by each axis is shown in brackets. 100,000 multilocus simulations were run.

**Supplementary Figure 27.**
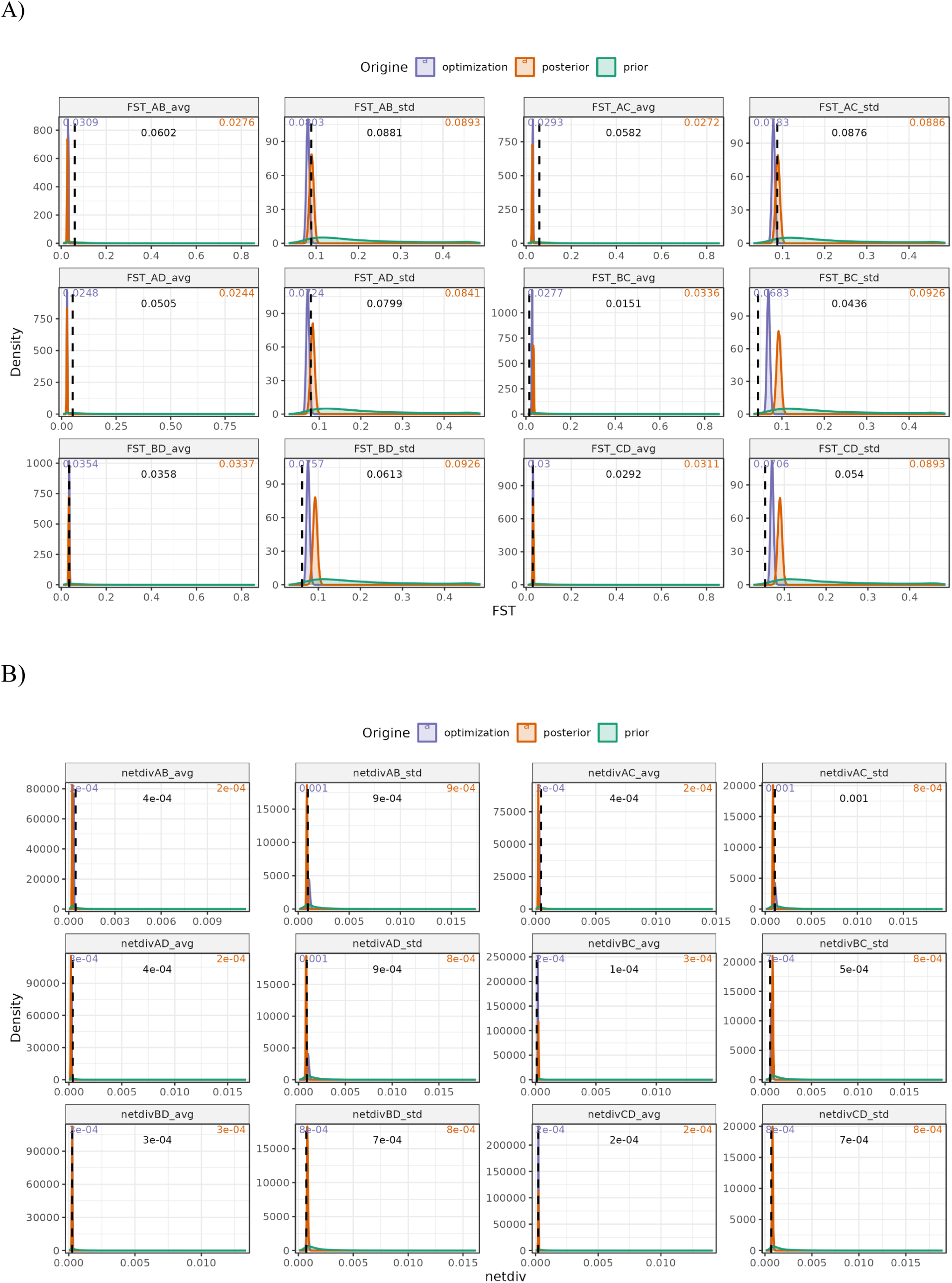
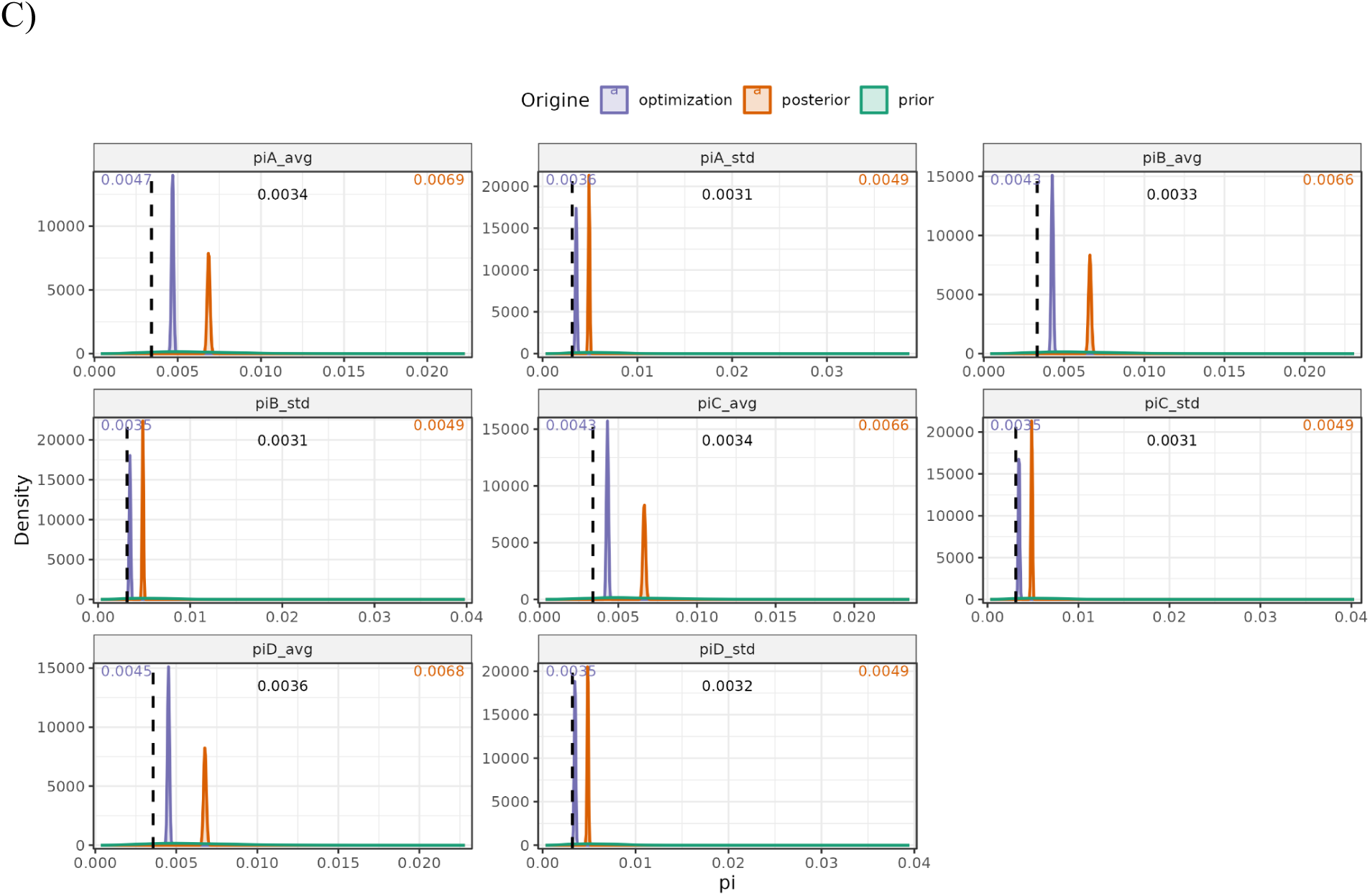
Per-statistic goodness-of-fit for the best model of the four-population inference (“topo3” i.e. the “genetic lineages” topology). Panels show: **A)** F_ST_, **B)** net divergence, *da* ( = d_XY_ - (π_X_+π_Y_)/2)) and **C)** intraspecific diversity, π. “avg” stands for the average of the statistic over loci and “std” for the standard deviation. The distribution of each summary statistic is shown for the prior simulations (green), the posterior simulations (red) and the optimized posterior simulations (blue). 4,000 simulations were randomly drawn for each category. The observed value of each statistic is depicted as a vertical dashed line (black). The median value for the posterior and optimized posterior are shown on the top, and they can be compared with the observed value. A is the Black Sea, B is the North Sea, C is the Atlantic and D is the western Mediterranean.

**Supplementary Figure 28.**
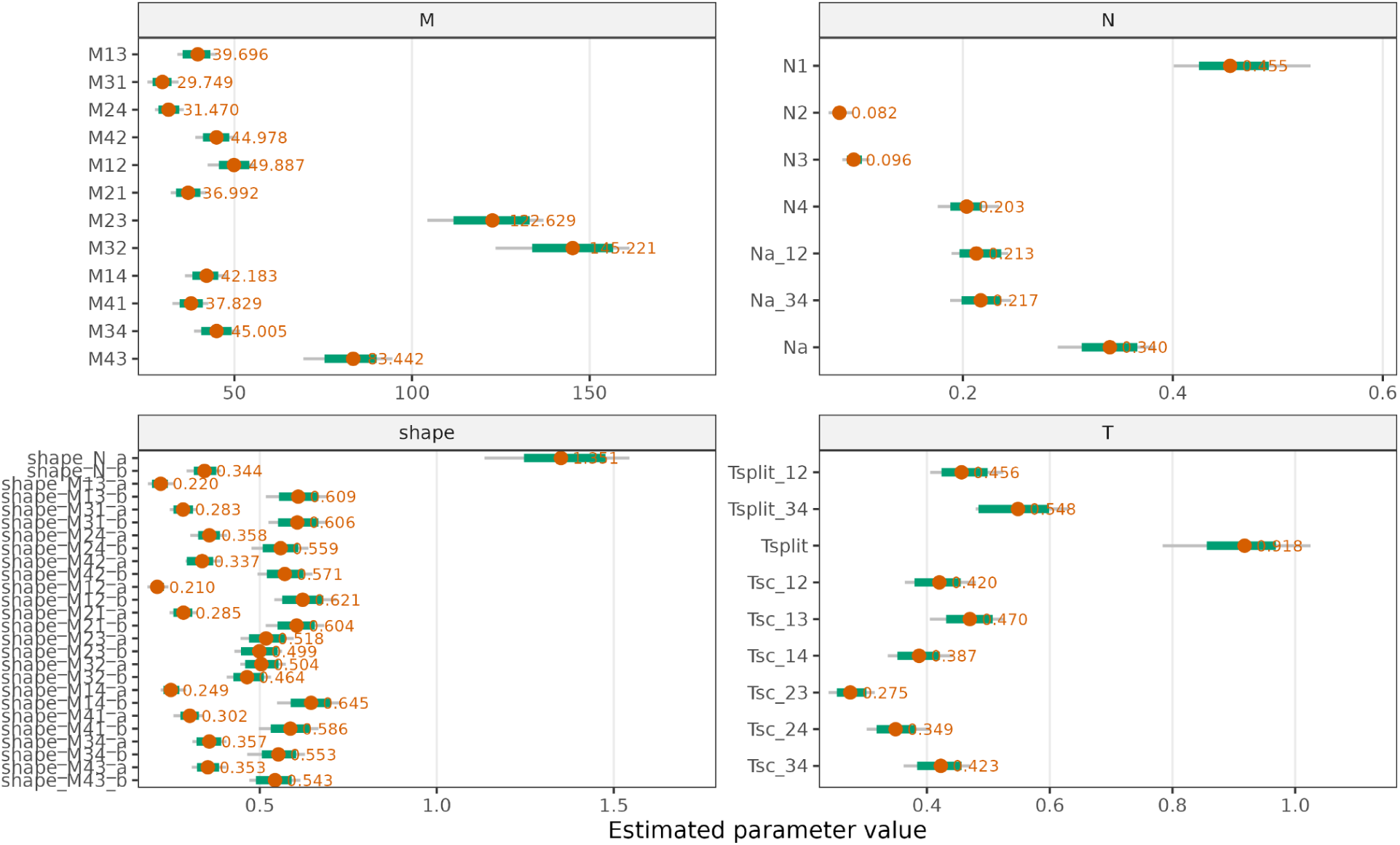
Estimated parameter values for the best model of the four-population inference (“topo3” i.e. the “genetic lineages” topology). Panels show: **A)** migration rates (in unit 4.*N.m*), **B)** effective population sizes (in unit *N*/*Nref* with *Nref* = 100,000), **C)** shapes of the beta distributions and **D)** times (in unit *T*/*Nref* with *Nref* = 100,000). The range from the minimum and maximum values of the posterior distribution is depicted as a grey segment. The range from quantile 2.5% and 97.5% of the optimized posterior distribution is depicted as a green segment. The median optimized posterior value for each parameter is depicted as a red dot and it is shown next to it. A is the Black Sea, B is the North Sea, C is the Atlantic and D is the western Mediterranean.

### Supplementary Statistics

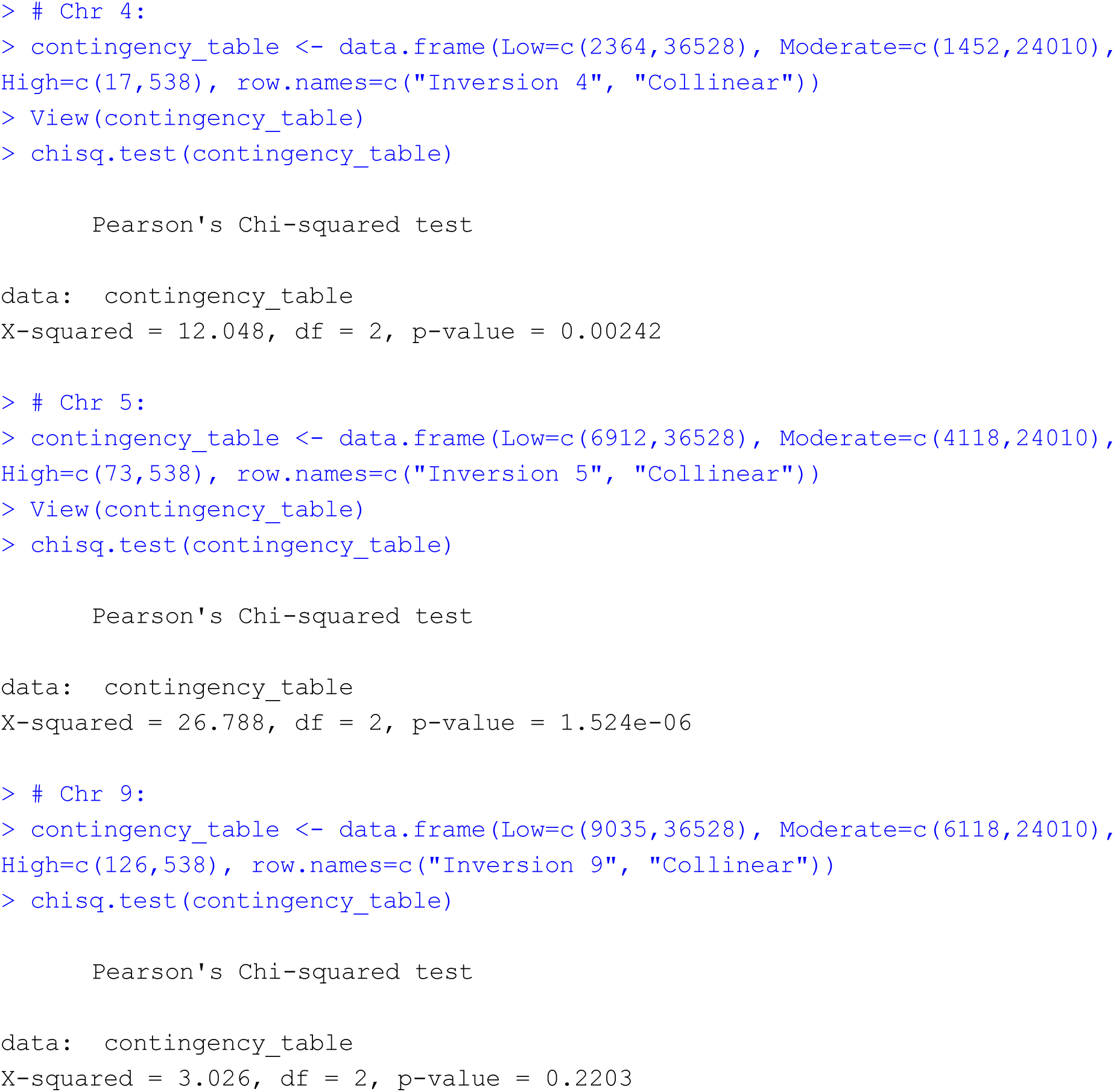

